# Individual Differences in Peripheral Hearing and Cognition Reveal Sentence Processing Differences in Healthy Older Adults

**DOI:** 10.1101/2020.06.16.118943

**Authors:** Ira Kurthen, Martin Meyer, Matthias Schlesewsky, Ina Bornkessel-Schlesewsky

## Abstract

When viewed cross-sectionally, aging seems to negatively affect speech comprehension. However, aging is a heterogeneous process, and variability among older adults is typically large. In this study, we investigated language comprehension as a function of individual differences in older adults. Specifically, we tested whether hearing thresholds, working memory, inhibition, and individual alpha frequency would predict event-related potential amplitudes in response to classic psycholinguistic manipulations at the sentence level. Twenty-nine healthy older adults (age range 61-76 years) listened to English sentences containing reduced relative clauses and object-relative clauses while their electroencephalogram was recorded. We found that hearing thresholds and working memory predicted P600 amplitudes early during reduced relative clause processing, while individual alpha frequency predicted P600 amplitudes at a later point in time. The results suggest that participants with better hearing and larger working memory capacity simultaneously activated both the preferred and the dispreferred interpretation of reduced relative clauses, while participants with worse hearing and smaller working memory capacity only activated the preferred interpretation. They also suggest that participants with a higher individual alpha frequency had a higher likelihood of successfully reanalysing the sentence towards the reduced relative clause reading than participants with a lower individual alpha frequency. By contrast, we found no relationship between object-relative clause processing and working memory or hearing thresholds. Taken together, the results support the view that older adults employ different strategies during auditory sentence processing dependent on their hearing and cognitive abilities and that there is no single ability that uniformly predicts sentence processing outcomes.

## Introduction

There is overwhelming evidence that aging negatively affects speech comprehension. The reasons are manifold: sensory degradation occurs as hearing loss develops and cognitive resources dwindle as brain structure and function ultimately succumb to age-related decline. However, as in all aging research, variability is large. In order to understand differential trajectories of speech comprehension in old age, key abilities that support speech comprehension in difficult listening situations need to be identified, which is one of the declared goals of Cognitive Hearing Science (Arlinger, Lunner, Lyxell, & Kathleen Pichora-Fuller, 2009). In this field of research, difficult listening situations have mostly been operationalized by introducing acoustic degradations to the speech signal, such as introducing noise or removing spectral content of the signal. However, a few studies have addressed the syntactic structure of the speech material itself, arguing that syntactic processing difficulty also constitutes an adverse listening condition (Wingfield, McCoy, Peelle, Tun, & Cox, 2006; Wingfield, Peelle, & Grossman, 2003).

Indeed, in cross-sectional research, and even in non-auditory studies, young and older adults usually differ in the quality of their language comprehension, with older adults exhibiting worse indicators of comprehension across a wide range of different measures (DeDe & Flax, 2016), such as slower reading times, difficulty in accessing infrequent words and in differentiating phonological neighbors, being slower in recognizing words, parsing sentences, and making more comprehension errors. All in all, there is ample cross-sectional evidence for between-group differences in language comprehension between younger and older adults. These mostly emerge not with simple language material, but when language material becomes more difficult to process (e.g. including double negation, comparatives, and doubly embedded relative clause sentences; Obler, Fein, Nicholas, & Albert, 1991, syntactically ambiguous garden-path sentences; Christianson, Williams, Zacks, & Ferreira, 2006; Kemper, Crow, & Kemtes, 2004, or non-prototypical animacy configurations; DeDe, 2015).

However, aging is a heterogeneous process (Lowsky, Olshansky, Bhattacharya, & Goldman, 2014) and chronological age can be understood “as a proxy for true mechanistic changes that influence functional capacity and adaptivity (including, but not limited to, cognition) across the lifetime” (S. W. S. MacDonald, DeCarlo, & Dixon, 2011, p. i59). Following this line of thought, there should be inter-individual variables more successful in explaining language comprehension than chronological age. These other variables will most likely co-vary with chronological age, and therefore at least partly bring about the group differences between younger and older adults. A study by Bornkessel-Schlesewsky et al. (2015) already showed that in a sample of healthy older adults, inter-individual variability outweighed effects of age. In another study, DeCaro, Peelle, Grossman, and Wingfield (2016) found that age did not significantly improve the prediction of comprehension accuracy when working memory capacity and hearing acuity were already present in the model. There are multiple candidate variables that may be related to successful language processing in older adults, including perceptual abilities which decline with age, such as hearing acuity (DeDe & Flax, 2016) and temporal processing abilities (Pichora-Fuller, 2003) in the case of spoken language. Other candidate mechanisms include cognitive abilities like processing speed (Salthouse, 1996), working memory (DeDe & Flax, 2016; Payne et al., 2014), inhibitory processes (Hasher & Zacks, 1988), and verbal fluency, which is thought to moderate the extent to which older adults use predictive processing (DeLong, Groppe, Urbach, & Kutas, 2012; Federmeier, McLennan, Ochoa, & Kutas, 2002).

All of these potential predictors have usually been investigated in separate studies and in single psycholinguistic paradigms. However, for the identification of key abilities that support speech comprehension in older adults, it is important to know whether there are overarching cognitive abilities that support speech comprehension in general, or whether different language processing challenges warrant involvement of different cognitive abilities. For our study, we thus chose two “classical” psycholinguistic paradigms. For an overview of the paradigms and the experimental conditions in our study, please see Table 1. First, we selected the paradigm employed by Osterhout and Holcomb (1992). In the following, we will refer to this as the reduced relative clause (RRC) paradigm because it involves a syntactically ambiguous relative clause construction. It is well suited for our study because English reduced relative clauses belong to the family of *garden-path sentences*, in which the preferred analysis of an ambiguous sentence region leads to an incorrect reading that needs to be corrected later. It has been shown that, in comparison to younger adults, older adults have a stronger tendency to adopt a “good-enough” interpretation of garden-path sentences (Christianson et al., 2006).

**Table 1.** Experimental Conditions.

| Paradigm | Condition | Example |
| --- | --- | --- |
| RRC | TVRR | "The broker persuaded to sell the stock was sent to jail." |
|  | TVDO | "The broker persuaded the investor to sell the stock." |
|  | IVWR | "The broker planned to sell the stock was sent to jail." |
|  | IVCO | "The broker planned to sell the stock." |
| ORC | ORAI | "The musician that the accident terrified angered the policeman a lot." |
|  | ORIA | "The accident that the musician witnessed angered the policeman a lot." |
|  | SRAI | "The musician that witnessed the accident angered the policeman a lot." |
|  | SRIA | "The accident that terrified the musician angered the policeman a lot." |

*Note.* This table shows the eight experimental conditions, clustered in the two paradigms, and lists an example sentence for each condition. RRC = reduced relative clause; ORC = object-relative clause; TVRR = transitive verb; reduced relative; TVDO = transitive verb, direct object; IVWR = intransitive verb, wrong; IVCO = intransitive verb, correct; ORAI = object-relative, animate - inanimate; ORIA = object-relative, inanimate - animate; SRAI = subject-relative, animate - inanimate; SRIA = subject-relative, inanimate - animate.

In a reduced relative clause (RRC) such as the TVRR example in Table 1, the ambiguous string *persuaded* – which is, in fact, a past participle – is initially interpreted as a past tense main clause verb (Bever, 1970). When *to* is subsequently encountered, *persuaded* must be reanalysed as a past participle within an RRC. A “good-enough” interpretation, by contrast, refers to cases in which the initial reading is not fully revised in spite of the conflicting evidence, i.e. in the case of our TVRR example, the assumption that the broker persuaded (someone) to do something would be (incorrectly) maintained. Crucially for present purposes, the RRC paradigm in Table 1 allows us to probe the extent to which participants reanalyse ambiguous RRC constructions. If a reanalysis has not taken place when the finite main clause verb (*was*) is encountered later in the sentence, it should render the sentence ungrammatical due to the slot of the main clause verb already having been filled by *persuaded*. This should engender an ungrammaticality-related response. A comparison between the TVRR and the IVWR sentences, which are indeed rendered ungrammatical at the position of *was*, can show the extent to which *persuaded* has been reinterpreted as a past participle. A second comparison, namely between TVRR vs. TVDO at the fourth word position (*to* vs *the*), shows the extent to which the initial disambiguation affects the well-formedness of the sentence.

For the second paradigm, we chose a variant of a manipulation that is commonly used in the current Cognitive Hearing Science literature. Most of the studies investigating relationships between language comprehension, syntactical processing, and aging have compared subject- and object-relative clause comprehension (Amichetti, White, & Wingfield, 2016; DeCaro et al., 2016; Wingfield et al., 2006). However, a considerable amount of evidence points to object-relative clauses not being more difficult to process than subject-relative clauses *per se*, but only when a certain animacy configuration is present, namely, when the subject of the main clause is animate and the subject of the object-relative clause is inanimate (DeDe, 2015; Traxler, Morris, & Seely, 2002; Weckerly & Kutas, 1999). Therefore, we based our second paradigm on Traxler et al.’s (2002) object relative clause design with an animacy manipulation. We further refer to it as the object relative clause (ORC) paradigm. It allows us to test predictive processes during actor computation. Taking the example from Table 1, ORAI sentences have an animate subject in the main clause and an inanimate subject in the object-relative clause, while the ORIA sentences have an inanimate subject in the main clause and and animate subject in the object-relative clause. Taking animacy as a prominence feature which strongly guides thematic role assignment (Bornkessel-Schlesewsky & Schlesewsky, 2009), one would assume that the animate object-relative clause subject (e.g., the *musician*) in the ORIA sentences is a prototypical instantiation of the actor role (being the agent that does something to the inanimate main clause subject, e.g., the *accident*). By contrast, the inanimate object-relative clause subject (e.g., the *accident*) in the ORAI sentences does not correspond to a prototypical actor. If participants make use of the previous information (animacy of the main clause subject and the presence of an object-relative clause), they should therefore predict an animate object-relative clause subject in both the ORIA and the ORAI sentences. When that prediction is not fulfilled in the ORAI sentences, we should observe a response related to the prediction error (Bornkessel-Schlesewsky & Schlesewsky, 2019).

Both our paradigms have reliably elicited inter-individual processing differences, as revealed by different indicators of processing difficulty. Kemper et al. (2004) found differences between high- and low-working-memory-span individuals in RRC processing, but no differences between age groups. However, Yoo and Dickey (2017) found a difference between younger and older adults during processing of reduced relative clauses, but neither working memory nor inhibition predicted the prolonged reading times. With regard to the ORC paradigm, Traxler, Williams, Blozis, and Morris (2005) showed that high-span subjects benefited more from animacy cues than low-span subjects. In an ERP study by Weckerly and Kutas (1999), there was only an N400 effect in response to inanimate object-relative clause subjects as compared to animate object-relative clause subjects in high comprehenders (i.e. participants who scored higher than 75% on the comprehension task for ORCs), but not in low comprehenders.

To measure processing difficulties, previous studies employed methods of either response accuracy (comprehension questions, Amichetti et al., 2016; DeCaro et al., 2016; Wingfield et al., 2006), or reading/listening times (eye-tracking; Traxler et al., 2002 and self-paced listening DeDe, 2015). Because we aimed for auditory presentation of our stimuli (thereby excluding reading measures), and because the RRC paradigm allowed for probing sentential processing at multiple points in time (thereby excluding end-of-sentence behavioral comprehension measures), we chose event-related potentials (ERPs) as our online sentence processing markers of choice. Both paradigms have previously been examined using ERPs. In the RRC paradigm, Osterhout and Holcomb (1992, 1993) observed P600 effects for both the reanalysis- and ungrammaticality related comparisons (i.e. for TVRR vs. TVDO and TVRR vs. IVWR, respectively). For the ORC paradigm, the study by Weckerly and Kutas (1999) revealed an N400 effect for good comprehenders as noted above (cf. also Frisch & Schlesewsky, 2001).

An additional reason for using ERPs is that they have previously exhibited modulation by cognitive ability (Bornkessel, Fiebach, & Friederici, 2004; Kim, Oines, & Miyake, 2018; Nakano, Saron, & Swaab, 2010). Friederici, Steinhauer, Mecklinger, and Meyer (1998) showed a P600 at disambiguating positions in garden-path sentences for readers with a high working memory span, but not for readers with a low working memory span. Weckerly and Kutas (1999) observed an N400 at an inanimate object-relative clause subjects only for good comprehenders, and DeLong et al. (2012) reported a frontal positivity in response to constraint violations only in older adults with high verbal fluency.

### Predictors

We selected several inter-individual predictors for ERP amplitude between the conditions to be compared. First, we chose peripheral hearing loss as measured by hearing thresholds. Hearing loss is highly prevalent in older adults – approximately 20% at age 60 and 50% at age 70 (Bisgaard & Ruf, 2017; Goman & Lin, 2016; Mick et al., 2019) – and hearing thresholds have been shown to influence many behavioral results in previous studies (DeCaro et al., 2016; DeDe & Flax, 2016; Wingfield et al., 2006), even in young adults (Ayasse, Penn, & Wingfield, 2019).

Second, we chose working memory capacity, which has featured prominently in many studies on inter-individual differences in language comprehension (e.g. Bornkessel, Fiebach, & Friederici, 2004; Friederici et al., 1998; Nakano et al., 2010)

A third predictor was individual alpha frequency (IAF), the peak frequency within the EEG alpha band (approximately 8 to 13 Hz), which is known to vary between individuals (Klimesch, 1999). IAF has been shown to correlate with cognitive ability (Angelakis, Lubar, & Stathopoulou, 2004; Angelakis, Lubar, Stathopoulou, & Kounios, 2004; Grandy, Werkle-Bergner, Chicherio, Lövdén, et al., 2013; Klimesch, Schimke, & Pfurtscheller, 1993; Mundy-Castle, 1958), and while it tends to decrease with age cross-sectionally, it is a stable neurophysiological trait (Grandy, Werkle-Bergner, Chicherio, Schmiedek, et al., 2013). We chose to investigate IAF because it is a rather general marker for cognitive ability, also reflected in its substantial correlation with the *g* factor of general intelligence (Grandy, Werkle-Bergner, Chicherio, Lövdén, et al., 2013) and because it has already been associated with individual differences in language processing (Bornkessel, Fiebach, & Friederici, 2004) as well as modulations of the late positivity in older adults (Bornkessel-Schlesewsky et al., 2015).

Lastly, we chose to investigate inhibition as a predictor for ERP amplitude. According to Hasher and Zacks (1988), inhibitory processes can serve as gatekeepers for working memory during language processing. These authors further proposed that the reduced efficiency of these processes in older adults may underlie the decline of cognitive abilities – including certain aspects of language processing – with increasing age. Inhibition, or executive control, has also been put forward as a mechanism to suppress an initial, preferred interpretation in favor of an alternative interpretation which better fits the sentential information (see Novick, Trueswell, & Thompson-Schill, 2010, for a review). Furthermore, Vuong and Martin (2014) showed that verbal Stroop performance predicted correct garden-path revisions (although see Engelhardt, Nigg, & Ferreira, 2017, for a study where intelligence is a better predictor of garden-path comprehension accuracy than inhibition).

### Study Design and Hypotheses

We aimed to investigate ERP amplitude in response to two classical psycholinguistic manipulations as a function of inter-individual differences in hearing and cognitive ability. If present, such a modulation would indicate different processing strategies, which in turn might explain the often-observed language comprehension benefits for older adults with better hearing and cognitive ability.

For the RRC paradigm, we first compared ERP amplitude in conditions TVRR and TVDO at the fourth position (…persuaded *to* vs. …persuaded *the*). The amplitude of the P600 between the infinitival marker *to* in the TVRR sentences and the definite article *the* in the TVDO sentences indicates how strongly the interpretation of *persuaded* had been biased towards a past tense main clause verb.

Additionally, we repeated the analysis described in Osterhout and Holcomb (1992), comparing ERP amplitude in conditions IVWR and TVRR at the eighth position (…planned to… *was* vs. …persuaded to… *was*, following the examples from Table 1). The auxiliary verbs at position eight in conditions IVWR und TVRR either rendered the sentence ungrammatical (IVWR) or continued the main clause (TVRR). Therefore, a comparison between these auxiliary verbs would reveal whether a successful reanalysis had previously taken place in condition TVRR. If it had, a finite main clause verb (such as an auxiliary) should be expected and therefore, one would expect a P600 for IVWR vs. TVRR to mark the ungrammaticality of the former. On the other hand, if a reanalysis had not taken place in the TVRR condition and *persuaded* was rather interpreted as a past tense main clause verb, both IVWR and TVRR should engender an ungrammaticality-related response at the position of the auxiliary and there should be no difference between the two conditions. Thus, the presence of a response for IVWR vs. TVRR at this position can be viewed as a marker of successful reanalysis towards a reduced relative clause earlier on in the sentence.

For the ORC paradigm, we followed the analysis by Weckerly and Kutas (1999). Specifically, we compared ERP amplitude in conditions ORAI and ORIA at the fifth position (The musician that the *accident*… vs. The accident that the *musician*…). Weckerly and Kutas (1999) showed that an N400 was elicited for an inanimate relative clause subject compared to an animate relative clause subject, arguably resulting from the additional processing costs of assigning an actor role to an inanimate subject (Bornkessel & Schlesewsky, 2006). Interestingly, this effect was only present in good comprehenders. Because comprehension accuracy of object-relative clauses has been shown to be associated with hearing loss and working memory capacity in older adults (DeCaro et al., 2016; Wingfield et al., 2006), it appears reasonable to assume that an N400 elicited after inanimate ORC subjects in comparison with animate ORC subjects may also be associated with these inter-individual variables.

As noted above, modulation of ERP amplitudes between different conditions by several inter-individual variables was of particular interest for this study. For the comparison between TVRR and TVDO sentences at the fourth position, we expected participants with (potentially) fewer resources available to exhibit higher P600s, meaning participants with higher hearing thresholds, lower working memory capacity, and lower IAF. We also expected participants with higher inhibition to exhibit higher P600s, because they would be more prone to suppress the second meaning of the ambiguous string *persuaded*, and would therefore be more surprised when encountering the unexpected continuation in the TVRR sentences.

For the second comparison, IVWR vs. TVRR, we expected participants with fewer resources to exhibit smaller P600s, because they might settle for a good-enough interpretation of the reduced relative clause and therefore show the same ungrammaticality response for the TVRR sentences at the eighth position as for the IVWR sentences. We assume that this again holds for participants with higher hearing thresholds, lower working memory capacity, and lower IAF.

Note that previous studies on older adults’ ORC processing compared comprehension in subject-vs. object-relative clauses (e.g. Amichetti et al., 2016; DeCaro et al., 2016; Wingfield et al., 2006, 2003). The stimuli in these studies involved animate subjects for both the main and the relative clause. This arrangement results in competition for the actor role, which appears to be the feature which renders ORCs difficult to process (DeDe, 2015; Traxler et al., 2002; Weckerly & Kutas, 1999). We decided to follow Weckerly and Kutas (1999) in comparing ORCs with an animacy manipulation. This conveniently solves the problem of otherwise having to compare noun phrases at different sentential positions. Although hearing thresholds and working memory have been found to predict ORC comprehension only when compared to SRC processing, we nevertheless hypothesize that they might also predict the sensitivity to animacy as a cue for sentence processing as reflected in the N400. We therefore hypothesized that lower hearing thresholds and higher working memory capacity would result in a larger N400 effect between ORAI and ORIA sentences.

It is possible that we might observe a modulation of ERP amplitudes by hearing thresholds and cognitive ability on the basis of altered auditory processing in general and not because of different processing strategies for linguistic material. If that were the case, we should also observe a modulation of earlier “pre-linguistic” auditory ERP amplitudes. To test for this association, we added a mismatch negativity (MMN; Näätänen, Pakarinen, Rinne, & Takegata, 2004) paradigm to the study. If hearing and cognitive abilities predict both MMN and N400/P600 amplitudes, this would suggest that hearing and/or cognitive ability affects auditory processing in general, and that this effect is not restricted to auditory sentence processing. If hearing and cognitive abilities only predict N400/P600 amplitudes, but not MMN amplitude, this would strengthen the argument that effects of hearing and cognition mainly come into play at later processing stages of sentence comprehension.

## Materials and Methods

All data and code associated with the study will be made available on the Open Science Framework upon acceptance of the paper.

### Participants

The sample consisted of 29 older adults (mean age = 66.14 yrs, sd = 3.70 yrs, range 61 - 76). Three more older adults participated in the study but were excluded due to excessive EEG artifacts. All participants were right-handed and reported no psychiatric or neurological disorders. Their native language was English and they had not learned another language before their seventh year of age. They did not wear a hearing aid and they reported not to have tinnitus. They also were not colorblind. Their peripheral hearing thresholds did not exceed 30 dB in the frequencies 0.5, 1, 2, and 4 kHz. They passed a screening session in which the exclusion criteria were tested via questionnaires. In order to exclude participants with Mild Cognitive Impairment, they were administered the Montreal Cognitive Assessment (MoCA; Nasreddine et al., 2005) and were invited to further participate in the study when they scored 26 points or more.

The study protocol was approved by the Human Research Ethics Committee of the University of South Australia. All participants gave written informed consent in accordance with the Declaration of Helsinki.

### Study Process

The study consisted of one session which took about three hours to complete. After participants passed the screening (20 minutes), they completed four cognitive tasks: Two inhibition tasks (Stroop Task; Golden, 1976, and Eriksen-Flanker Task; Eriksen & Eriksen, 1974) and two working memory tasks (Reading (Sentence) Span (RS) and Operation Span (OS); modelled after Lewandowsky, Oberauer, Yang, & Ecker, 2010). Because the two working memory tasks were rather similar, they were administered in counterbalanced order. After that, participants took part in an EEG experiment which took about 45 minutes to complete. At the beginning and end of the EEG session, resting state EEG was measured (two minutes with eyes open, two minutes with eyes closed). After the first resting state session, a short MMN paradigm was administered, which took about three and a half minutes. After that, the main EEG task started. In this main task, participants listened to acoustically presented sentences and rated their acceptability. Participants received a 50 AUD Coles & Myer gift card for their participation.

### Hearing Thresholds

The computer-based hearing tests were administered via a custom MATLAB software built upon the MAP auditory toolbox (Meddis et al., 2013). We measured absolute pure-tone hearing thresholds (pure-tone audiometry; PTA) by means of a probe-detection paradigm. Participants were played either one or two sine wave tones for 250 ms each and indicated whether they had heard two, one, or no sounds. The probe was always 10 dB SPL lower than the cue and the loudness of cue and probe was varied by means of an adaptive procedure. Participants practiced the task with sine wave tones of 1 kHz and were subsequently tested on frequencies 0.25, 0.5, 1, 2, 4, 6, and 8 kHz. The average hearing threshold for each participant was calculated by averaging the thresholds for 0.5, 1, 2, and 4 kHz. The measurement procedure and the stimuli have been described in detail elsewhere (Giroud et al., 2018; Lecluyse & Meddis, 2009; Lecluyse, Tan, McFerran, & Meddis, 2013).

### Working Memory Tasks

The two working memory task were a RS and an OS task. They were programmed in PsychoPy2 (Version 1.90.2) and modelled after Lewandowsky et al. (2010). Sentences were very easy to classify as “correct” or “false”, but not at first glance (example: “The earth is larger than the sun.”). The difficulty in this task was kept low because this improved the correspondence between the RS measure and a latent measure of working memory capacity (Lewandowsky et al., 2010). The equations in the OS task were also very easy (only addition and subtraction with one- or two-digit numbers; no subtraction with borrowing).

### Inhibition Tasks

The Flanker task was also programmed in PsychoPy2 (Version 1.90.2). Participants were presented with 30 trials showing five arrows all pointing in the same direction, left or right (congruent), or the middle arrow pointing into the opposite direction than the other four (incongruent), or only one arrow pointing either left or right, with four squares around it (neutral). The Flanker inhibition score was calculated by subtracting the mean reaction time to the incongruent stimuli from the mean reaction time to the congruent stimuli.

We used a pen-and-paper version of the Stroop task to obtain the Stroop interference score. Participants had 45 seconds each to work through three sheets. Sheet one consisted of the words RED, BLUE, and GREEN printed in black, and participants had to read those out aloud as fast as possible, which yielded score W (number of words read). Sheet two consisted of the characters “XXXX” printed in either red, blue, or green. Participants had to name the colors of the printed characters as fast as possible, which yielded the score C (number of colors named). Sheet three consisted of the words RED, BLUE, and GREEN printed in either red, blue, or green, but never in the color they represented. Pseudo-randomization of the order of words and colors was carried out via Mix (van Casteren & Davis, 2006). Participants again had to name the colors of the printed characters as fast as possible, which yielded the score CW (number of colors named). An interference score IG was calculated with the formulae Pcw=(W*C)/(W+C) and IG=CW-Pcw (Golden & Freshwater, 1978), which is the most commonly used Stroop interference score (Scarpina & Tagini, 2017).

### Sentence Stimuli

In total, the main EEG experiment used 600 sentence stimuli. Stimuli were recorded by a male native speaker of Australian English (mean F0 = 98.44 Hz, sd = 5.17 Hz).

Please see Table 1 for an overview of the experimental conditions. Sentence materials for the RRC paradigm were taken from Osterhout and Holcomb (1992), Experiment 2. We adopted their conditions 1 (short intransitive verb sentences; IVCO), 3 (long, grammatically incorrect intransitive verb sentence; IVWR), and 4 (reduced relative clause/long intransitive verb sentence; TVRR). However, instead of condition 2 in the original experiment, we chose to present sentences with a transitive verb and its direct object (condition TVDO), because, in contrast to condition 2 of Osterhout and Holcomb (1992), this resulted in a grammatically correct and linguistically highly acceptable condition. This replacement was chosen in order to achieve an overall higher proportion of grammatically correct sentences in the whole experiment.

Sentence materials for the ORC paradigm were taken from Traxler et al. (2002), Experiment 3. We exactly adopted their four conditions, two of which contained subject-relative (SR) clauses and two of which contained object-relative (OR) clauses. These sub-divided conditions further differed with regard to the animacy of their main clause and relative clause subjects. In the SRAI and the ORAI conditions, the main clause subject was animate and the relative clause subject was inanimate, while in the SRIA and the ORIA conditions, the main clause subject was inanimate and the relative clause subject was animate. As Traxler et al.’s original experiment only contained 28 sentences per condition, we added two more sets of sentences. Because both paradigms contained sentence materials that were not part of the original studies, all sentences for both paradigms can be found in Supplementary Tables S1-S8.

Participants were presented with 240 sentences, subdivided into eight blocks of 30 sentences each. Each participant was presented with all of the sentences in the ORC paradigm (30 per condition). Because there were 120 stimuli available for each condition of the RRC paradigm (480 in total), we subdivided these into four lists of 120 sentences (30 sentences per condition) using a Latin Square design. List presentation was counterbalanced across participants, with each participant presented with one of the four lists, interspersed with the ORC sentences. Pseudo-randomization of trials was carried out via Mix (van Casteren & Davis, 2006), with the constraint that sentences from one condition must not be played directly after one another.

### Test for Differences in Speech Parameters Between Conditions

In order to test for differences in speech parameters at the word positions of interest between the conditions, we extracted mean F0 (pitch), duration, and mean intensity via a custom-written Praat (Boersma & van Heuven, 2001) script and compared them using Welch two-sample t-tests. Table 2 shows the mean values per condition for each word positions of interest as well as t-test results. Speech parameters at the word positions of interest did not differ significantly between conditions, there was only a significant difference in intensity at word position 4 between the TVRR and TVDO conditions (*to* vs. *the*). However, that difference was just slightly above 1 dB (−1.25 dB) and, due to the very short duration of the words, most likely not perceivable by our participants. Even if it had been perceivable, this should not discredit our results, because we did not aim for complete indistinctiveness of the conditions, but we rather were interested in how participants would differentially utilize these cues for comprehension.

**Table 2.** Pitch, duration, and intensity comparison of critical word positions.

|  |  | pitch [Hz] |  |  |  | duration [s] |  |  |  | intensity [dB SPL] |  |  |  |
| --- | --- | --- | --- | --- | --- | --- | --- | --- | --- | --- | --- | --- | --- |
|  | w.pos | m | t | df | p | m | t | df | p | m | t | df | p |
| TVRR | 4 | 92.43 | -0.917 | 128.07 | 0.3608 | 0.11 | 1.022 | 234.37 | 0.308 | 65.71 | -5.922 | 234.78 | <.001 |
| TVDO | 4 | 90.78 |  |  |  | 0.11 |  |  |  | 64.46 |  |  |  |
| IVWR | 8 | 84.31 | -1.722 | 227.33 | 0.087 | 0.19 | -0.219 | 229.27 | 0.8273 | 64.73 | 0.221 | 231.96 | 0.8251 |
| TVRR | 8 | 85.16 |  |  |  | 0.19 |  |  |  | 64.69 |  |  |  |
| ORAI | 5 | 102.14 | 1.335 | 57.784 | 0.1871 | 0.39 | -1.050 | 49.361 | 0.299 | 69.17 | -0.595 | 58 | 0.5545 |
| ORIA | 5 | 99.76 |  |  |  | 0.41 |  |  |  | 69.42 |  |  |  |

*Note.* This table shows the mean values per condition for pitch, duration, and intensity of each word positions of interest as well as the results of the Welch two-sample t-tests used to compare them.

### Procedure

At the beginning of each trial, an asterisk was presented on the screen for 500 ms, after which auditory presentation of the sentence commenced. The asterisk continued to be displayed throughout the auditory presentation of the sentence. After a gap of 500 ms after the sentence had ended, participants were prompted to rate the acceptability of the sentence on a scale from 1 (“The sentence was not a good English sentence at all”) to 4 (“The sentence was a very good English sentence”). Participants had 4 seconds to respond to the question by means of a keyboard button press. If they did not respond within this time frame, the next trial began. The inter-trial interval was 1500 ms long. Between blocks, participants took self-paced breaks.

Before testing started, participants were given a set of eight items as a practice block. These eight items contained two sentences per condition from a subset of the RRC paradigm which was not presented to the participant later. During the practice block, participants’ response behavior was monitored and the task was explained again if necessary (e.g. if the participant never responded to the practice items or if the participant always responded with the same button). After the practice session, participants were encouraged to attenuate or amplify the stimuli in order to obtain a comfortable sound level.

### EEG Recording and Preprocessing

Participants’ EEG was recorded continuously from 59 Ag/AgCl electrodes (ActiCAP, Brain Products) with a BrainVision actiCHamp Active Electrodes amplifier system (Brain Products GmbH, Gilching, Germany) at 500 Hz. The electrodes were spaced according to the 10–20 system, with FT9, FT10, Fp1, Fp2, and TP9 missing because these electrodes were used for other purposes (electrooculogram (EOG) and reference). For monitoring eye movements and blinks, the horizontal and vertical EOG was recorded with supra- and infraorbital electrodes on the left eye and two electrodes placed next to the external canthi of the left and right eyes. Impedances were reduced below 25 kOhm. A forehead ground (Fz) and a left mastoid reference (TP9) were used. Data were analyzed in MATLAB Release 2016b (The MathWorks, Inc., Natick, Massachusetts, United States) using the FieldTrip Toolbox (Version 20190419; Oostenveld, Fries, Maris, & Schoffelen, 2011). For pre-processing, data were first visually screened for noisy channels. Afterwards, trials were defined, starting 2000 ms before sentence onset and ending 500 ms after the end of the sentence. After that, an automatic artifact rejection (AAR) procedure was employed. For AAR, data were first filtered between 0.1 and 10 Hz and z-values were computed for each trial. Trials that exhibited a z-value higher than a certain threshold (mostly 60, but this had to be adjusted for some participants) were marked as bad trials. In parallel, data were filtered between 110 and 140 Hz and again, z-values were computed for each trial. Filtering took place within such a high frequency range in order to specifically identify trials that contained muscle activity. Again, trials that exhibited a z-value higher than a certain threshold (mostly 30, but this had to be adjusted for some participants) were marked as bad trials. After identification of bad channels and trials, the continuous data was read from disk, filtered between 0.1 and 30 Hz with a non-causal zero-phase two-pass 5th order Butterworth IIR filter with -6 dB half-amplitude cutoff. Then, data was segmented into trials, without the ones marked as bad in the earlier pre-processing step. A vertical and a horizontal eye channel were computed as difference waves between the two vertical and two horizontal eye electrodes. Then, data were submitted to an Independent Component Analysis (ICA; Jung et al., 2000) in order to extract and subsequently exclude components related to eye movement, remaining muscle activity, and heartbeat. For the ICA, data were high-pass filtered at 1 Hz in order to improve stationarity of the components. After the removal of artefactual components, the remaining components were back-projected to the original, 0.1-Hz-filtered data. Then, data were visually screened for trials that contained artifacts that survived the AAR and the ICA procedures, which were then removed.

For each participant, each condition, each trial, and each channel, we extracted three mean voltage values of interest: in a pre-stimulus time window (150 - 5 ms before the onset of the critical word), in the N400 time window (300 - 500 ms after onset of the critical word), and in the P600 time window (600 - 900 ms after onset of the critical word). These values were not baseline corrected, because we included the pre-stimulus activity as a factor in the analysis (for a description of this method see Alday, 2019). Critical words were at the fourth position in conditions TVRR and TVDO, at the eighth position in conditions IVWR and TVRR, and at the fifth position in conditions ORAI and ORIA.

### IAF

IAF was quantified from participants’ eyes-closed resting state EEG before and after the experiment. The two-minute segments were cut into 60 two-second trials. Data were band-pass filtered between 0.1 and 30 Hz with a non-causal zero-phase two-pass 5th order Butterworth IIR filter with -6 dB half-amplitude cutoff and re-referenced to linked mastoids. Then, only eye channels and only 9 postero-occipital channels (Pz, P1, P2, POz, PO3, PO4, Oz, O1, O2) were retained. A vertical and a horizontal eye channel were computed as difference waves between the two vertical and two horizontal eye electrodes, respectively. An automatic artifact rejection procedure computed z-values in the horizontal and vertical eye channels per time point per trial and if a z-value at any time point in a trial exceeded 4, this trial was marked as bad. If any of the chosen channels had been marked as a bad channel in the main experiment (see above), they were interpolated using spline interpolation (Perrin, Pernier, Bertnard, Giard, & Echallier, 1987). With the *restingIAF* function from the *restingIAF* toolbox (Corcoran, Alday, Schlesewsky, & Bornkessel-Schlesewsky, 2018), we calculated power spectral density between one and 30 Hz for each channel and smoothed them with a Savitzky-Golay filter (Savitzky & Golay, 1964, with a frame width of 11 and a polynomial degree of 5). The function looked for evidence for peak activity in the smoothed power spectra between 5 and 14 Hz and quantified IAF for each channel following the peak alpha frequency as well as the centre of gravity methods. In order for the function to yield an average IAF quantification, a minimum of three channels had to yield an individual quantification. IAF estimates before and after the main experiment were averaged. Peak alpha frequency and centre of gravity IAF quantifications were highly correlated (*r* (24) = 0.94, *p* < 0.001), but the centre of gravity method yielded an IAF value for 30 of the 32 participants, while the peak alpha frequency method only yielded an IAF value for 26 participants. We therefore chose centre of gravity IAF for further calculations. The IAF of the two participants without estimate was interpolated with the median IAF of the whole sample.

### MMN

For a quantification of participants’ MMN, we presented participants the Passive Auditory Oddball MMN paradigm from the ERP CORE package by Emily S. Kappenman and Steven J. Luck while their EEG was recorded. Participants listened to a total of 290 1000 Hz sine wave tones with a duration of 100 ms including 5 ms rise and fall times, 230 of which were presented at a standard volume of 80 dB and 60 of which were presented at a deviant volume of 70 dB. The inter-stimulus interval was jittered between 450 and 550 ms. Before the experimental trials, the standard sine wave tone was presented in ten warm-up trials, which were excluded from the analysis. Participants were instructed to watch a silent movie during the presentation of the sounds. During preprocessing, the EEG was first band-pass filtered between 0.1 and 30 Hz with a non-causal zero-phase two-pass 5th order Butterworth IIR filter with -6 dB half-amplitude cutoff and segmented into trials of 580 ms length; a 200 ms prestimulus baseline and 380 ms after stimulus onset. Then, a vertical and a horizontal eye channel were computed as difference waves between the two vertical and two horizontal eye electrodes, respectively. Then, the same automatic artifact rejection procedure as in the IAF quantification was applied, and any channels marked as bad in the main experiment (see above) were interpolated using spline interpolation (Perrin et al., 1987). Furthermore, data were re-referenced to linked mastoids. Following Duncan et al. (2009), we chose a frontocentral cluster encompassing Fc, FCz, Cz, FC1, and FC2 as the location of the MMN. The difference wave of ERP traces in response to deviant vs. standard tones was calculated and averaged across all channels of the MMN cluster per participant. We quantified the MMN as the negative peak amplitude measured between 110 and 180 ms after sound onset.

## Statistical Analyses

Behavioral and EEG data were analyzed in R Version 3.6.2 (R Core Team, 2018). Linear mixed effects models (LMEMs) were fitted using the package *lme4* (Bates, Mächler, Bolker, & Walker, 2015).

For the analysis of differences in acceptability scores between the conditions in the RRC and ORC paradigms, two separate LMEMs with repeated contrasts were run. A repeated contrasts model has the advantage of only comparing neighboring factors, thereby reducing the number of statistical tests (Schad, Vasishth, Hohenstein, & Kliegl, 2020).

For the ERP analysis, in order to reduce the levels of the channel dimension of the EEG data while still remaining free of assumptions regarding the topography of our effects to avoid “double dipping” (Kriegeskorte, Simmons, Bellgowan, & Baker, 2009), channels were clustered regarding the two factors laterality (left: F7, F5, F3, FC5, FC3, T7, C5, C3, TP7, CP5, CP3, P7, P5, P3, PO7, PO3, FT7; medial: F1, F2, Fz, FCz, FC1, FC2, C1, C2, Cz, CP1, CP2, CPz, P1, P2, Pz, POz; right: F8, F6, F4, FC6, FC4, T8, C6, C4, TP8, CP6, CP4, P8, P6, P4, PO8, PO4) and sagittality (anterior: F7/8, F5/6, F3/4, F1/2, Fz, FC5/6, FC3/4, FC1/2, FCz, FT7/8, T7/8, C5/6, C3/4, C1/2, Cz; posterior: TP7/8, CP5/6, CP3/4, CP1/2, CPz, P7/8, P5/6, P3/4, P1/2, Pz, PO7/8, PO3/4, POz), and voltage values per cluster were obtained by averaging across channels.

We fitted LMEMs to predict ERP amplitude in the N400 (ORAI-ORIA comparison) and P600 (TVRR-TVDO and IVWR-TVRR comparisons) time windows on a trial-by-trial basis.

We first fitted a basic model for each comparison, predicting N400 or P600 amplitude. The models always included a factor of condition with two levels, thereby mimicking a direct comparison between conditions, like traditional ERP analyses. The factor *condition* was encoded via treatment coding, with the “baseline” conditions (TVDO in the TVRR-TVDO comparison, TVRR in the IVWR-TVRR comparison, and ORIA in the ORAI-ORIA comparison) being coded as 0 and the ERP-component-eliciting condition being coded as 1. Other fixed effects were pre-stimulus amplitude (Alday, 2019; Alday, Schlesewsky, & Bornkessel-Schlesewsky, 2017), an interaction term between pre-stimulus amplitude and condition, and full main effects as well as interactions of condition, laterality, and sagittality. Laterality and sagittality were encoded via sum coding. Random factors included a random slope of condition per participant as well as random intercepts of participant and item. Please note that *item* denotes a single sentence and not a sentence cluster. This is a prototypical model formula for the basic models: ERP amplitude *∼* prestim*condition + condition * laterality * sagittality + (condition|participant) + (1|item).

To investigate a potential moderating influence of our variables of interest (VOI), which consisted of PTA, RS, OS, IAF, Flanker, and Stroop (see Table 3 for a correlation matrix of the VOI as well as age), we updated the basic models by adding each VOI separately to the interaction term of condition, laterality, and sagittality. In the PTA models, participant-controlled attenuation/amplification residualized for PTA was also added to the models as a fixed effect, to control for effects due to insufficient amplification of the stimuli. Random factors included a random slope of condition per participant as well as random intercepts of participant and item. The prototype of all formulae was as follows: ERP amplitude *∼* prestim*condition + condition * laterality * sagittality * VOI + (condition|participant) + (1|item).

**Table 3.**
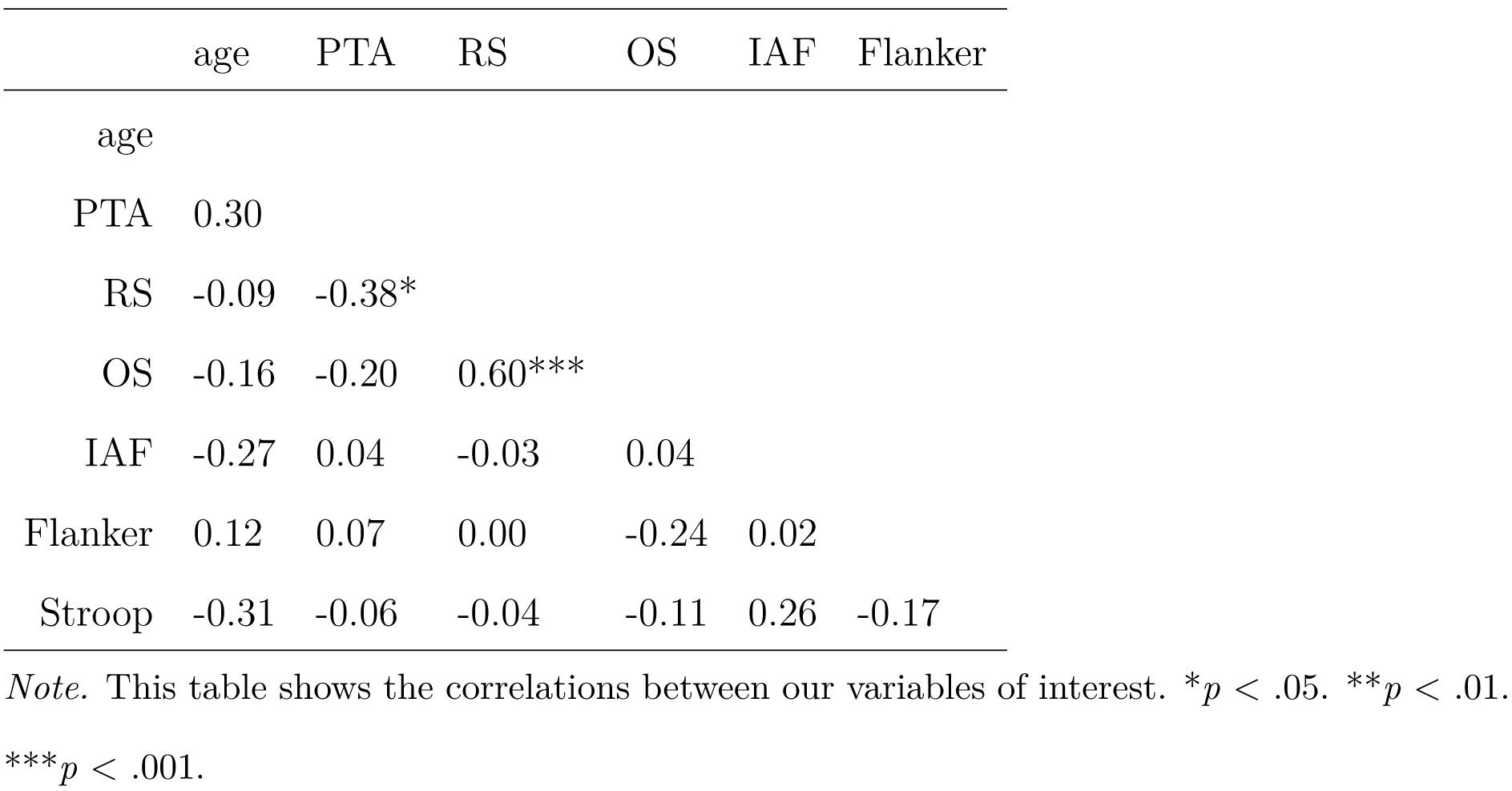
Correlation Matrix of Variables of Interest.

|  | age | PTA | RS | OS | IAF | Flanker |
| --- | --- | --- | --- | --- | --- | --- |
| age |  |  |  |  |  |  |
| PTA | 0.30 |  |  |  |  |  |
| RS | -0.09 | -0.38* |  |  |  |  |
| OS | -0.16 | -0.20 | 0.60*** |  |  |  |
| IAF | -0.27 | 0.04 | -0.03 | 0.04 |  |  |
| Flanker | 0.12 | 0.07 | 0.00 | -0.24 | 0.02 |  |
| Stroop | -0.31 | -0.06 | -0.04 | -0.11 | 0.26 | -0.17 |

*Note.* This table shows the correlations between our variables of interest. \* $p < .05$ . \*\* $p < .01$ .
\*\*\* $p < .001$ .

We chose to report and interpret only models that fulfil the following criteria: First, we needed to make sure that our VOI is indeed a better predictor than chronological age. Therefore, the model with a certain of our variables of interest needed to have a better fit as measured by the Akaike Information Criterion (AIC; Akaike, 1974) to the data than chronological age. Second, the model needed to exhibit at least one significant interaction effect between condition and the VOI, signaling a moderation of ERP amplitude by the VOI. Although only the models which fulfil these criteria are reported in the text, all fitted models are reported in Supplementary Tables S9-S32.

Finally, we calculated Pearson correlations between MMN amplitude and each of the VOI.

We further analyzed how our VOI would predict acceptability ratings of the sentences in the conditions we analyzed the ERPs from. To this end, cumulative link mixed models (CLMMs) were fitted by means of the *ordinal* package (Christensen, 2019) with the following formula: rating *∼* condition * VOI + (condition|participant) + (1|item).

## Results

### Behavioral Results

For the LMEMs with repeated contrasts used to test for differences in the acceptability ratings between the conditions in the RRC paradigm, the conditions were ordered as follows: We expected the lowest ratings for the grammatically incorrect IVWR sentences, the second-lowest ratings for the temporarily ambiguous TVRR sentences, the second-highest ratings for the TVDO sentences, and the highest ratings for the IVCO sentences. The difference between IVWR and TVRR ratings was significant (*b* = 0.71, *t*(84) = 7.11, *p* < 0.001), as was the difference between TVRR and TVDO ratings (*b* = 0.41, *t*(84) = 4.14, *p* < 0.001). The difference between TVDO and IVCO ratings was not significant (*b* = 0.18, *t*(84) = 1.81, *p* = 0.07). Scores are shown in Figure 1, left panel.

**Figure 1.**
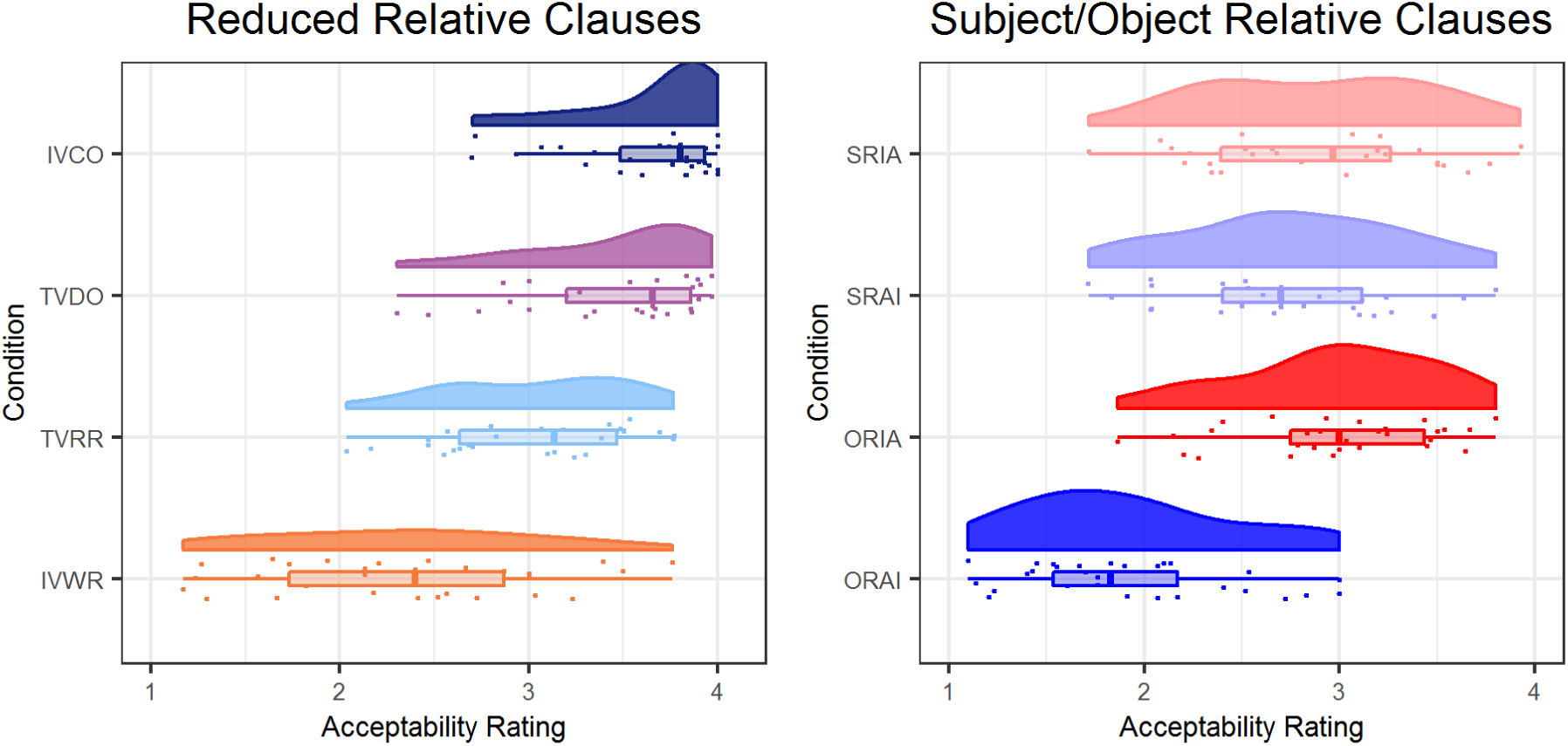
This figure shows the distributions of acceptability ratings in the RRC (left) and ORC (right) paradigms.

For the ORC paradigm, the conditions were ordered as follows: We expected the lowest ratings for the ORAI sentences, the second-lowest ratings for the ORIA sentences, the second-highest ratings for the SRIA sentences, and the highest ratings for the SRAI sentences. The difference between ORAI and ORIA ratings was significant (*b* = 1.06, *t*(84) = 15.12, *p* < 0.001), but the difference between ORIA and SRIA ratings was not (*b* = 0.10, *t*(84) = 1.38, *p* = 0.17). The difference between SRIA and SRAI ratings was significant (*b* = 0.14, *t*(84) = 2.00, *p* = 0.049). Scores are shown in Figure 1, right panel.

### ERP Results

#### RRC: TVRR-TVDO Comparison

The first comparison in the RRC paradigm addressed ERP amplitude in the P600 time window in response to the fourth position in the TVRR sentences vs. the TVDO sentences (“The broker persuaded *to*…” vs. “The broker persuaded *the*…”).

The basic model did not contain a significant main effect of condition nor a significant interaction effect between condition and laterality or sagittality (see also Figure 2). However, this was not a hindrance for the following analyses, because the aim of the present study was to identify variables that would distinguish between participants who show a P600 and those who do not.

**Figure 2.**
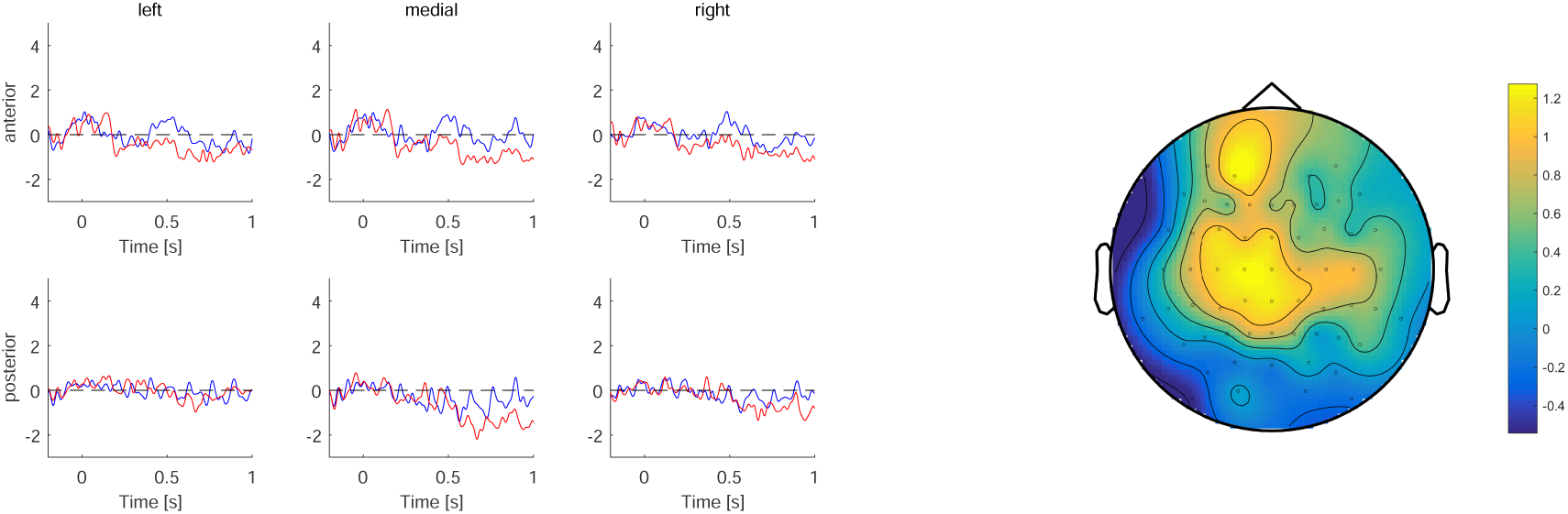
Left: Grand average ERPs centered at the start of the word at position 4 of TVRR (blue) vs. TVDO (red) sentences. Right: Topographic map of difference wave voltage in µV averaged across the P600 time window (500-900 ms after critical word onset)

Regarding the models containing the VOI, we first compared the fitted models to the same model fitted with age instead of the VOI and only kept those models that had a lower AIC than the model with age (see Table 4 for an overview of evidence ratios; Wagenmakers & Farrell, 2004). In the TVRR-TVDO comparison, all VOI models except for the IAF model had a lower AIC than the age model.

**Table 4.**
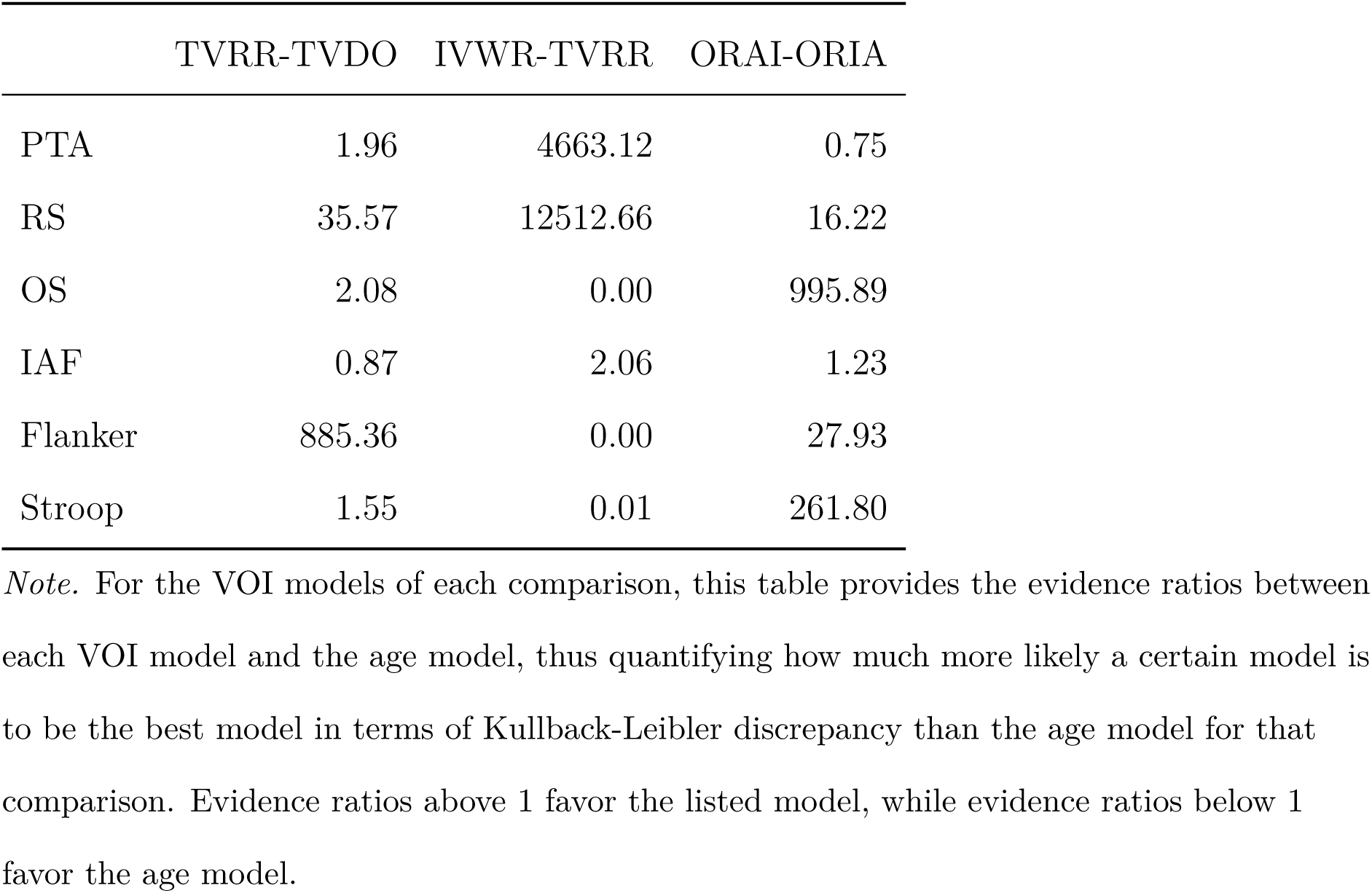
AIC evidence ratios for VOI models against age models.

In a second step, we checked whether the remaining models contained a significant interaction effect between condition and the VOI, signaling a moderation of ERP amplitude by the VOI. Only the PTA and RS models contained a significant interaction effect with condition. Effects plots of the interactions can be found in Figure 5. To view these effects for each cluster separately, see Supplementary Figure S1.

In the PTA model, the interaction effect of condition and PTA was significant, *b* = 0.65, *t*(27.97) = 2.39, *p* = 0.02. Across the topography, participants with higher hearing thresholds (i.e. worse hearing) exhibited a larger P600 than participants with lower hearing thresholds (i.e. better hearing).

In the RS model, the interaction effect of condition and RS was significant, *b* = -0.68, *t*(28.98) = -2.44, *p* = 0.02. Across the topography, participants with higher RS scores (i.e. better working memory) exhibited a smaller P600 than participants with lower RS scores (i.e. worse working memory).

#### RRC: IVWR-TVRR Comparison

The second comparison in the RRC paradigm involved the eighth position of the IVWR sentences vs. the TVRR sentences (“The broker persuaded to sell the stock *was*…” vs. “The broker planned to sell the stock *was*…”).

The basic model contained significant interaction effects between condition and laterality (medial), *b* = 0.40, *t*(4292.78) = 2.10, *p* = 0.04, and between condition and sagittality, *b* = 0.45, *t*(4296.97) = 3.36, *p* = 0.001, indicating that the IVWR sentences were more positive than the TVRR sentences at medial as well as posterior channels (see also Figure 3). IVWR sentences relative to TVRR sentences elicited a P600 at the eighth position.

**Figure 3.**
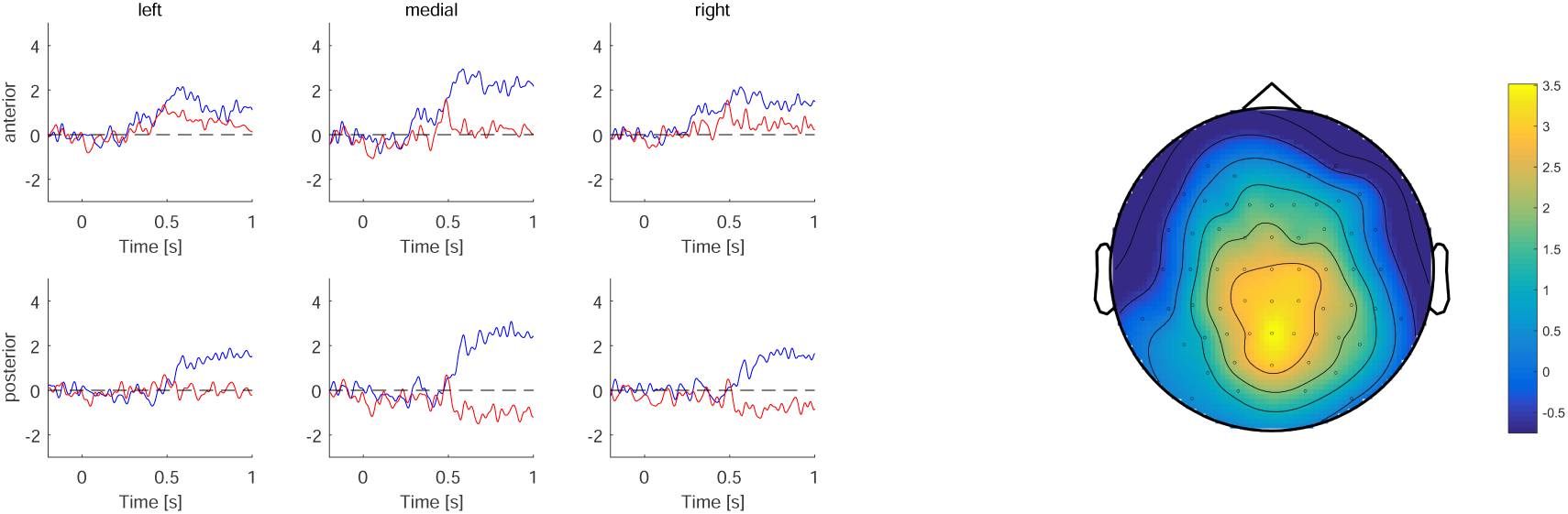
Left: Grand average ERPs centered at the start of the word at position 8 of IVWR (blue) vs. TVRR (red) sentences. Right: Topographic map of difference wave voltage in µV averaged across the P600 time window (500-900 ms after critical word onset)

By comparing the models fitted with the VOI to the same model fitted with age instead of the VOI, we found that PTA, RS, and IAF had a lower AIC than the age model. Only the IAF model contained a significant interaction effect with condition. An effects plot of the models can be found in Figure 5.

In the IAF model, there was a significant interaction effect of condition and IAF, *b* = 0.85, *t*(27.20) = 2.46, *p* = 0.02. Across the topography, participants with a higher IAF exhibited a larger P600 than participants with a lower IAF.

#### ORC

ERP amplitudes in response to the fifth position of ORIA vs. ORAI sentences were compared (“The accident that the *musician*…” vs. “The musician that the *accident*…”). This comparison took place in the N400 time window.

The basic model did not contain a significant main effect of condition nor a significant interaction effect between condition and laterality or sagittality (see also Figure 4).

**Figure 4.**
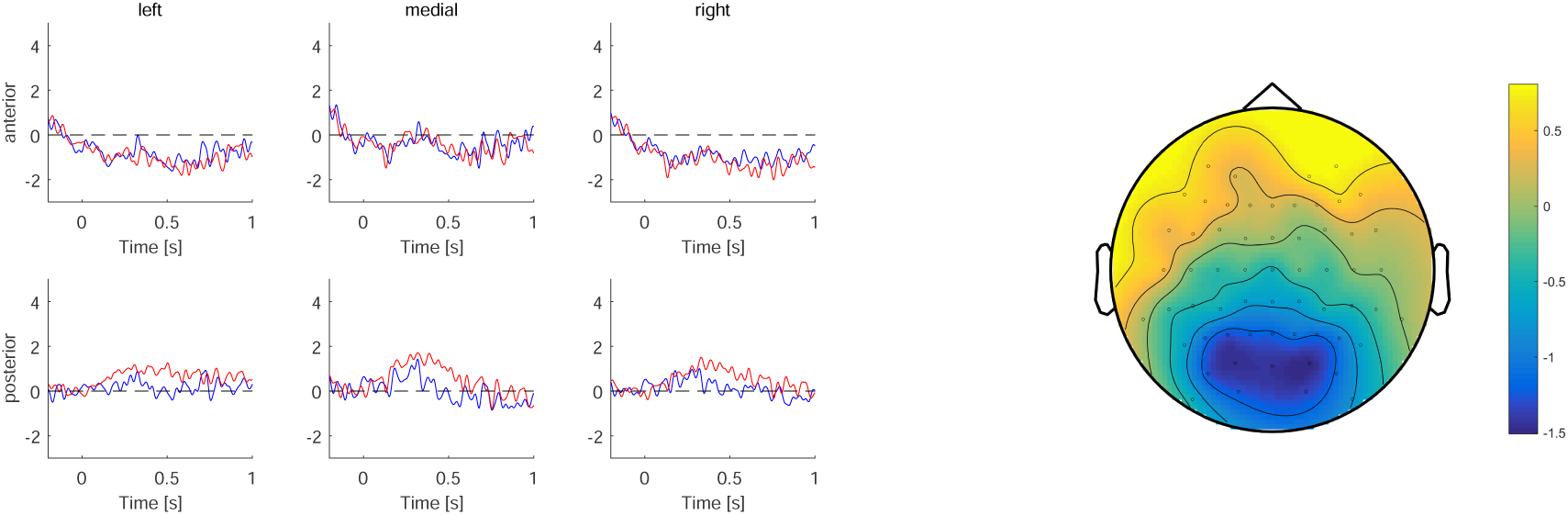
Right: Grand average ERPs centered at the start of the word at position 5 of ORAI (blue) vs. ORIA (red) sentences. Right: Topographic map of difference wave voltage in µV averaged across the N400 time window (300-500 ms after critical word onset)

**Figure 5.**
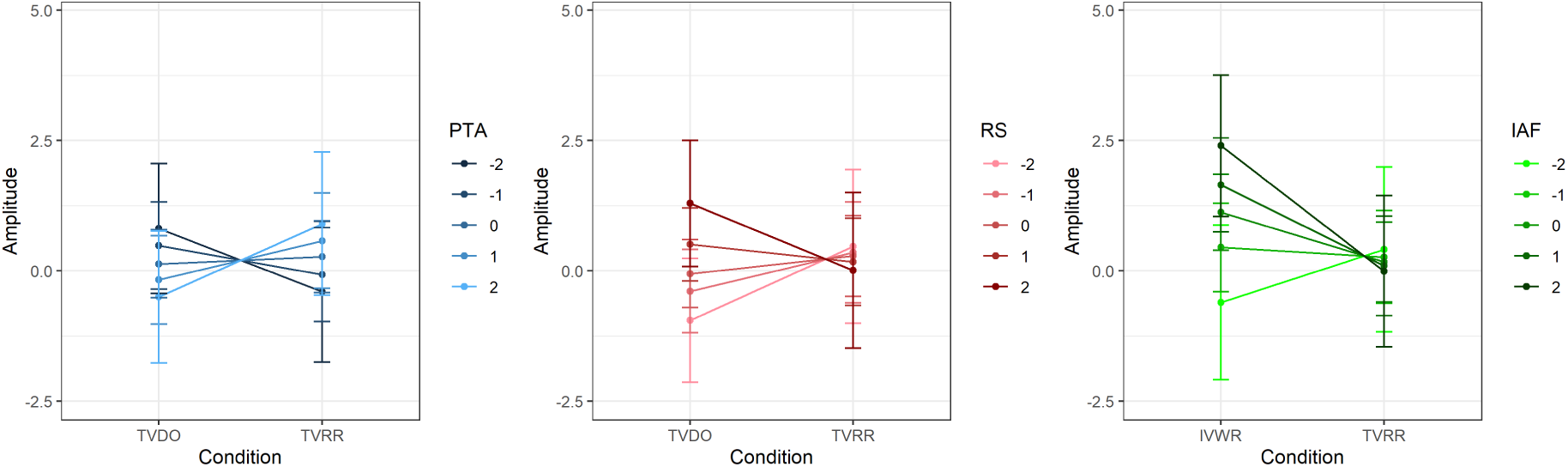
Effects plots of P600 amplitude of the models with a significant condition*VOI interaction. VOI values were z-scored. Left: Effects plot of P600 amplitude by condition*PTA interaction. Middle: Effects plot of P600 amplitude by condition*RS interaction. Right: Effects plot of P600 amplitude by condition*IAF interaction.

By comparing the models fitted with the VOI to the same model fitted with age instead of the VOI, we found that all VOI models except for the PTA model had a lower AIC than the age models. However, none of the models contained a significant interaction effect between condition and the VOI.

#### MMN

The grand averages of the MMN experiment and the topography of the difference wave are shown in Figure 6. We first tested for the presence of the MMN by running a one-sample two-sided t-test of the MMN amplitude against zero. The test showed that MMN amplitude was significantly lower than zero, *m* = -4.66, *t*(31) = -10.61, *p <* 0.001.

**Figure 6.**
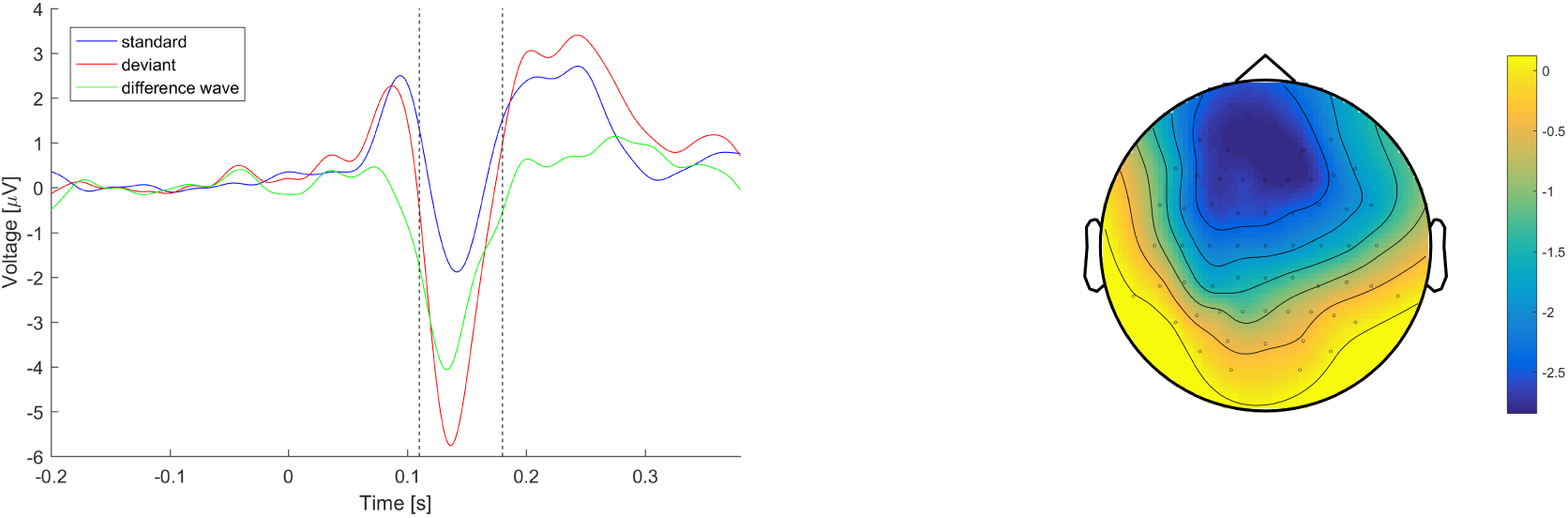
Left: Grand average ERP traces in response to the standard (blue) and deviant (red) sounds as well as the difference wave of the two traces (green). Right: Topographic map of difference wave voltage in µV between 110 and 180 ms after sound onset

In a next step, we calculated six Person correlations between MMN amplitude and each of the VOI. None of the correlation coefficients was significant. There was no evidence for a modulation of the MMN by hearing thresholds or cognitive ability.

#### Acceptability Ratings by VOI

As a next step, we aimed to ascertain whether the VOI would, in addition to moderating ERP differences, also moderate acceptability rating differences. For the three data sets with a significant condition*VOI interaction, we fitted CLMMs to the acceptability ratings, again on a single-trial basis. However, none of the three predictors (PTA and RS for TVRR-TVDO sentences, IAF for IVWR-TVRR sentences) showed a significant interaction effect with condition in these models.

## Discussion

In the present study, we investigated how syntactically difficult sentence material is processed by healthy older adults differing in perceptual and cognitive abilities.

Specifically, we presented older adults with two different paradigms, probing both reanalysis and actor computation, and related the resulting ERPs to their hearing and cognitive abilities.

### Individual Differences in Reduced Relative Clause Processing

Starting with the reanalysis paradigm, we found a clear acceptability hierarchy in our four conditions. The unproblematic IVCO (intransitive verb, correct) and TVDO (transitive verb, direct object) sentences were rated highest, followed by the temporarily ambiguous TVRR (transitive verb, reduced relative) sentences, and then by the grammatically incorrect IVWR (intransitive verb, wrong) sentences.

In the ERP analysis, we probed processing of the TVRR sentences at two points in time. First, we compared ERPs in response to the word at the fourth position of the TVRR sentences (i.e. right at that point in time when the ambiguity was resolved) to ERPs in response to the word at the fourth position of the TVDO sentences, which began in the same way as the TVRR sentences, but continued with the preferred interpretation. Across the sample, there was no significant difference between the two conditions in the P600 time window. This was not a hindrance for the following analyses, because it is entirely possible that there was no difference in the grand average means because there were more participants who did not show a P600 effect than participants who did show a P600 effect. The aim of the present study was to identify variables that would distinguish between these participants. The analyses involving our participant-level VOI (hearing thresholds, working memory, IAF, and inhibition) revealed that participants with worse peripheral hearing and participants with lower working memory capacity exhibited a P600 effect in response to TVRR sentences relative to TVDO sentences. Both of these effects were not specific to any topographical region, but were distributed broadly across the scalp.

Second, we compared ERPs in response to the eighth position of the TVRR sentences to ERPs in response to the eighth position of the IVWR sentences. This comparison allowed us to test for successful reanalysis of the TVRR sentences towards the dispreferred RRC interpretation. If reanalysis of the TVRR had been successful, the “was” at the eighth position would be a necessary component of the sentence. If reanalysis had not been successful, and instead, participants had gone with a “good-enough” interpretation of the sentence up until that point, then the “was” would render the sentence ungrammatical, just as in the IVWR condition. This in turn implies that a between-condition difference in the ERPs in the P600 time window would be indicative of reanalysis success: if there is no difference, reanalysis was unsuccessful, whereas if there is a difference, reanalysis was successful. Across the sample, there was a significant difference between the conditions at medial and posterior channels, thus indicating that, overall, our participants could discriminate between the temporarily ambiguous TVRR sentences and the ungrammatical IVWR sentences. This is also reflected in the significant difference in acceptability ratings between the two conditions.

We again tested whether our VOI would predict the ERP difference between the conditions. Participants with a higher IAF exhibited a higher P600 effect than participants with a lower IAF. This suggests that participants with a higher IAF were more successful in reanalysis. In summary, we found that hearing thresholds, working memory, and IAF predicted reduced relative clause processing at different stages. Inhibition, by contrast, was not found to modulate the amplitude of ERP indicators of reduced relative clause processing.

Overall, an interesting pattern emerged from these two complementary analyses. The comparison at the first point in time revealed stronger effects for participants with worse hearing and lower working memory capacity. On the other hand, at the second point in time, the effects were stronger for participants with a higher IAF.

How can these findings be reconciled? First of all, this pattern suggests that different processing strategies were favored by different participants depending on their hearing and cognitive abilities. In this paradigm, this may be a result of a parallel parsing strategy (Fiebach, Vos, & Friederici, 2004; Frisch, Schlesewsky, Saddy, & Alpermann, 2002), i.e., simultaneous activation of multiple interpretations of the temporarily ambiguous sentence. It is possible that our better-hearing as well as our high-span participants simultaneously activated both the preferred and the dispreferred interpretation (see M. C. MacDonald, Just, & Carpenter, 1992). By contrast, the worse-hearing and the low-span participants only activated the preferred interpretation, thus resulting in higher processing effort, as reflected in a larger P600, when the ambiguity was resolved towards the dispreferred interpretation. Correspondingly, our higher-IAF participants exhibited a larger P600 at the later comparison point, thus indicating a higher likelihood of a successful reanalysis having taken place. We suggest that this pattern may reflect a dissociation between the effort required by the reanalysis and the likelihood of correctly computing the target interpretation. While reanalysis cost is dependent on cognitive resources and is therefore higher for individuals with worse hearing and lower working memory capacity, the likelihood of reanalysis success depends on IAF. This intriguing result will be further explored in the Implications section below.

A resource-based view could explain why the results with hearing thresholds are very similar to the results with working memory span for the TVRR-TVDO comparison. Several studies have tested the “effortfulness hypothesis”, which posits that successful perception in the face of degraded input (e.g. because of raised hearing thresholds) consumes resources which are then missing in downstream processing steps such as memory encoding (McCoy et al., 2005; Tun, Benichov, & Wingfield, 2010; Tun, McCoy, & Wingfield, 2009). This hypothesis could also explain our results for the TVRR-TVDO comparison. Possibly, participants with lower hearing thresholds deploy fewer resources in order to achieve successful perception of the sensory input, which would in turn allow them to allocate more resources to keeping both the preferred and the dispreferred interpretation in memory. Additionally, participants with a higher working memory capacity would have more resources available in general, and therefore, a higher recruitment of resources during perception would still allow participants with a larger resource pool to keep both interpretations of the RRC in memory.

### Individual Differences in the Orocessing of Object Relative Clauses

In the object relative clause / actor computation paradigm, we found that ORAI (object-relative, animate - inanimate) sentences were clearly rated as least acceptable. ORIA (object-relative inanimate - animate) and SRIA (subject-relative, inanimate - animate) sentences did not differ in their ratings, and SRAI (subject-relative, animate - inanimate) sentences were only slightly more acceptable than SRIA sentences. We expected this difference in acceptability ratings within the OR clauses due to animacy, with previous studies demonstrating that animacy is an important cue for OR clause processing (DeDe, 2015; Traxler et al., 2002; Weckerly & Kutas, 1999).

In the ERP analysis, we probed actor computation in the ORAI sentences compared to the ORIA sentences. Specifically, we compared ERPs in response to the subject of the relative clause (fifth position). Based on previous research showing processing difficulties for inanimate object-relative clause subjects as compared to animate object-relative clause subjects (DeDe, 2015; Traxler et al., 2002; Weckerly & Kutas, 1999), we expected an N400 for ORAI sentences in comparison with ORIA sentences.

Across the sample, there was no significant difference between the two conditions in the N400 time window. Again, this was not a hindrance for the VOI analyses, because the aim of the present study was to identify variables that would distinguish between these participants.

We again tested whether our VOI would predict the ERP difference between the conditions. However, although almost all models with the VOI provided a better fit to the data than models including only age, none exhibited a significant interaction with N400 amplitude. This was surprising, given the vast literature on ORC processing in older and hearing-impaired adults (e.g. DeCaro et al., 2016; Wingfield et al., 2006, 2003). It is possible that the manipulation was simply not strong enough to reliably elicit an N400 in enough participants. In comparison to the RRC paradigm, where we analyzed responses to ungrammatical (IVWR) and dispreferred (TVRR) sentences, here in the ORC paradigm, the sentences were perfectly grammatical, albeit with a non-prototypical animacy configuration. Older adults as a group may, as a result of their experience, have had a high degree of exposure to inanimate agents and therefore would not necessarily rely on an internal model that favors animate agents.

In order to examine between-participant variability for this comparison more directly, we plotted the random slopes of condition per participant for N400 amplitude derived from the basic ORAI vs. ORIA model. Random slopes were indeed rather variable, and almost equally distributed to the right and to the left of the zero line (see Supplementary Figure S2, left panel).

As the study by Weckerly and Kutas (1999) only found the effect in question for good comprehenders, we conducted an additional analysis to ascertain whether N400 amplitude in the most difficult ORAI condition would be related to acceptability ratings (see Supplementary Table S33 and Figure Supplementary Figure S3). Participants with a larger (= more negative) N400 were less likely to give a low rating to the ORAI sentences than participants with a smaller N400. Assuming that good comprehenders would be more likely to give a good rating, this result suggests that N400 amplitude and comprehension are related in a similar way as in the Weckerly and Kutas (1999) study. Interestingly, this effect does not appear to be predicted by any of our VOI.

### VOI and Behaviour

As a follow-up analysis, we analyzed whether the VOI that moderated ERPs would also moderate acceptability ratings. However, none of the VOI (PTA and RS for the TVRR-TVDO comparison and IAF for the IVWR-TVRR comparison) moderated acceptability rating differences. This is not entirely surprising given that neurophysiological data typically show more sensitivity to certain manipulations than behavioral data and are sometimes even used to test for differences in effort in the face of similar behavioral outcomes (see, for example, Bornkessel, McElree, Schlesewsky, & Friederici, 2004; Rolke, Heil, Streb, & Hennighausen, 2001).

### Mismatch Negativity (MMN)

We included a MMN paradigm in the study in order to test whether the modulatory influence of hearing and cognitive abilities would also extend to pre-linguistic auditory ERP components. If this were the case, our VOI would arguably modulate central auditory processing in general, irrespective of the linguistic computations necessary for sentence comprehension. However, there was no correlation between MMN amplitude and any of our VOI. While we do not wish to take the absence of evidence for the evidence for absence, we nevertheless at least see a much stronger effect of the VOI on sentence processing than on central auditory processing in general.

### Implications

Overall, we observed modulation of ERPs by hearing and cognitive abilities at two different stages of RRC processing.

The finding that sentence comprehension (and, thereby, also sentence processing) is predicted by hearing impairment is well established, especially in older adults (Wingfield et al., 2006). However, in these studies, participants are usually grouped depending on whether their sine wave perception exceeds a certain sound level threshold or not. Our findings on hearing thresholds could be considered surprising, because, if our sample had been clinically tested for their hearing ability, most, if not all of them, would likely have been classified as having normal hearing. Nevertheless, we found a significant relationship between hearing thresholds and ERP amplitudes in the RRC paradigm. A study by Ayasse et al. (2019) found that even in young adults who pass a screen for normal hearing, slightly elevated hearing thresholds detrimentally affected processing of difficult syntactic constructions. This suggests that it is important to consider hearing thresholds as continuous variables rather than considering people within certain threshold ranges as homogeneous groups.

We have explored these results in light of the “effortfulness hypothesis”. The results can also be considered from the perspective of the predictive coding framework. This theory of brain function describes the brain as an empirical Bayesian device that continually aims to minimize prediction error, which is “the difference between the input observed and that predicted by the generative model” (Friston, 2005, p. 821). This principle is implemented at all levels of the cortical hierarchy. Prediction error results from a mismatch between the sensory input that propagates to higher cortical levels by means of feedforward connections and the prediction of the generative model of the environment that is projected to lower cortical levels by feedback connections (Friston, 2005, 2010). Prediction error can also result in an update of the generative model, which serves the purpose of minimizing prediction error in the future when confronted with similar input. As Moran, Symmonds, Dolan, and Friston (2014) propose, aging can be viewed as reflecting “a progressive refinement and optimization of generative models” (Moran et al., 2014, p. 1). They note that the often observed attenuation of older adults’ evoked responses compared to those of younger adults may be due to older adults’ accumulation of sensory experience, resulting in less model updating.

Conceptually preceded by the similar account of *analysis by synthesis* (Bever & Poeppel, 2010; Halle & Stevens, 1962), the notion of such generative models is prolific in language comprehension research (e.g. Bornkessel-Schlesewsky & Schlesewsky, 2019; Pickering & Garrod, 2007, 2013). Based on Moran et al. (2014), one would therefore expect older adults to have a higher tendency to refrain from updating their internal model after encountering an error in that model. This absence of model updating would result in a non-updated version of e.g. a garden-path sentence and could explain the difference between younger and older adults in adopting a “good-enough” interpretation of garden-path sentences (Christianson et al., 2006). However, as there is typically considerable inter-individual variability in older adults, also in language-related ERP research (Bornkessel-Schlesewsky et al., 2015; DeLong et al., 2012), it is useful to examine the individual differences that underlie this variability. In our study, IAF moderated the P600 amplitude difference between the ungrammatical IVWR and the reduced relative TVRR sentences. Although it is still unclear how exactly IAF is related to cognitive performance, an association between the two has been found repeatedly, and it has been suggested that IAF reflects cognitive performance at the level of general intelligence (Grandy, Werkle-Bergner, Chicherio, Lövdén, et al., 2013) rather than a specific cognitive ability per se. A similar account proposes that a high IAF reflects a trait or state that fosters optimal cognitive performance rather than optimal cognitive performance itself (“cognitive preparedness”, Angelakis, Lubar, Stathopoulou, & Kounios, 2004). Evidence corroborating this hypothesis on the metabolic level showed that IAF is positively associated with regional cerebral blood flow (Jann, Koenig, Dierks, Boesch, & Federspiel, 2010), which facilitates rapid reorientation during cognitive tasks.

Returning to the results of our study, this notion of IAF as fostering mental flexibility and reorientation can also be applied to the reanalysis of sentences in which an ambiguity has been resolved towards a dispreferred interpretation. The larger P600 in the IVWR-TVRR comparison for participants with higher IAFs would therefore reflect their stronger inclination towards reanalysis. To put it in predictive coding terms: participants with a higher IAF were more inclined to update their internal model of the TVRR sentence, thus leading to a higher likelihood of the target reading being correctly computed.

In the ORC paradigm, we did not observe a modulation of ERP amplitude by hearing or cognitive ability. However, following the results of Weckerly and Kutas (1999) and assuming a relation between their comprehension scores and our acceptability scores, a larger N400 was related to a better acceptability rating of the ORAI sentences. Apparently, the N400 in this manipulation is more strongly related to the outcome of sentence processing than to any of our VOI. Considering two-component theories of intelligence that posit a “fluid” and a “crystallized” set of cognitive abilities (Cattell, 1971; Horn, 1982; Hülür, Gasimova, Robitzsch, & Wilhelm, 2018), it is possible that the N400 would be better explained by a crystallized form of cognition like vocabulary size than by one of our cognitive VOI, all of which represent fluid cognitive measurements.

Future research should address whether the N400 amplitude in this comparison can be predicted with crystallized rather than fluid cognitive abilities. Also, it should try to discover how hearing thresholds and working memory relate to ORC processing at the neural level, thus linking back to previous behavioral studies (Amichetti et al., 2016; DeCaro et al., 2016; Wingfield et al., 2006, 2003).

## Conclusion

In the present study, we examined how hearing thresholds, working memory, IAF, and inhibition influence auditory sentence processing in healthy older adults. We found that hearing thresholds, working memory, and IAF modulated RRC processing at different time points. We did not observe a modulation of processing of ORCs differing in their animacy configuration, possibly due to the more subtle nature of the manipulation. In conclusion, there is no single hearing-related or cognitive variable that can be considered beneficial for auditory sentence comprehension in general, but it depends on the phenomenon in question.

## Acknowledgements

This work was supported by grants from the Swiss National Science Foundation (grant no. 172268 and 172268/2 to IK). During the work on her dissertation, IK was a pre-doctoral fellow of the International Max Planck Research School on the Life Course (LIFE). IBS acknowledges the support of an Australian Research Council Future Fellowship (FT160100437). The authors thank Madison Jade Richter for her extensive assistance in data acquisition and Maria Kliesch for German versions of the PsychoPy experiments.

