## Supplementary Tables for "Individual Differences in Peripheral Hearing and Cognition Reveal Sentence Processing Differences in Healthy Older Adults"

Table S1: Sentence stimuli of the TVDO condition.

1. The broker persuaded the investor to conceal the transaction.
2. The man hired the salesperson to help the store.
3. The doctor implored the relatives to see the patient.
4. The reporter selected the photographer to get the story.
5. The woman advised the child to see the play.
6. The senator forced the assistant to chair the committee.
7. The judge advised the jury to sentence the defendant.
8. The tailor hired the sewer to fix the suit.
9. The swimmer urged the athlete to lose weight.
10. The general permitted the officer to leave the army.
11. The minister invited the priest to give the sermon.
12. The mechanic trusted the trainee to repair the car.
13. The singer allowed the substitute to perform the opera.
14. The teacher urged the professor to improve his teaching.
15. The burglar induced the friend to rob the bank.
16. The policeman ordered the guard to watch the bank.
17. The dentist invited the lawyer to meet the actress.
18. The janitor persuaded the plumber to fix the faucet.
19. The nurse induced the relatives to leave the patient.
20. The prince advised the duke to marry the princess.
21. The journalist encouraged the intern to write the story.
22. The governor encouraged the businesswoman to meet the mayor.
23. The nephew persuaded the aunt to borrow the money.
24. The salesman induced the assistant to leave the company.
25. The executive ordered the employee to balance the budget.
26. The professor allowed the associate to teach the course.
27. The writer urged the lector to edit the novel.
28. The woman hired the nanny to prepare the meal.
29. The mother permitted the husband to adopt the child.
30. The senator bribed the traitor to sell the secrets.
31. The scientist selected the student to win the prize.
32. The doctor implored the co-worker to perform the surgery.
33. The baker trusted the chef to bake the cake.
34. The lawyer selected the partner to take the case.
35. The grandmother implored the husband to buy the presents.
36. The policeman ordered the constable to arrest the man.
37. The teacher bribed the janitor to steal the money.
38. The student forced the classmate to do the assignment.
39. The senator encouraged the colleague to run for president.
40. The waitress forced the barkeeper to help the passenger.
41. The writer allowed the secretary to write the book.
42. The ballerina invited the actress to perform the dance.
43. The butler bribed the daughter to unlock the safe.
44. The electrician hired the help to repair the furnace.
45. The athlete induced the representative to sign the contract.
46. The politician invited the candidate to give a speech.
47. The librarian trusted the user to buy the books.
48. The photographer persuaded the assistant to take the pictures.
49. The artist implored the manager to sell the painting.
50. The philanthropist encouraged the magnate to donate the money.
51. The banker bribed the custodian to steal the money.
52. The worker permitted the apprentice to go on vacation.

53. The judge ordered the prosecutor to stop the trial.
54. The student selected the promoter to organize the party.
55. The soldier bribed the guard to leave his post.
56. The quarterback forced the defender to throw the ball.
57. The politician urged the neighbour to run for office.
58. The secretary trusted the trainee to write the letter.
59. The activist invited the supporter to address the audience.
60. The accountant advised the intern to balance the books.
61. The pilot induced the co-pilot to fly the plane.
62. The actress implored the substitute to learn her lines.
63. The teacher urged the psychologist to help the child.
64. The shopper encouraged the friend to buy the coat.
65. The child trusted the nanny to clean his room.
66. The musician invited the drummer to sing the song.
67. The sheriff ordered the deputy to arrest the man.
68. The doctor selected the specialist to perform the operation.
69. The chairman persuaded the manager to answer the question.
70. The detective hired the help to follow the suspect.
71. The criminal permitted the cellmate to escape from jail.
72. The golfer advised the athlete to play for money.
73. The student allowed the roommate to have a party.
74. The writer advised the manager to sign the contract.
75. The cannibal forced the victim to eat the minister.
76. The athlete permitted the substitute to play the game.
77. The diplomat allowed the assistant to discuss the treaty.
78. The scientist hired the student to conduct the experiment.
79. The soldier ordered the private to push the button.
80. The secretary permitted the apprentice to attend the meeting.
81. The carpenter persuaded the trainee to build the table.
82. The executive implored the board to sell the company.
83. The mistress trusted the lover to keep the secret.
84. The astronomer urged the fiancé to watch the comet.
85. The gangster selected the accomplice to plan the robbery.
86. The astronaut allowed the spacetourist to touch the moon.
87. The refugee encouraged the child to cross the border.
88. The monk forced the soldier to pray for peace.
89. The schoolgirl invited the classmate to recite the poem.
90. The dancer encouraged the actor to join the ballet.
91. The pilot induced the co-pilot to change his course.
92. The dentist advised the manager to buy new equipment.
93. The zookeeper urged the instructor to train the animals.
94. The milkman implored the colleague to deliver the milk.
95. The umpire bribed the player to change his mind.
96. The accountant forced the trainee to erase the numbers.
97. The photographer persuaded the bride to take the portrait.
98. The nurse ordered the apprentice to empty the bedpans.
99. The hunter selected the dog to track the moose.
100. The waitress hired the teenager to serve the banquet.
101. The guard induced the soldier to free the prisoners.
102. The embezzler trusted the agitator to confess his crime.
103. The salesman ordered the subordinate to sell bad cars.
104. The musician allowed the youth to join the orchestra.
105. The referee permitted the assistant to make the decision.
106. The sailor ordered the private to fire the torpedo.

107. The librarian invited the researcher to give a speech.
108. The runner urged the friend to finish the race.
109. The driver implored the policewoman to stop the taxi.
110. The manager encouraged the administrator to hire the man.
111. The electrician hired the help to install the light.
112. The prisoner permitted the cellmate to escape from jail.
113. The coach induced the opponent to forfeit the game.
114. The repairman trusted the apprentice to fix the television.
115. The model persuaded the co-worker to pose without clothes.
116. The engineer selected the company to build the bridge.
117. The chauffeur allowed the nephew to drive the limousine.
118. The protestor forced the activist to stop the march.
119. The spy advised the confederate to tell the truth.
120. The florist invited the colleague to decorate the church.

Table S2: Sentence stimuli of the TVRR condition.

1. The broker persuaded to conceal the transaction was sent to jail.
2. The man hired to help the store was fired for theft.
3. The doctor implored to see the patient had left the hospital.
4. The reporter selected to get the story was given a raise.
5. The woman advised to see the play was leaving the theater.
6. The senator forced to chair the committee was sent the money.
7. The judge advised to sentence the defendant was reluctant to proceed.
8. The tailor hired to fix the suit had repaired the rip.
9. The swimmer urged to lose weight was beginning a diet.
10. The general permitted to leave the army had received an award.
11. The minister invited to give the sermon was about to arrive.
12. The mechanic trusted to repair the car had quit his job.
13. The singer allowed to perform the opera was past her prime.
14. The teacher urged to improve his teaching had taken a vacation.
15. The burglar induced to rob the bank was caught red handed.
16. The policeman ordered to watch the bank had caught the thieves.
17. The dentist invited to meet the actress was nervous last night.
18. The janitor persuaded to fix the faucet had botched the job.
19. The nurse induced to leave the patient was reprimanded very severely.
20. The prince advised to marry the princess had proposed last night.
21. The journalist encouraged to write the story had missed the deadline.
22. The governor encouraged to meet the mayor was running for reelection.
23. The nephew persuaded to borrow the money was in substantial debt.
24. The salesman induced to leave the company was known for dishonesty.
25. The executive ordered to balance the budget was fired for incompetence.
26. The professor allowed to teach the course was preparing his lectures.
27. The writer urged to edit the novel had requested more money.
28. The woman hired to prepare the meal had burned the meat.
29. The mother permitted to adopt the child had told her husband.
30. The senator bribed to sell the secrets was arrested for espionage.
31. The scientist selected to win the prize had arrived by plane.
32. The doctor implored to perform the surgery had left the country.
33. The baker trusted to bake the cake had won many awards.
34. The lawyer selected to take the case was very highly regarded.
35. The grandmother implored to buy the presents had forgotten her purse.
36. The policeman ordered to arrest the man had hurt his hand.
37. The teacher bribed to steal the money was fined for incompetence.
38. The student forced to do the assignment was failing the course.
39. The senator encouraged to run for president had written the article.
40. The waitress forced to help the passenger was ready to quit.
41. The writer allowed to write the book had received an advance.
42. The ballerina invited to perform the dance was practicing every day.
43. The butler bribed to unlock the safe was caught last night.
44. The electrician hired to repair the furnace had finished the job.
45. The athlete induced to sign the contract had injured his leg.
46. The politician invited to give a speech was given an award.
47. The librarian trusted to buy the books had completed the purchases.
48. The photographer persuaded to take the pictures had loaded the camera.
49. The artist implored to sell the painting had moved to Chicago.
50. The philanthropist encouraged to donate the money was eager to help.
51. The banker bribed to steal the money had moved to Australia.
52. The worker permitted to go on vacation was given a raise.

53. The judge ordered to stop the trial was asked to resign.
54. The student selected to organize the party had begun the preparations.
55. The soldier bribed to leave his post was reprimanded last week.
56. The quarterback forced to throw the ball was intercepted three times.
57. The politician urged to run for office was meeting with voters.
58. The secretary trusted to write the letter was given a raise.
59. The activitist invited to address the audience had prepared all night.
60. The accountant advised to balance the books had discovered an error.
61. The pilot induced to fly the plane had boarded the plane.
62. The actress implored to learn her lines was ready to quit.
63. The teacher urged to help the child had prepared the lesson.
64. The shopper encouraged to buy the coat was given a discount.
65. The child trusted to clean his room had fallen asleep instead.
66. The musician invited to sing the song was not very good.
67. The sheriff ordered to arrest the man had been tricked again.
68. The doctor selected to perform the operation was ready to begin again.
69. The chairman persuaded to answer the question was preparing to resign.
70. The detective hired to follow the suspect had lost the trail.
71. The criminal permitted to escape from jail was captured last night.
72. The golfer advised to play for money had lost the match.
73. The student allowed to have a party had bought new albums.
74. The writer advised to sign the contract was given a deadline.
75. The cannibal forced to eat the minister had set the table.
76. The athlete permitted to play the game was injured last time.
77. The diplomat allowed to discuss the treaty was threatened this morning.
78. The scientist hired to conduct the experiment was given a computer.
79. The soldier ordered to push the button had closed his eyes.
80. The secretary permitted to attend the meeting had made the coffee.
81. The carpenter persuaded to build the table had been given money.
82. The executive implored to sell the company was fired last week.
83. The mistress trusted to keep the secret had threatened her lover.
84. The astronomer urged to watch the comet was too busy reading.
85. The gangster selected to plan the robbery had no experience stealing.
86. The astronaut allowed to touch the moon was wearing a helmet.
87. The refugee encouraged to cross the border had bribed the guard.
88. The monk forced to pray for peace was secretly a soldier.
89. The schoolgirl invited to recite the poem had been very nervous.
90. The dancer encouraged to join the ballet was talented in music.
91. The pilot induced to change his course had seen the gun.
92. The dentist advised to buy new equipment was sued by patients.
93. The zookeeper urged to train the animals had seen the circus.
94. The milkman implored to deliver the milk was afraid of dogs.
95. The umpire bribed to change his mind had been warned before.
96. The accountant forced to erase the numbers was arrested last week.
97. The photographer persuaded to take the portrait had disliked the model.
98. The nurse ordered to empty the bedpans was given a raise.
99. The hunter selected to track the moose had sold his gun.
100. The waitress hired to serve the banquet was wearing a uniform.
101. The guard induced to free the prisoners had been paid off.
102. The embezzler trusted to confess his crime was leaving the country.
103. The salesman ordered to sell bad cars had a guilty conscience.
104. The musician allowed to join the orchestra was not very good.
105. The referee permitted to make the decision had to think fast.
106. The sailor ordered to fire the torpedo was afraid of war.

107. The librarian invited to give a speech had drunk too much.
108. The runner urged to finish the race was awarded a medal.
109. The driver implored to stop the taxi had threatened the passenger.
110. The manager encouraged to hire the man was impressed by him.
111. The electrician hired to install the light had a good reputation.
112. The prisoner permitted to escape from jail was shot this morning.
113. The coach induced to forfeit the game had several sick players.
114. The repairman trusted to fix the television was selling bad parts.
115. The model persuaded to pose without clothes had to sign papers.
116. The engineer selected to build the bridge was paid very well.
117. The chauffeur allowed to drive the limousine had wrecked two cars.
118. The protestor forced to stop the march was threatened by police.
119. The spy advised to tell the truth had lied for years.
120. The florist invited to decorate the church was good at weddings.

Table S3: Sentence stimuli of the IVWR condition.

1. The broker planned to conceal the transaction was sent to jail.
2. The man intended to help the store was fired for theft.
3. The doctor agreed to see the patient had left the hospital.
4. The reporter struggled to get the story was given a raise.
5. The woman agreed to see the play was leaving the theater.
6. The senator attempted to chair the committee was sent the money.
7. The judge hoped to sentence the defendant was reluctant to proceed.
8. The tailor began to fix the suit had repaired the rip.
9. The swimmer decided to lose weight was beginning a diet.
10. The general refused to leave the army had received an award.
11. The minister started to give the sermon was about to arrive.
12. The mechanic refused to repair the car had quit his job.
13. The singer decided to perform the opera was past her prime.
14. The teacher planned to improve his teaching had taken a vacation.
15. The burglar schemed to rob the bank was caught red handed.
16. The policeman intended to watch the bank had caught the thieves.
17. The dentist hoped to meet the actress was nervous last night.
18. The janitor tried to fix the faucet had botched the job.
19. The nurse hesitated to leave the patient was reprimanded very severely.
20. The prince yearned to marry the princess had proposed last night.
21. The journalist attempted to write the story had missed the deadline.
22. The governor hoped to meet the mayor was running for reelection.
23. The nephew hesitated to borrow the money was in substantial debt.
24. The salesman tried to leave the company was known for dishonesty.
25. The executive planned to balance the budget was fired for incompetence.
26. The professor tried to teach the course was preparing his lectures.
27. The writer decided to edit the novel had requested more money.
28. The woman struggled to prepare the meal had burned the meat.
29. The mother agreed to adopt the child had told her husband.
30. The senator schemed to sell the secrets was arrested for espionage.
31. The scientist aspired to win the prize had arrived by plane.
32. The doctor began to perform the surgery had left the country.
33. The baker started to bake the cake had won many awards.
34. The lawyer declined to take the case was very highly regarded.
35. The grandmother intended to buy the presents had forgotten her purse.
36. The policeman struggled to arrest the man had hurt his hand.
37. The teacher schemed to steal the money was fined for incompetence.
38. The student hesitated to do the assignment was failing the course.
39. The senator aspired to run for president had written the article.
40. The waitress refused to help the passenger was ready to quit.
41. The writer started to write the book had received an advance.
42. The ballerina aspired to perform the dance was practicing every day.
43. The butler schemed to unlock the safe was caught last night.
44. The electrician attempted to repair the furnace had finished the job.
45. The athlete hoped to sign the contract had injured his leg.
46. The politician began to give a speech was given an award.
47. The librarian decided to buy the books had completed the purchases.
48. The photographer began to take the pictures had loaded the camera.
49. The artist declined to sell the painting had moved to Chicago.
50. The philanthropist intended to donate the money was eager to help.
51. The banker planned to steal the money had moved to Australia.
52. The worker refused to go on vacation was given a raise.

53. The judge tried to stop the trial was asked to resign.
54. The student aspired to organize the party had begun the preparations.
55. The soldier schemed to leave his post was reprimanded last week.
56. The quarterback tried to throw the ball was intercepted three times.
57. The politician decided to run for office was meeting with voters.
58. The secretary started to write the letter was given a raise.
59. The activist hoped to address the audience had prepared all night.
60. The accountant attempted to balance the books had discovered an error.
61. The pilot hesitated to fly the plane had boarded the plane.
62. The actress struggled to learn her lines was ready to quit.
63. The teacher began to help the child had prepared the lesson.
64. The shopper declined to buy the coat was given a discount.
65. The child agreed to clean his room had fallen asleep instead.
66. The musician started to sing the song was not very good.
67. The sheriff attempted to arrest the man had been tricked again.
68. The doctor agreed to perform the operation was ready to begin again.
69. The chairman refused to answer the question was preparing to resign.
70. The detective planned to follow the suspect had lost the trail.
71. The criminal struggled to escape from jail was captured last night.
72. The golfer aspired to play for money had lost the match.
73. The student intended to have a party had bought new albums.
74. The writer agreed to sign the contract was given a deadline.
75. The cannibal declined to eat the minister had set the table.
76. The athlete began to play the game was injured last time.
77. The diplomat hesitated to discuss the treaty was threatened this morning.
78. The scientist hoped to conduct the experiment was given a computer.
79. The soldier refused to push the button had closed his eyes.
80. The secretary planned to attend the meeting had made the coffee.
81. The carpenter intended to build the table had been given money.
82. The executive declined to sell the company was fired last week.
83. The mistress struggled to keep the secret had threatened her lover.
84. The astronomer attempted to watch the comet was too busy reading.
85. The gangster began to plan the robbery had no experience stealing.
86. The astronaut aspired to touch the moon was wearing a helmet.
87. The refugee schemed to cross the border had bribed the guard.
88. The monk decided to pray for peace was secretly a soldier.
89. The schoolgirl decided to recite the poem had been very nervous.
90. The dancer tried to join the ballet was talented in music.
91. The pilot hesitated to change his course had seen the gun.
92. The dentist planned to buy new equipment was sued by patients.
93. The zookeeper intended to train the animals had seen the circus.
94. The milkman agreed to deliver the milk was afraid of dogs.
95. The umpire refused to change his mind had been warned before.
96. The accountant declined to erase the numbers was arrested last week.
97. The photographer attempted to take the portrait had disliked the model.
98. The nurse struggled to empty the bedpans was given a raise.
99. The hunter began to track the moose had sold his gun.
100. The waitress hoped to serve the banquet was wearing a uniform.
101. The guard decided to free the prisoners had been paid off.
102. The embezzler started to confess his crime was leaving the country.
103. The salesman schemed to sell bad cars had a guilty conscience.
104. The musician aspired to join the orchestra was not very good.
105. The referee tried to make the decision had to think fast.
106. The sailor hesitated to fire the torpedo was afraid of war.

107. The librarian agreed to give a speech had drunk too much.
108. The runner struggled to finish the race was awarded a medal.
109. The driver refused to stop the taxi had threatened the passenger.
110. The manager decided to hire the man was impressed by him.
111. The electrician tried to install the light had a good reputation.
112. The prisoner schemed to escape from jail was shot this morning.
113. The coach planned to forfeit the game had several sick players.
114. The repairman began to fix the television was selling bad parts.
115. The model declined to pose without clothes had to sign papers.
116. The engineer intended to build the bridge was paid very well.
117. The chauffeur aspired to drive the limousine had wrecked two cars.
118. The protestor hesitated to stop the march was threatened by police.
119. The spy started to tell the truth had lied for years.
120. The florist hoped to decorate the church was good at weddings.

Table S4: Sentence stimuli of the IVCO condition.

| |
|---|
| 1. The broker planned to conceal the transaction. |
| 2. The man intended to help the store. |
| 3. The doctor agreed to see the patient. |
| 4. The reporter struggled to get the story. |
| 5. The woman agreed to see the play. |
| 6. The senator attempted to chair the committee. |
| 7. The judge hoped to sentence the defendant. |
| 8. The tailor began to fix the suit. |
| 9. The swimmer decided to lose weight. |
| 10. The general refused to leave the army. |
| 11. The minister started to give the sermon. |
| 12. The mechanic refused to repair the car. |
| 13. The singer decided to perform the opera. |
| 14. The teacher planned to improve his teaching. |
| 15. The burglar schemed to rob the bank. |
| 16. The policeman intended to watch the bank. |
| 17. The dentist hoped to meet the actress. |
| 18. The janitor tried to fix the faucet. |
| 19. The nurse hesitated to leave the patient. |
| 20. The prince yearned to marry the princess. |
| 21. The journalist attempted to write the story. |
| 22. The governor hoped to meet the mayor. |
| 23. The nephew hesitated to borrow the money. |
| 24. The salesman tried to leave the company. |
| 25. The executive planned to balance the budget. |
| 26. The professor tried to teach the course. |
| 27. The writer decided to edit the novel. |
| 28. The woman struggled to prepare the meal. |
| 29. The mother agreed to adopt the child. |
| 30. The senator schemed to sell the secrets. |
| 31. The scientist aspired to win the prize. |
| 32. The doctor began to perform the surgery. |
| 33. The baker started to bake the cake. |
| 34. The lawyer declined to take the case. |
| 35. The grandmother intended to buy the presents. |
| 36. The policeman struggled to arrest the man. |
| 37. The teacher schemed to steal the money. |
| 38. The student hesitated to do the assignment. |
| 39. The senator aspired to run for president. |
| 40. The waitress refused to help the passenger. |
| 41. The writer started to write the book. |
| 42. The ballerina aspired to perform the dance. |
| 43. The butler schemed to unlock the safe. |
| 44. The electrician attempted to repair the furnace. |
| 45. The athlete hoped to sign the contract. |
| 46. The politician began to give a speech. |
| 47. The librarian decided to buy the books. |
| 48. The photographer began to take the pictures. |
| 49. The artist declined to sell the painting. |
| 50. The philanthropist intended to donate the money. |
| 51. The banker planned to steal the money. |
| 52. The worker refused to go on vacation. |

53. The judge tried to stop the trial.
54. The student aspired to organize the party.
55. The soldier schemed to leave his post.
56. The quarterback tried to throw the ball.
57. The politician decided to run for office.
58. The secretary started to write the letter.
59. The activitist hoped to address the audience.
60. The accountant attempted to balance the books.
61. The pilot hesitated to fly the plane.
62. The actress struggled to learn her lines.
63. The teacher began to help the child.
64. The shopper declined to buy the coat.
65. The child agreed to clean his room.
66. The musician started to sing the song.
67. The sheriff attempted to arrest the man.
68. The doctor agreed to perform the operation.
69. The chairman refused to answer the question.
70. The detective planned to follow the suspect.
71. The criminal struggled to escape from jail.
72. The golfer aspired to play for money.
73. The student intended to have a party.
74. The writer agreed to sign the contract.
75. The cannibal declined to eat the minister.
76. The athlete began to play the game.
77. The diplomat hesitated to discuss the treaty.
78. The scientist hoped to conduct the experiment.
79. The soldier refused to push the button.
80. The secretary planned to attend the meeting.
81. The carpenter intended to build the table.
82. The executive declined to sell the company.
83. The mistress struggled to keep the secret.
84. The astronomer attempted to watch the comet.
85. The gangster began to plan the robbery.
86. The astronaut aspired to touch the moon.
87. The refugee schemed to cross the border.
88. The monk decided to pray for peace.
89. The schoolgirl decided to recite the poem.
90. The dancer tried to join the ballet.
91. The pilot hesitated to change his course.
92. The dentist planned to buy new equipment.
93. The zookeeper intended to train the animals.
94. The milkman agreed to deliver the milk.
95. The umpire refused to change his mind.
96. The accountant declined to erase the numbers.
97. The photographer attempted to take the portrait.
98. The nurse struggled to empty the bedpans.
99. The hunter began to track the moose.
100. The waitress hoped to serve the banquet.
101. The guard decided to free the prisoners.
102. The embezzler started to confess his crime.
103. The salesman schemed to sell bad cars.
104. The musician aspired to join the orchestra.
105. The referee tried to make the decision.
106. The sailor hesitated to fire the torpedo.

107. The librarian agreed to give a speech.
108. The runner struggled to finish the race.
109. The driver refused to stop the taxi.
110. The manager decided to hire the man.
111. The electrician tried to install the light.
112. The prisoner schemed to escape from jail.
113. The coach planned to forfeit the game.
114. The repairman began to fix the television.
115. The model declined to pose without clothes.
116. The engineer intended to build the bridge.
117. The chauffeur aspired to drive the limousine.
118. The protestor hesitated to stop the march.
119. The spy started to tell the truth.
120. The florist hoped to decorate the church.

Table S5: Sentence stimuli of the SRAI condition.

| |
|---|
| 1. The musician that witnessed the accident angered the policeman a lot. |
| 2. The contestant that misplaced the prize made a big impression on Mary. |
| 3. The cowboy that carried the pistol was known to be unreliable. |
| 4. The scientist that studied the climate did not interest the reporter. |
| 5. The director that watched the movie received a prize at the film festival. |
| 6. The student that attended the school was visited by the governor. |
| 7. The teacher that watched the play upset a few of the students. |
| 8. The woman that reported the accident caused a number of serious injuries. |
| 9. The plumber that dropped the wrench was found near the back door. |
| 10. The banker that refused the loan created a problem for the mayor. |
| 11. The lawyer that reviewed the trial was covered by the national media. |
| 12. The psychologist that printed the notes got lost somewhere in the basement. |
| 13. The child that loaded the revolver injured the teenage babysitter. |
| 14. The golfer that mastered the game was ignored by most sportswriters. |
| 15. The salesman that examined the product was mentioned in the newsletter. |
| 16. The fireman that fought the fire caused only a small amount of damage. |
| 17. The fish that attacked the lure impressed the fisherman quite a lot. |
| 18. The farmer that purchased the tractor arrived at the store late last night. |
| 19. The gardener that trimmed the plants helped make the house more attractive. |
| 20. The pilot that crashed the plane was grounded by the safety board. |
| 21. The elephant that drank the water was located in the heart of Africa. |
| 22. The actor that rehearsed the play was given first prize at the awards dinner. |
| 23. The student that practiced the instrument had been around for a few months. |
| 24. The spy that encoded the message was smuggled out of the country in a crate. |
| 25. The journalist that composed the article caused a big scandal. |
| 26. The diner that consumed the meat came from a distant foreign country. |
| 27. The woman that coveted the jewelry didn't do anything for the old man. |
| 28. The dieter that desired the dessert was not very healthy, Susan said. |
| 29. The man that sailed the sea was very agitated that day. |
| 30. The burglar that stole the necklace was famous throughout the country. |

Table S6: Sentence stimuli of the SRIA condition.

1. The accident that terrified the musician angered the policeman a lot.
2. The prize that delighted the contestant made a big impression on Mary.
3. The pistol that injured the cowboy was known to be unreliable.
4. The climate that annoyed the scientist did not interest the reporter.
5. The movie that pleased the director received a prize at the film festival.
6. The school that educated the student was visited by the governor.
7. The play that angered the teacher upset a few of the students.
8. The accident that bothered the woman caused a number of serious injuries.
9. The wrench that bruised the plumber was found near the back door.
10. The loan that worried the banker created a problem for the mayor.
11. The trial that confused the lawyer was covered by the national media.
12. The notes that annoyed the psychologist got lost somewhere in the basement.
13. The revolver that scared the child injured the teenage babysitter.
14. The game that excited the golfer was ignored by most sportswriters.
15. The product that excited the salesman was mentioned in the newsletter.
16. The fire that burned the fireman caused only a small amount of damage.
17. The lure that attracted the fish impressed the fisherman quite a lot.
18. The tractor that impressed the farmer arrived at the store late last night.
19. The plants that pleased the gardener helped make the house more attractive.
20. The plane that worried the pilot was grounded by the safety board.
21. The water that cooled the elephant was located in the heart of Africa.
22. The play that delighted the actor was given first prize at the awards dinner.
23. The instrument that frustrated the student had been around for a few months.
24. The message that alarmed the spy was smuggled out of the country in a crate.
25. The article that bothered the journalist caused a big scandal.
26. The meat that satisfied the diner came from a distant foreign country.
27. The jewelry that dazzled the woman didn't do anything for the old man.
28. The dessert that tempted the dieter was not very healthy, Susan said.
29. The sea that troubled the man was very agitated that day.
30. The necklace that fascinated the burglar was famous throughout the country.

Table S7: Sentence stimuli of the ORAI condition.

| |
|---|
| 1. The musician that the accident terrified angered the policeman a lot. |
| 2. The contestant that the prize delighted made a big impression on Mary. |
| 3. The cowboy that the pistol injured was known to be unreliable. |
| 4. The scientist that the climate annoyed did not interest the reporter. |
| 5. The director that the movie pleased received a prize at the film festival. |
| 6. The student that the school educated was visited by the governor. |
| 7. The teacher that the play angered upset a few of the students. |
| 8. The woman that the accident bothered caused a number of serious injuries. |
| 9. The plumber that the wrench bruised was found near the back door. |
| 10. The banker that the loan worried created a problem for the mayor. |
| 11. The lawyer that the trial confused was covered by the national media. |
| 12. The psychologist that the notes annoyed got lost somewhere in the basement. |
| 13. The child that the revolver scared injured the teenage babysitter. |
| 14. The golfer that the game excited was ignored by most sportswriters. |
| 15. The salesman that the product excited was mentioned in the newsletter. |
| 16. The fireman that the fire burned caused only a small amount of damage. |
| 17. The fish that the lure attracted impressed the fisherman quite a lot. |
| 18. The farmer that the tractor impressed arrived at the store late last night. |
| 19. The gardener that the plants pleased helped make the house more attractive. |
| 20. The pilot that the plane worried was grounded by the safety board. |
| 21. The elephant that the water cooled was located in the heart of Africa. |
| 22. The actor that the play delighted was given first prize at the awards dinner. |
| 23. The student that the instrument frustrated had been around for a few months. |
| 24. The spy that the message alarmed was smuggled out of the country in a crate. |
| 25. The journalist that the article bothered caused a big scandal. |
| 26. The diner that the meat satisfied came from a distant foreign country. |
| 27. The woman that the jewelry dazzled didn't do anything for the old man. |
| 28. The dieter that the dessert tempted was not very healthy, Susan said. |
| 29. The man that the sea troubled was very agitated that day. |
| 30. The burglar that the necklace fascinated was famous throughout the country. |

Table S8: Sentence stimuli of the ORIA condition.

1. The accident that the musician witnessed angered the policeman a lot.
2. The prize that the contestant misplaced made a big impression on Mary.
3. The pistol that the cowboy carried was known to be unreliable.
4. The climate that the scientist studied did not interest the reporter.
5. The movie that the director watched received a prize at the film festival.
6. The school that the student attended was visited by the governor.
7. The play that the teacher watched upset a few of the students.
8. The accident that the woman reported caused a number of serious injuries.
9. The wrench that the plumber dropped was found near the back door.
10. The loan that the banker refused created a problem for the mayor.
11. The trial that the lawyer confused was covered by the national media.
12. The notes that the psychologist printed got lost somewhere in the basement.
13. The revolver that the child loaded injured the teenage babysitter.
14. The game that the golfer mastered was ignored by most sportswriters.
15. The product that the salesman examined was mentioned in the newsletter.
16. The fire that the fireman fought caused only a small amount of damage.
17. The lure that the fish attacked impressed the fisherman quite a lot.
18. The tractor that the farmer purchased arrived at the store late last night.
19. The plants that the gardener trimmed helped make the house more attractive.
20. The plane that the pilot crashed was grounded by the safety board.
21. The water that the elephant drank was located in the heart of Africa.
22. The play that the actor rehearsed was given first prize at the awards dinner.
23. The instrument that the student practiced had been around for a few months.
24. The message that the spy encoded was smuggled out of the country in a crate.
25. The article that the journalist composed caused a big scandal.
26. The meat that the diner consumed came from a distant foreign country.
27. The jewelry that the woman coveted didn't do anything for the old man.
28. The dessert that the dieter desired was not very healthy, Susan said.
29. The sea that the man sailed was very agitated that day.
30. The necklace that the burglar stole was famous throughout the country.

Table S9: Summary of Linear Mixed Model for the P600 amplitude predicted by condition (TVRR vs. TVDO), laterality, and sagittality

| | Estimate | Std. Error | df | t value | Pr(>\|t\|) | |
|---|---|---|---|---|---|---|
| (Intercept) | -0.03 | 0.34 | 51.77 | -0.09 | 0.929 | |
| prestim | 0.27 | 0.02 | 4335.50 | 15.75 | <0.001 | *** |
| condition: TVRR | 0.09 | 0.40 | 68.08 | 0.22 | 0.826 | |
| lat: [S.medial] | -0.13 | 0.14 | 4290.08 | -0.93 | 0.351 | |
| lat: [S.right] | -0.27 | 0.14 | 4291.00 | -1.97 | 0.049 | * |
| sag: [S.posterior] | -0.70 | 0.10 | 4303.40 | -6.99 | <0.001 | *** |
| prestim:conditionTVRR | -0.01 | 0.02 | 4337.87 | -0.43 | 0.666 | |
| condition: TVRR:lat[S.medial] | 0.19 | 0.20 | 4289.73 | 0.98 | 0.328 | |
| condition: TVRR:lat[S.right] | -0.06 | 0.20 | 4290.62 | -0.31 | 0.759 | |
| condition: TVRR:sag[S.posterior] | 0.19 | 0.14 | 4299.00 | 1.33 | 0.183 | |
| lat[S.medial]:sag[S.posterior] | -0.20 | 0.14 | 4289.37 | -1.44 | 0.15 | |
| lat[S.right]:sag[S.posterior] | 0.07 | 0.14 | 4289.37 | 0.51 | 0.613 | |
| condition: TVRR:lat[S.medial]:sag[S.posterior] | 0.07 | 0.20 | 4289.41 | 0.34 | 0.736 | |
| condition: TVRR:lat[S.right]:sag[S.posterior] | 0.08 | 0.20 | 4289.42 | 0.42 | 0.673 | |

Table S10: Summary of Linear Mixed Model for the P600 amplitude predicted by condition (TVRR vs. TVDO), laterality, sagittality, and age

| | Estimate | Std. Error | df | t value | Pr(>\|t\|) | |
|---|---|---|---|---|---|---|
| (Intercept) | -0.06 | 0.34 | 51.61 | -0.17 | 0.868 | |
| prestim | 0.27 | 0.02 | 4326.65 | 15.75 | <0.001 | *** |
| condition: TVRR | 0.15 | 0.39 | 69.19 | 0.38 | 0.707 | |
| lat: [S.medial] | -0.12 | 0.14 | 4290.07 | -0.87 | 0.383 | |
| lat: [S.right] | -0.29 | 0.14 | 4291.02 | -2.03 | 0.042 | * |
| sag: [S.posterior] | -0.68 | 0.10 | 4303.17 | -6.67 | <0.001 | *** |
| age_z | -0.17 | 0.31 | 26.92 | -0.57 | 0.574 | |
| prestim:conditionTVRR | -0.01 | 0.02 | 4322.47 | -0.42 | 0.674 | |
| condition: TVRR:lat[S.medial] | 0.17 | 0.20 | 4289.73 | 0.84 | 0.403 | |
| condition: TVRR:lat[S.right] | -0.04 | 0.20 | 4290.54 | -0.21 | 0.836 | |
| condition: TVRR:sag[S.posterior] | 0.15 | 0.14 | 4298.83 | 1.03 | 0.302 | |
| lat[S.medial]:sag[S.posterior] | -0.20 | 0.14 | 4289.39 | -1.39 | 0.165 | |
| lat[S.right]:sag[S.posterior] | 0.08 | 0.14 | 4289.39 | 0.54 | 0.592 | |
| condition: TVRR:age_z | 0.40 | 0.32 | 27.87 | 1.24 | 0.224 | |
| lat[S.medial]:age_z | 0.05 | 0.15 | 4289.38 | 0.30 | 0.761 | |
| lat[S.right]:age_z | -0.08 | 0.15 | 4289.38 | -0.55 | 0.581 | |
| sag[S.posterior]:age_z | 0.16 | 0.11 | 4289.38 | 1.53 | 0.125 | |
| condition: TVRR:lat[S.medial]:sag[S.posterior] | 0.07 | 0.20 | 4289.43 | 0.33 | 0.744 | |
| condition: TVRR:lat[S.right]:sag[S.posterior] | 0.08 | 0.20 | 4289.44 | 0.41 | 0.685 | |
| condition: TVRR:lat[S.medial]:age_z | -0.15 | 0.21 | 4289.39 | -0.69 | 0.489 | |
| condition: TVRR:lat[S.right]:age_z | 0.12 | 0.21 | 4289.54 | 0.56 | 0.576 | |
| condition: TVRR:sag[S.posterior]:age_z | -0.25 | 0.15 | 4289.38 | -1.66 | 0.098 | . |
| lat[S.medial]:sag[S.posterior]:age_z | 0.03 | 0.15 | 4289.38 | 0.22 | 0.826 | |
| lat[S.right]:sag[S.posterior]:age_z | 0.03 | 0.15 | 4289.39 | 0.23 | 0.821 | |
| condition: TVRR:lat[S.medial]:sag[S.posterior]:age_z | -0.01 | 0.21 | 4289.38 | -0.05 | 0.959 | |
| condition: TVRR:lat[S.right]:sag[S.posterior]:age_z | -0.02 | 0.21 | 4289.39 | -0.08 | 0.938 | |

Table S11: Summary of Linear Mixed Model for the P600 amplitude predicted by condition (TVRR vs. TVDO), laterality, sagittality, and PTA

| | Estimate | Std. Error | df | t value | Pr(>\|t\|) | |
|---|---|---|---|---|---|---|
| (Intercept) | -0.05 | 0.33 | 50.64 | -0.14 | 0.891 | |
| prestim | 0.27 | 0.02 | 4287.11 | 15.71 | <0.001 | *** |
| condition: TVRR | 0.10 | 0.38 | 73.14 | 0.27 | 0.784 | |
| lat: [S.medial] | -0.13 | 0.14 | 4289.90 | -0.93 | 0.351 | |
| lat: [S.right] | -0.28 | 0.14 | 4290.88 | -2.00 | 0.045 | * |
| sag: [S.posterior] | -0.70 | 0.10 | 4303.49 | -6.97 | <0.001 | *** |
| PTA_z | -0.33 | 0.27 | 25.83 | -1.20 | 0.243 | |
| att_resid_z | 0.37 | 0.27 | 27.48 | 1.36 | 0.185 | |
| prestim:conditionTVRR | -0.01 | 0.02 | 4270.71 | -0.38 | 0.703 | |
| condition: TVRR:lat[S.medial] | 0.19 | 0.20 | 4289.54 | 0.98 | 0.327 | |
| condition: TVRR:lat[S.right] | -0.06 | 0.20 | 4290.51 | -0.29 | 0.772 | |
| condition: TVRR:sag[S.posterior] | 0.19 | 0.14 | 4299.26 | 1.32 | 0.188 | |
| lat[S.medial]:sag[S.posterior] | -0.20 | 0.14 | 4289.18 | -1.44 | 0.149 | |
| lat[S.right]:sag[S.posterior] | 0.07 | 0.14 | 4289.18 | 0.50 | 0.619 | |
| condition: TVRR:PTA_z | 0.65 | 0.27 | 27.97 | 2.39 | 0.024 | * |
| lat[S.medial]:PTA_z | -0.01 | 0.14 | 4289.19 | -0.07 | 0.943 | |
| lat[S.right]:PTA_z | -0.11 | 0.14 | 4289.24 | -0.80 | 0.425 | |
| sag[S.posterior]:PTA_z | 0.08 | 0.10 | 4289.23 | 0.79 | 0.43 | |
| condition: TVRR:lat[S.medial]:sag[S.posterior] | 0.07 | 0.20 | 4289.22 | 0.35 | 0.729 | |
| condition: TVRR:lat[S.right]:sag[S.posterior] | 0.08 | 0.20 | 4289.24 | 0.42 | 0.674 | |
| condition: TVRR:lat[S.medial]:PTA_z | 0.03 | 0.19 | 4289.18 | 0.16 | 0.873 | |
| condition: TVRR:lat[S.right]:PTA_z | 0.06 | 0.19 | 4289.21 | 0.30 | 0.762 | |
| condition: TVRR:sag[S.posterior]:PTA_z | -0.09 | 0.14 | 4289.21 | -0.64 | 0.522 | |
| lat[S.medial]:sag[S.posterior]:PTA_z | -0.01 | 0.14 | 4289.17 | -0.07 | 0.94 | |
| lat[S.right]:sag[S.posterior]:PTA_z | -0.03 | 0.14 | 4289.18 | -0.22 | 0.822 | |
| condition: TVRR:lat[S.medial]:sag[S.posterior]:PTA_z | 0.07 | 0.19 | 4289.17 | 0.35 | 0.727 | |
| condition: TVRR:lat[S.right]:sag[S.posterior]:PTA_z | -0.02 | 0.19 | 4289.18 | -0.10 | 0.923 | |

Table S12: Summary of Linear Mixed Model for the P600 amplitude predicted by condition (TVRR vs. TVDO), laterality, sagittality, and RS

| | Estimate | Std. Error | df | t value | Pr(>\|t\|) | |
|---|---|---|---|---|---|---|
| (Intercept) | -0.02 | 0.32 | 55.62 | -0.07 | 0.944 | |
| prestim | 0.27 | 0.02 | 4220.55 | 15.88 | <0.001 | *** |
| condition: TVRR | 0.08 | 0.38 | 74.48 | 0.21 | 0.836 | |
| lat: [S.medial] | -0.13 | 0.14 | 4290.91 | -0.93 | 0.352 | |
| lat: [S.right] | -0.27 | 0.14 | 4291.91 | -1.96 | 0.05 | * |
| sag: [S.posterior] | -0.70 | 0.10 | 4305.48 | -6.97 | <0.001 | *** |
| RS_z | 0.56 | 0.26 | 26.56 | 2.15 | 0.041 | * |
| prestim:conditionTVRR | -0.01 | 0.02 | 4247.04 | -0.53 | 0.595 | |
| condition: TVRR:lat[S.medial] | 0.19 | 0.20 | 4290.52 | 0.97 | 0.33 | |
| condition: TVRR:lat[S.right] | -0.06 | 0.20 | 4291.49 | -0.31 | 0.756 | |
| condition: TVRR:sag[S.posterior] | 0.19 | 0.14 | 4300.66 | 1.31 | 0.19 | |
| lat[S.medial]:sag[S.posterior] | -0.20 | 0.14 | 4290.13 | -1.44 | 0.149 | |
| lat[S.right]:sag[S.posterior] | 0.07 | 0.14 | 4290.13 | 0.51 | 0.612 | |
| condition: TVRR:RS_z | -0.68 | 0.28 | 28.98 | -2.44 | 0.021 | * |
| lat[S.medial]:RS_z | 0.17 | 0.14 | 4290.23 | 1.24 | 0.216 | |
| lat[S.right]:RS_z | -0.03 | 0.14 | 4290.25 | -0.23 | 0.816 | |
| sag[S.posterior]:RS_z | 0.13 | 0.10 | 4292.29 | 1.32 | 0.186 | |
| condition: TVRR:lat[S.medial]:sag[S.posterior] | 0.07 | 0.20 | 4290.17 | 0.34 | 0.736 | |
| condition: TVRR:lat[S.right]:sag[S.posterior] | 0.08 | 0.20 | 4290.18 | 0.42 | 0.673 | |
| condition: TVRR:lat[S.medial]:RS_z | 0.00 | 0.20 | 4290.26 | 0.01 | 0.989 | |
| condition: TVRR:lat[S.right]:RS_z | -0.12 | 0.20 | 4290.20 | -0.58 | 0.56 | |
| condition: TVRR:sag[S.posterior]:RS_z | -0.13 | 0.14 | 4291.28 | -0.93 | 0.351 | |
| lat[S.medial]:sag[S.posterior]:RS_z | 0.02 | 0.14 | 4290.14 | 0.15 | 0.884 | |
| lat[S.right]:sag[S.posterior]:RS_z | 0.04 | 0.14 | 4290.12 | 0.32 | 0.75 | |
| condition: TVRR:lat[S.medial]:sag[S.posterior]:RS_z | -0.02 | 0.20 | 4290.13 | -0.11 | 0.909 | |
| condition: TVRR:lat[S.right]:sag[S.posterior]:RS_z | -0.06 | 0.20 | 4290.12 | -0.32 | 0.749 | |

Table S13: Summary of Linear Mixed Model for the P600 amplitude predicted by condition (TVRR vs. TVDO), laterality, sagittality, and OS

| | Estimate | Std. Error | df | t value | Pr(>\|t\|) | |
|---|---|---|---|---|---|---|
| (Intercept) | -0.04 | 0.34 | 51.89 | -0.13 | 0.895 | |
| prestim | 0.27 | 0.02 | 4317.79 | 15.76 | <0.001 | *** |
| condition: TVRR | 0.13 | 0.39 | 69.37 | 0.32 | 0.749 | |
| lat: [S.medial] | -0.14 | 0.14 | 4289.66 | -0.97 | 0.332 | |
| lat: [S.right] | -0.28 | 0.14 | 4290.74 | -1.96 | 0.051 | . |
| sag: [S.posterior] | -0.70 | 0.10 | 4302.27 | -6.95 | <0.001 | *** |
| OS_z | 0.16 | 0.30 | 27.67 | 0.52 | 0.604 | |
| prestim:conditionTVRR | -0.01 | 0.02 | 4311.90 | -0.50 | 0.618 | |
| condition: TVRR:lat[S.medial] | 0.20 | 0.20 | 4289.30 | 1.01 | 0.312 | |
| condition: TVRR:lat[S.right] | -0.04 | 0.20 | 4290.27 | -0.20 | 0.84 | |
| condition: TVRR:sag[S.posterior] | 0.21 | 0.14 | 4298.00 | 1.45 | 0.147 | |
| lat[S.medial]:sag[S.posterior] | -0.20 | 0.14 | 4288.94 | -1.42 | 0.156 | |
| lat[S.right]:sag[S.posterior] | 0.07 | 0.14 | 4288.93 | 0.51 | 0.607 | |
| condition: TVRR:OS_z | -0.42 | 0.31 | 28.83 | -1.35 | 0.188 | |
| lat[S.medial]:OS_z | 0.05 | 0.15 | 4288.93 | 0.36 | 0.718 | |
| lat[S.right]:OS_z | 0.00 | 0.15 | 4289.19 | 0.01 | 0.989 | |
| sag[S.posterior]:OS_z | -0.00 | 0.10 | 4290.03 | -0.02 | 0.982 | |
| condition: TVRR:lat[S.medial]:sag[S.posterior] | 0.06 | 0.20 | 4288.98 | 0.31 | 0.757 | |
| condition: TVRR:lat[S.right]:sag[S.posterior] | 0.08 | 0.20 | 4288.99 | 0.42 | 0.675 | |
| condition: TVRR:lat[S.medial]:OS_z | -0.07 | 0.21 | 4288.94 | -0.32 | 0.745 | |
| condition: TVRR:lat[S.right]:OS_z | -0.19 | 0.21 | 4289.06 | -0.91 | 0.362 | |
| condition: TVRR:sag[S.posterior]:OS_z | -0.17 | 0.15 | 4290.18 | -1.15 | 0.249 | |
| lat[S.medial]:sag[S.posterior]:OS_z | -0.02 | 0.15 | 4288.95 | -0.11 | 0.915 | |
| lat[S.right]:sag[S.posterior]:OS_z | -0.01 | 0.15 | 4288.92 | -0.10 | 0.923 | |
| condition: TVRR:lat[S.medial]:sag[S.posterior]:OS_z | 0.05 | 0.21 | 4288.94 | 0.22 | 0.823 | |
| condition: TVRR:lat[S.right]:sag[S.posterior]:OS_z | 0.00 | 0.21 | 4288.93 | 0.02 | 0.986 | |

Table S14: Summary of Linear Mixed Model for the P600 amplitude predicted by condition (TVRR vs. TVDO), laterality, sagittality, and IAF

| | Estimate | Std. Error | df | t value | Pr(>\|t\|) | |
|---|---|---|---|---|---|---|
| (Intercept) | -0.05 | 0.34 | 51.80 | -0.15 | 0.884 | |
| prestim | 0.27 | 0.02 | 4338.01 | 15.73 | <0.001 | *** |
| condition: TVRR | 0.10 | 0.40 | 67.83 | 0.24 | 0.812 | |
| lat: [S.medial] | -0.15 | 0.14 | 4289.94 | -1.05 | 0.294 | |
| lat: [S.right] | -0.26 | 0.14 | 4290.93 | -1.84 | 0.066 | . |
| sag: [S.posterior] | -0.71 | 0.10 | 4303.66 | -7.05 | <0.001 | *** |
| IAF_z | 0.16 | 0.30 | 28.75 | 0.55 | 0.586 | |
| prestim:conditionTVRR | -0.01 | 0.02 | 4338.00 | -0.42 | 0.673 | |
| condition: TVRR:lat[S.medial] | 0.19 | 0.20 | 4289.60 | 0.93 | 0.352 | |
| condition: TVRR:lat[S.right] | -0.08 | 0.20 | 4290.50 | -0.38 | 0.707 | |
| condition: TVRR:sag[S.posterior] | 0.19 | 0.14 | 4298.97 | 1.33 | 0.182 | |
| lat[S.medial]:sag[S.posterior] | -0.19 | 0.14 | 4289.24 | -1.38 | 0.169 | |
| lat[S.right]:sag[S.posterior] | 0.07 | 0.14 | 4289.25 | 0.51 | 0.609 | |
| condition: TVRR:IAF_z | -0.06 | 0.32 | 30.28 | -0.19 | 0.852 | |
| lat[S.medial]:IAF_z | 0.14 | 0.14 | 4289.24 | 0.98 | 0.329 | |
| lat[S.right]:IAF_z | -0.13 | 0.14 | 4289.30 | -0.88 | 0.378 | |
| sag[S.posterior]:IAF_z | 0.09 | 0.10 | 4289.69 | 0.90 | 0.368 | |
| condition: TVRR:lat[S.medial]:sag[S.posterior] | 0.07 | 0.20 | 4289.29 | 0.33 | 0.743 | |
| condition: TVRR:lat[S.right]:sag[S.posterior] | 0.08 | 0.20 | 4289.29 | 0.38 | 0.702 | |
| condition: TVRR:lat[S.medial]:IAF_z | 0.05 | 0.21 | 4289.24 | 0.25 | 0.804 | |
| condition: TVRR:lat[S.right]:IAF_z | 0.12 | 0.21 | 4289.28 | 0.56 | 0.575 | |
| condition: TVRR:sag[S.posterior]:IAF_z | -0.02 | 0.15 | 4289.46 | -0.12 | 0.908 | |
| lat[S.medial]:sag[S.posterior]:IAF_z | -0.06 | 0.14 | 4289.27 | -0.44 | 0.663 | |
| lat[S.right]:sag[S.posterior]:IAF_z | -0.01 | 0.14 | 4289.24 | -0.07 | 0.944 | |
| condition: TVRR:lat[S.medial]:sag[S.posterior]:IAF_z | 0.01 | 0.21 | 4289.26 | 0.06 | 0.953 | |
| condition: TVRR:lat[S.right]:sag[S.posterior]:IAF_z | 0.06 | 0.21 | 4289.28 | 0.27 | 0.785 | |

Table S15: Summary of Linear Mixed Model for the P600 amplitude predicted by condition (TVRR vs. TVDO), laterality, sagittality, and Flanker

| | Estimate | Std. Error | df | t value | Pr(>\|t\|) | |
|---|---|---|---|---|---|---|
| (Intercept) | -0.05 | 0.33 | 52.93 | -0.15 | 0.884 | |
| prestim | 0.27 | 0.02 | 4332.98 | 15.78 | <0.001 | *** |
| condition: TVRR | 0.08 | 0.40 | 68.42 | 0.21 | 0.834 | |
| lat: [S.medial] | -0.14 | 0.14 | 4290.34 | -1.01 | 0.311 | |
| lat: [S.right] | -0.25 | 0.14 | 4291.28 | -1.76 | 0.079 | . |
| sag: [S.posterior] | -0.69 | 0.10 | 4303.95 | -6.89 | <0.001 | *** |
| flanker_z | -0.37 | 0.30 | 27.27 | -1.22 | 0.232 | |
| prestim:conditionTVRR | -0.01 | 0.02 | 4339.53 | -0.44 | 0.663 | |
| condition: TVRR:lat[S.medial] | 0.19 | 0.20 | 4290.02 | 0.94 | 0.35 | |
| condition: TVRR:lat[S.right] | -0.07 | 0.20 | 4290.89 | -0.34 | 0.737 | |
| condition: TVRR:sag[S.posterior] | 0.17 | 0.14 | 4299.33 | 1.22 | 0.223 | |
| lat[S.medial]:sag[S.posterior] | -0.20 | 0.14 | 4289.71 | -1.42 | 0.156 | |
| lat[S.right]:sag[S.posterior] | 0.07 | 0.14 | 4289.70 | 0.47 | 0.641 | |
| conditionTVRR:flanker_z | -0.11 | 0.33 | 28.55 | -0.35 | 0.731 | |
| lat[S.medial]:flanker_z | -0.15 | 0.15 | 4289.91 | -0.95 | 0.341 | |
| lat[S.right]:flanker_z | 0.37 | 0.15 | 4289.72 | 2.40 | 0.016 | * |
| sag[S.posterior]:flanker_z | 0.11 | 0.11 | 4289.92 | 1.00 | 0.319 | |
| condition: TVRR:lat[S.medial]:sag[S.posterior] | 0.07 | 0.20 | 4289.75 | 0.34 | 0.737 | |
| condition: TVRR:lat[S.right]:sag[S.posterior] | 0.09 | 0.20 | 4289.75 | 0.43 | 0.667 | |
| conditionTVRR:lat[S.medial]:flanker_z | -0.20 | 0.22 | 4289.79 | -0.94 | 0.349 | |
| conditionTVRR:lat[S.right]:flanker_z | 0.04 | 0.22 | 4289.85 | 0.20 | 0.842 | |
| conditionTVRR:sag[S.posterior]:flanker_z | -0.24 | 0.15 | 4290.02 | -1.54 | 0.123 | |
| lat[S.medial]:sag[S.posterior]:flanker_z | 0.04 | 0.15 | 4289.90 | 0.25 | 0.805 | |
| lat[S.right]:sag[S.posterior]:flanker_z | -0.07 | 0.15 | 4289.70 | -0.45 | 0.652 | |
| conditionTVRR:lat[S.medial]:sag[S.posterior]:flanker_z | 0.01 | 0.22 | 4289.81 | 0.06 | 0.953 | |
| conditionTVRR:lat[S.right]:sag[S.posterior]:flanker_z | -0.00 | 0.22 | 4289.69 | -0.01 | 0.996 | |

Table S16: Summary of Linear Mixed Model for the P600 amplitude predicted by condition (TVRR vs. TVDO), laterality, sagittality, and Stroop

| | Estimate | Std. Error | df | t value | Pr(>\|t\|) | |
|---|---|---|---|---|---|---|
| (Intercept) | -0.03 | 0.34 | 51.83 | -0.10 | 0.922 | |
| prestim | 0.27 | 0.02 | 4339.76 | 15.64 | <0.001 | *** |
| condition: TVRR | 0.08 | 0.40 | 68.14 | 0.19 | 0.85 | |
| lat: [S.medial] | -0.13 | 0.14 | 4290.11 | -0.95 | 0.345 | |
| lat: [S.right] | -0.29 | 0.14 | 4291.20 | -2.05 | 0.041 | * |
| sag: [S.posterior] | -0.71 | 0.10 | 4303.67 | -7.10 | <0.001 | *** |
| stroop_z | 0.07 | 0.29 | 28.74 | 0.24 | 0.81 | |
| prestim:conditionTVRR | -0.01 | 0.02 | 4354.89 | -0.33 | 0.739 | |
| condition: TVRR:lat[S.medial] | 0.18 | 0.20 | 4289.75 | 0.93 | 0.354 | |
| condition: TVRR:lat[S.right] | -0.04 | 0.20 | 4290.73 | -0.22 | 0.826 | |
| condition: TVRR:sag[S.posterior] | 0.20 | 0.14 | 4299.11 | 1.41 | 0.16 | |
| lat[S.medial]:sag[S.posterior] | -0.19 | 0.14 | 4289.37 | -1.37 | 0.172 | |
| lat[S.right]:sag[S.posterior] | 0.06 | 0.14 | 4289.37 | 0.46 | 0.646 | |
| condition: TVRR:stroop_z | 0.16 | 0.31 | 29.92 | 0.50 | 0.62 | |
| lat[S.medial]:stroop_z | 0.03 | 0.15 | 4289.42 | 0.22 | 0.826 | |
| lat[S.right]:stroop_z | 0.13 | 0.15 | 4290.19 | 0.91 | 0.362 | |
| sag[S.posterior]:stroop_z | 0.13 | 0.10 | 4289.73 | 1.28 | 0.201 | |
| condition: TVRR:lat[S.medial]:sag[S.posterior] | 0.06 | 0.20 | 4289.41 | 0.29 | 0.772 | |
| condition: TVRR:lat[S.right]:sag[S.posterior] | 0.09 | 0.20 | 4289.42 | 0.45 | 0.653 | |
| condition: TVRR:lat[S.medial]:stroop_z | 0.17 | 0.21 | 4289.49 | 0.85 | 0.397 | |
| condition: TVRR:lat[S.right]:stroop_z | -0.22 | 0.21 | 4289.90 | -1.05 | 0.293 | |
| condition: TVRR:sag[S.posterior]:stroop_z | -0.08 | 0.15 | 4289.65 | -0.58 | 0.562 | |
| lat[S.medial]:sag[S.posterior]:stroop_z | -0.11 | 0.15 | 4289.37 | -0.76 | 0.446 | |
| lat[S.right]:sag[S.posterior]:stroop_z | 0.08 | 0.15 | 4289.36 | 0.51 | 0.607 | |
| condition: TVRR:lat[S.medial]:sag[S.posterior]:stroop_z | 0.10 | 0.21 | 4289.37 | 0.49 | 0.623 | |
| condition: TVRR:lat[S.right]:sag[S.posterior]:stroop_z | -0.06 | 0.21 | 4289.36 | -0.31 | 0.756 | |

Table S17: Summary of Linear Mixed Model for the P600 amplitude predicted by condition (IVWR vs. TVRR), laterality, and sagittality

| | Estimate | Std. Error | df | t value | Pr(>\|t\|) | |
|---|---|---|---|---|---|---|
| (Intercept) | 0.25 | 0.39 | 53.99 | 0.64 | 0.523 | |
| prestim | 0.27 | 0.02 | 4409.01 | 15.40 | <0.001 | *** |
| condition: IVWR | 0.80 | 0.48 | 68.47 | 1.68 | 0.098 | . |
| lat: [S.medial] | -0.16 | 0.14 | 4292.14 | -1.17 | 0.241 | |
| lat: [S.right] | -0.49 | 0.14 | 4292.93 | -3.58 | <0.001 | *** |
| sag: [S.posterior] | -1.03 | 0.10 | 4295.91 | -10.66 | <0.001 | *** |
| prestim:conditionIVWR | 0.03 | 0.02 | 4431.26 | 1.27 | 0.204 | |
| condition: IVWR:lat[S.medial] | 0.40 | 0.19 | 4292.78 | 2.10 | 0.036 | * |
| condition: IVWR:lat[S.right] | -0.02 | 0.19 | 4292.57 | -0.13 | 0.899 | |
| condition: IVWR:sag[S.posterior] | 0.45 | 0.14 | 4296.97 | 3.36 | 0.001 | *** |
| lat[S.medial]:sag[S.posterior] | -0.09 | 0.14 | 4291.96 | -0.65 | 0.516 | |
| lat[S.right]:sag[S.posterior] | 0.03 | 0.14 | 4292.07 | 0.19 | 0.852 | |
| condition: IVWR:lat[S.medial]:sag[S.posterior] | -0.05 | 0.19 | 4292.00 | -0.26 | 0.796 | |
| condition: IVWR:lat[S.right]:sag[S.posterior] | 0.04 | 0.19 | 4292.04 | 0.19 | 0.846 | |

Table S18: Summary of Linear Mixed Model for the P600 amplitude predicted by condition (IVWR vs. TVRR), laterality, sagittality, and age

| | Estimate | Std. Error | df | t value | Pr(>\|t\|) | |
|---|---|---|---|---|---|---|
| (Intercept) | 0.33 | 0.38 | 56.00 | 0.88 | 0.382 | |
| prestim | 0.27 | 0.02 | 4383.02 | 15.38 | <0.001 | *** |
| condition: IVWR | 0.66 | 0.45 | 75.71 | 1.47 | 0.147 | |
| lat: [S.medial] | -0.14 | 0.14 | 4290.70 | -1.03 | 0.303 | |
| lat: [S.right] | -0.45 | 0.14 | 4291.35 | -3.26 | 0.001 | ** |
| sag: [S.posterior] | -1.03 | 0.10 | 4294.36 | -10.46 | <0.001 | *** |
| age_z | 0.55 | 0.33 | 25.85 | 1.68 | 0.106 | |
| prestim:conditionIVWR | 0.03 | 0.02 | 4408.80 | 1.35 | 0.176 | |
| condition: IVWR:lat[S.medial] | 0.34 | 0.19 | 4291.29 | 1.78 | 0.075 | . |
| condition: IVWR:lat[S.right] | -0.03 | 0.19 | 4291.06 | -0.16 | 0.871 | |
| condition: IVWR:sag[S.posterior] | 0.42 | 0.14 | 4295.36 | 3.08 | 0.002 | ** |
| lat[S.medial]:sag[S.posterior] | -0.09 | 0.14 | 4290.49 | -0.65 | 0.514 | |
| lat[S.right]:sag[S.posterior] | 0.03 | 0.14 | 4290.60 | 0.19 | 0.848 | |
| condition: IVWR:age_z | -0.95 | 0.35 | 26.07 | -2.74 | 0.011 | * |
| lat[S.medial]:age_z | 0.09 | 0.15 | 4290.49 | 0.62 | 0.533 | |
| lat[S.right]:age_z | 0.20 | 0.15 | 4290.57 | 1.36 | 0.175 | |
| sag[S.posterior]:age_z | 0.01 | 0.10 | 4290.55 | 0.13 | 0.9 | |
| condition: IVWR:lat[S.medial]:sag[S.posterior] | -0.03 | 0.19 | 4290.51 | -0.15 | 0.877 | |
| condition: IVWR:lat[S.right]:sag[S.posterior] | 0.03 | 0.19 | 4290.57 | 0.15 | 0.88 | |
| condition: IVWR:lat[S.medial]:age_z | -0.31 | 0.21 | 4290.48 | -1.51 | 0.131 | |
| condition: IVWR:lat[S.right]:age_z | -0.03 | 0.21 | 4290.52 | -0.16 | 0.873 | |
| condition: IVWR:sag[S.posterior]:age_z | -0.18 | 0.15 | 4290.65 | -1.25 | 0.212 | |
| lat[S.medial]:sag[S.posterior]:age_z | -0.01 | 0.15 | 4290.48 | -0.07 | 0.941 | |
| lat[S.right]:sag[S.posterior]:age_z | 0.01 | 0.15 | 4290.46 | 0.04 | 0.97 | |
| condition: IVWR:lat[S.medial]:sag[S.posterior]:age_z | 0.11 | 0.21 | 4290.47 | 0.53 | 0.598 | |
| condition: IVWR:lat[S.right]:sag[S.posterior]:age_z | -0.04 | 0.21 | 4290.46 | -0.21 | 0.835 | |

Table S19: Summary of Linear Mixed Model for the P600 amplitude predicted by condition (IVWR vs. TVRR), laterality, sagittality, and PTA

| | Estimate | Std. Error | df | t value | Pr(>\|t\|) | |
|---|---|---|---|---|---|---|
| (Intercept) | 0.25 | 0.35 | 63.67 | 0.71 | 0.483 | |
| prestim | 0.27 | 0.02 | 4392.96 | 15.53 | <0.001 | *** |
| condition: IVWR | 0.79 | 0.46 | 72.26 | 1.71 | 0.092 | . |
| lat: [S.medial] | -0.16 | 0.14 | 4292.09 | -1.16 | 0.248 | |
| lat: [S.right] | -0.49 | 0.14 | 4292.92 | -3.59 | <0.001 | *** |
| sag: [S.posterior] | -1.03 | 0.10 | 4296.07 | -10.71 | <0.001 | *** |
| PTA_z | 0.71 | 0.28 | 27.48 | 2.57 | 0.016 | * |
| att_resid_z | 0.55 | 0.25 | 29.22 | 2.20 | 0.036 | * |
| prestim:conditionIVWR | 0.03 | 0.02 | 4421.81 | 1.36 | 0.175 | |
| condition: IVWR:lat[S.medial] | 0.40 | 0.19 | 4292.73 | 2.09 | 0.037 | * |
| condition: IVWR:lat[S.right] | -0.03 | 0.19 | 4292.53 | -0.14 | 0.888 | |
| condition: IVWR:sag[S.posterior] | 0.45 | 0.14 | 4296.96 | 3.31 | 0.001 | *** |
| lat[S.medial]:sag[S.posterior] | -0.09 | 0.14 | 4291.90 | -0.64 | 0.519 | |
| lat[S.right]:sag[S.posterior] | 0.02 | 0.14 | 4292.01 | 0.18 | 0.856 | |
| condition: IVWR:PTA_z | -0.60 | 0.35 | 27.54 | -1.74 | 0.093 | . |
| lat[S.medial]:PTA_z | 0.17 | 0.13 | 4291.92 | 1.32 | 0.188 | |
| lat[S.right]:PTA_z | 0.03 | 0.13 | 4291.94 | 0.25 | 0.801 | |
| sag[S.posterior]:PTA_z | -0.16 | 0.09 | 4292.38 | -1.70 | 0.09 | . |
| condition: IVWR:lat[S.medial]:sag[S.posterior] | -0.05 | 0.19 | 4291.93 | -0.25 | 0.804 | |
| condition: IVWR:lat[S.right]:sag[S.posterior] | 0.04 | 0.19 | 4291.98 | 0.19 | 0.846 | |
| condition: IVWR:lat[S.medial]:PTA_z | -0.24 | 0.18 | 4291.91 | -1.28 | 0.202 | |
| condition: IVWR:lat[S.right]:PTA_z | -0.10 | 0.18 | 4291.91 | -0.54 | 0.592 | |
| condition: IVWR:sag[S.posterior]:PTA_z | -0.23 | 0.13 | 4292.39 | -1.75 | 0.081 | . |
| lat[S.medial]:sag[S.posterior]:PTA_z | 0.05 | 0.13 | 4291.88 | 0.35 | 0.724 | |
| lat[S.right]:sag[S.posterior]:PTA_z | -0.03 | 0.13 | 4291.88 | -0.26 | 0.794 | |
| condition: IVWR:lat[S.medial]:sag[S.posterior]:PTA_z | 0.03 | 0.18 | 4291.89 | 0.15 | 0.882 | |
| condition: IVWR:lat[S.right]:sag[S.posterior]:PTA_z | 0.02 | 0.18 | 4291.88 | 0.10 | 0.92 | |

Table S20: Summary of Linear Mixed Model for the P600 amplitude predicted by condition (IVWR vs. TVRR), laterality, sagittality, and RS

| | Estimate | Std. Error | df | t value | Pr(>\|t\|) | |
|---|---|---|---|---|---|---|
| (Intercept) | 0.24 | 0.38 | 56.35 | 0.63 | 0.533 | |
| prestim | 0.27 | 0.02 | 4397.30 | 15.48 | <0.001 | *** |
| condition: IVWR | 0.81 | 0.46 | 72.59 | 1.76 | 0.083 | . |
| lat: [S.medial] | -0.16 | 0.14 | 4292.21 | -1.18 | 0.236 | |
| lat: [S.right] | -0.49 | 0.14 | 4293.03 | -3.59 | <0.001 | *** |
| sag: [S.posterior] | -1.03 | 0.10 | 4296.09 | -10.68 | <0.001 | *** |
| RS_z | -0.45 | 0.31 | 26.28 | -1.47 | 0.154 | |
| prestim:conditionIVWR | 0.03 | 0.02 | 4422.53 | 1.22 | 0.223 | |
| condition: IVWR:lat[S.medial] | 0.41 | 0.19 | 4292.84 | 2.14 | 0.032 | * |
| condition: IVWR:lat[S.right] | -0.03 | 0.19 | 4292.65 | -0.14 | 0.891 | |
| condition: IVWR:sag[S.posterior] | 0.46 | 0.14 | 4297.15 | 3.43 | 0.001 | *** |
| lat[S.medial]:sag[S.posterior] | -0.09 | 0.14 | 4292.02 | -0.65 | 0.514 | |
| lat[S.right]:sag[S.posterior] | 0.03 | 0.14 | 4292.13 | 0.18 | 0.854 | |
| condition: IVWR:RS_z | 0.60 | 0.34 | 26.61 | 1.73 | 0.096 | . |
| lat[S.medial]:RS_z | -0.11 | 0.13 | 4292.01 | -0.83 | 0.405 | |
| lat[S.right]:RS_z | 0.10 | 0.13 | 4292.25 | 0.78 | 0.436 | |
| sag[S.posterior]:RS_z | 0.24 | 0.09 | 4292.01 | 2.57 | 0.01 | * |
| condition: IVWR:lat[S.medial]:sag[S.posterior] | -0.05 | 0.19 | 4292.06 | -0.26 | 0.796 | |
| condition: IVWR:lat[S.right]:sag[S.posterior] | 0.04 | 0.19 | 4292.11 | 0.19 | 0.846 | |
| condition: IVWR:lat[S.medial]:RS_z | 0.37 | 0.19 | 4292.05 | 1.96 | 0.05 | . |
| condition: IVWR:lat[S.right]:RS_z | -0.12 | 0.19 | 4292.15 | -0.63 | 0.529 | |
| condition: IVWR:sag[S.posterior]:RS_z | 0.18 | 0.13 | 4292.00 | 1.36 | 0.175 | |
| lat[S.medial]:sag[S.posterior]:RS_z | -0.01 | 0.13 | 4292.04 | -0.05 | 0.958 | |
| lat[S.right]:sag[S.posterior]:RS_z | -0.03 | 0.13 | 4292.00 | -0.24 | 0.807 | |
| condition: IVWR:lat[S.medial]:sag[S.posterior]:RS_z | 0.01 | 0.19 | 4292.02 | 0.07 | 0.942 | |
| condition: IVWR:lat[S.right]:sag[S.posterior]:RS_z | 0.02 | 0.19 | 4292.01 | 0.09 | 0.931 | |

Table S21: Summary of Linear Mixed Model for the P600 amplitude predicted by condition (IVWR vs. TVRR), laterality, sagittality, and OS

| | Estimate | Std. Error | df | t value | Pr(>\|t\|) | |
|---|---|---|---|---|---|---|
| (Intercept) | 0.28 | 0.38 | 55.84 | 0.75 | 0.459 | |
| prestim | 0.27 | 0.02 | 4404.09 | 15.41 | <0.001 | *** |
| condition: IVWR | 0.77 | 0.47 | 69.41 | 1.63 | 0.107 | |
| lat: [S.medial] | -0.15 | 0.14 | 4292.24 | -1.07 | 0.283 | |
| lat: [S.right] | -0.50 | 0.14 | 4292.96 | -3.62 | <0.001 | *** |
| sag: [S.posterior] | -1.03 | 0.10 | 4295.94 | -10.61 | <0.001 | *** |
| OS_z | -0.44 | 0.33 | 27.11 | -1.35 | 0.189 | |
| prestim:conditionIVWR | 0.03 | 0.02 | 4429.99 | 1.27 | 0.206 | |
| condition: IVWR:lat[S.medial] | 0.38 | 0.19 | 4292.87 | 1.99 | 0.047 | * |
| condition: IVWR:lat[S.right] | -0.02 | 0.19 | 4292.62 | -0.09 | 0.929 | |
| condition: IVWR:sag[S.posterior] | 0.45 | 0.14 | 4296.96 | 3.27 | 0.001 | ** |
| lat[S.medial]:sag[S.posterior] | -0.09 | 0.14 | 4292.06 | -0.63 | 0.529 | |
| lat[S.right]:sag[S.posterior] | 0.03 | 0.14 | 4292.17 | 0.20 | 0.845 | |
| condition: IVWR:OS_z | 0.30 | 0.38 | 27.48 | 0.79 | 0.438 | |
| lat[S.medial]:OS_z | -0.15 | 0.14 | 4292.04 | -1.03 | 0.302 | |
| lat[S.right]:OS_z | 0.08 | 0.14 | 4292.20 | 0.59 | 0.557 | |
| sag[S.posterior]:OS_z | -0.02 | 0.10 | 4292.11 | -0.16 | 0.873 | |
| condition: IVWR:lat[S.medial]:sag[S.posterior] | -0.05 | 0.19 | 4292.09 | -0.29 | 0.774 | |
| condition: IVWR:lat[S.right]:sag[S.posterior] | 0.04 | 0.19 | 4292.15 | 0.21 | 0.836 | |
| condition: IVWR:lat[S.medial]:OS_z | 0.21 | 0.20 | 4292.04 | 1.03 | 0.304 | |
| condition: IVWR:lat[S.right]:OS_z | -0.08 | 0.20 | 4292.13 | -0.42 | 0.676 | |
| condition: IVWR:sag[S.posterior]:OS_z | 0.10 | 0.14 | 4292.09 | 0.69 | 0.489 | |
| lat[S.medial]:sag[S.posterior]:OS_z | -0.03 | 0.14 | 4292.04 | -0.19 | 0.848 | |
| lat[S.right]:sag[S.posterior]:OS_z | -0.02 | 0.14 | 4292.04 | -0.11 | 0.912 | |
| condition: IVWR:lat[S.medial]:sag[S.posterior]:OS_z | 0.06 | 0.20 | 4292.04 | 0.31 | 0.76 | |
| condition: IVWR:lat[S.right]:sag[S.posterior]:OS_z | -0.03 | 0.20 | 4292.06 | -0.13 | 0.9 | |

Table S22: Summary of Linear Mixed Model for the P600 amplitude predicted by condition (IVWR vs. TVRR), laterality, sagittality, and IAF

| | Estimate | Std. Error | df | t value | Pr(>\|t\|) | |
|---|---|---|---|---|---|---|
| (Intercept) | 0.26 | 0.39 | 53.68 | 0.66 | 0.51 | |
| prestim | 0.27 | 0.02 | 4404.06 | 15.38 | <0.001 | *** |
| condition: IVWR | 0.70 | 0.45 | 73.88 | 1.56 | 0.123 | |
| lat: [S.medial] | -0.17 | 0.14 | 4291.00 | -1.23 | 0.218 | |
| lat: [S.right] | -0.48 | 0.14 | 4291.82 | -3.55 | <0.001 | *** |
| sag: [S.posterior] | -1.04 | 0.10 | 4294.90 | -10.78 | <0.001 | *** |
| IAF_z | -0.10 | 0.34 | 27.82 | -0.31 | 0.758 | |
| prestim:conditionIVWR | 0.03 | 0.02 | 4420.50 | 1.37 | 0.17 | |
| condition: IVWR:lat[S.medial] | 0.37 | 0.19 | 4291.64 | 1.96 | 0.051 | . |
| condition: IVWR:lat[S.right] | -0.01 | 0.19 | 4291.50 | -0.07 | 0.948 | |
| condition: IVWR:sag[S.posterior] | 0.46 | 0.14 | 4296.16 | 3.36 | 0.001 | *** |
| lat[S.medial]:sag[S.posterior] | -0.09 | 0.14 | 4290.83 | -0.68 | 0.499 | |
| lat[S.right]:sag[S.posterior] | 0.02 | 0.14 | 4290.94 | 0.17 | 0.864 | |
| condition: IVWR:IAF_z | 0.85 | 0.35 | 27.20 | 2.46 | 0.021 | * |
| lat[S.medial]:IAF_z | 0.12 | 0.14 | 4290.81 | 0.86 | 0.388 | |
| lat[S.right]:IAF_z | -0.06 | 0.14 | 4290.81 | -0.42 | 0.678 | |
| sag[S.posterior]:IAF_z | 0.17 | 0.10 | 4290.84 | 1.72 | 0.085 | . |
| condition: IVWR:lat[S.medial]:sag[S.posterior] | -0.03 | 0.19 | 4290.86 | -0.16 | 0.876 | |
| condition: IVWR:lat[S.right]:sag[S.posterior] | 0.02 | 0.19 | 4290.92 | 0.12 | 0.901 | |
| condition: IVWR:lat[S.medial]:IAF_z | 0.17 | 0.20 | 4290.86 | 0.87 | 0.383 | |
| condition: IVWR:lat[S.right]:IAF_z | -0.07 | 0.20 | 4290.98 | -0.35 | 0.726 | |
| condition: IVWR:sag[S.posterior]:IAF_z | -0.07 | 0.14 | 4290.82 | -0.52 | 0.605 | |
| lat[S.medial]:sag[S.posterior]:IAF_z | 0.05 | 0.14 | 4290.85 | 0.36 | 0.717 | |
| lat[S.right]:sag[S.posterior]:IAF_z | 0.03 | 0.14 | 4290.83 | 0.21 | 0.831 | |
| condition: IVWR:lat[S.medial]:sag[S.posterior]:IAF_z | -0.18 | 0.20 | 4290.83 | -0.92 | 0.357 | |
| condition: IVWR:lat[S.right]:sag[S.posterior]:IAF_z | 0.10 | 0.20 | 4290.82 | 0.49 | 0.627 | |

Table S23: Summary of Linear Mixed Model for the P600 amplitude predicted by condition (IVWR vs. TVRR), laterality, sagittality, and Flanker

| | Estimate | Std. Error | df | t value | Pr(>\|t\|) | |
|---|---|---|---|---|---|---|
| (Intercept) | 0.24 | 0.39 | 54.04 | 0.63 | 0.532 | |
| prestim | 0.27 | 0.02 | 4410.48 | 15.42 | <0.001 | *** |
| condition: IVWR | 0.80 | 0.48 | 68.44 | 1.68 | 0.098 | . |
| lat: [S.medial] | -0.17 | 0.14 | 4292.19 | -1.21 | 0.226 | |
| lat: [S.right] | -0.49 | 0.14 | 4292.94 | -3.60 | <0.001 | *** |
| sag: [S.posterior] | -1.02 | 0.10 | 4295.99 | -10.51 | <0.001 | *** |
| flanker_z | -0.11 | 0.34 | 26.04 | -0.31 | 0.756 | |
| prestim:conditionIVWR | 0.03 | 0.02 | 4432.08 | 1.26 | 0.209 | |
| condition: IVWR:lat[S.medial] | 0.39 | 0.19 | 4292.88 | 2.02 | 0.043 | * |
| condition: IVWR:lat[S.right] | -0.02 | 0.19 | 4292.58 | -0.10 | 0.92 | |
| condition: IVWR:sag[S.posterior] | 0.44 | 0.14 | 4297.14 | 3.27 | 0.001 | ** |
| lat[S.medial]:sag[S.posterior] | -0.09 | 0.14 | 4292.03 | -0.67 | 0.501 | |
| lat[S.right]:sag[S.posterior] | 0.03 | 0.14 | 4292.13 | 0.20 | 0.842 | |
| conditionIVWR:flanker_z | 0.02 | 0.39 | 26.57 | 0.06 | 0.95 | |
| lat[S.medial]:flanker_z | -0.09 | 0.15 | 4292.16 | -0.57 | 0.567 | |
| lat[S.right]:flanker_z | -0.05 | 0.15 | 4292.14 | -0.32 | 0.749 | |
| sag[S.posterior]:flanker_z | 0.20 | 0.11 | 4292.04 | 1.87 | 0.062 | . |
| condition: IVWR:lat[S.medial]:sag[S.posterior] | -0.05 | 0.19 | 4292.07 | -0.24 | 0.809 | |
| condition: IVWR:lat[S.right]:sag[S.posterior] | 0.03 | 0.19 | 4292.11 | 0.16 | 0.875 | |
| conditionIVWR:lat[S.medial]:flanker_z | -0.10 | 0.21 | 4292.14 | -0.49 | 0.626 | |
| conditionIVWR:lat[S.right]:flanker_z | 0.07 | 0.21 | 4292.20 | 0.33 | 0.745 | |
| conditionIVWR:sag[S.posterior]:flanker_z | -0.17 | 0.15 | 4292.14 | -1.12 | 0.261 | |
| lat[S.medial]:sag[S.posterior]:flanker_z | -0.05 | 0.15 | 4292.01 | -0.34 | 0.732 | |
| lat[S.right]:sag[S.posterior]:flanker_z | 0.03 | 0.15 | 4292.05 | 0.18 | 0.855 | |
| conditionIVWR:lat[S.medial]:sag[S.posterior]:flanker_z | 0.05 | 0.21 | 4292.01 | 0.22 | 0.825 | |
| conditionIVWR:lat[S.right]:sag[S.posterior]:flanker_z | -0.08 | 0.21 | 4292.03 | -0.38 | 0.704 | |

Table S24: Summary of Linear Mixed Model for the P600 amplitude predicted by condition (IVWR vs. TVRR), laterality, sagittality, and Stroop

| | Estimate | Std. Error | df | t value | Pr(>\|t\|) | |
|---|---|---|---|---|---|---|
| (Intercept) | 0.23 | 0.38 | 54.42 | 0.60 | 0.55 | |
| prestim | 0.27 | 0.02 | 4409.09 | 15.41 | <0.001 | *** |
| condition: IVWR | 0.80 | 0.48 | 68.07 | 1.68 | 0.098 | . |
| lat: [S.medial] | -0.17 | 0.14 | 4291.97 | -1.23 | 0.219 | |
| lat: [S.right] | -0.49 | 0.14 | 4292.80 | -3.58 | <0.001 | *** |
| sag: [S.posterior] | -1.03 | 0.10 | 4295.71 | -10.67 | <0.001 | *** |
| stroop_z | 0.24 | 0.32 | 27.03 | 0.73 | 0.469 | |
| prestim:conditionIVWR | 0.03 | 0.02 | 4432.29 | 1.23 | 0.221 | |
| condition: IVWR:lat[S.medial] | 0.39 | 0.19 | 4292.61 | 2.06 | 0.04 | * |
| condition: IVWR:lat[S.right] | -0.02 | 0.19 | 4292.43 | -0.12 | 0.903 | |
| condition: IVWR:sag[S.posterior] | 0.45 | 0.14 | 4296.83 | 3.29 | 0.001 | *** |
| lat[S.medial]:sag[S.posterior] | -0.09 | 0.14 | 4291.77 | -0.68 | 0.494 | |
| lat[S.right]:sag[S.posterior] | 0.02 | 0.14 | 4291.87 | 0.17 | 0.861 | |
| condition: IVWR:stroop_z | -0.02 | 0.37 | 27.28 | -0.07 | 0.948 | |
| lat[S.medial]:stroop_z | 0.11 | 0.14 | 4291.86 | 0.82 | 0.411 | |
| lat[S.right]:stroop_z | 0.01 | 0.14 | 4291.97 | 0.06 | 0.951 | |
| sag[S.posterior]:stroop_z | 0.03 | 0.10 | 4291.75 | 0.32 | 0.747 | |
| condition: IVWR:lat[S.medial]:sag[S.posterior] | -0.04 | 0.19 | 4291.81 | -0.19 | 0.846 | |
| condition: IVWR:lat[S.right]:sag[S.posterior] | 0.03 | 0.19 | 4291.85 | 0.18 | 0.854 | |
| condition: IVWR:lat[S.medial]:stroop_z | 0.10 | 0.20 | 4291.81 | 0.51 | 0.608 | |
| condition: IVWR:lat[S.right]:stroop_z | -0.02 | 0.20 | 4291.99 | -0.09 | 0.928 | |
| condition: IVWR:sag[S.posterior]:stroop_z | 0.11 | 0.14 | 4291.82 | 0.77 | 0.443 | |
| lat[S.medial]:sag[S.posterior]:stroop_z | 0.07 | 0.14 | 4291.75 | 0.52 | 0.605 | |
| lat[S.right]:sag[S.posterior]:stroop_z | 0.02 | 0.14 | 4291.76 | 0.16 | 0.87 | |
| condition: IVWR:lat[S.medial]:sag[S.posterior]:stroop_z | -0.18 | 0.20 | 4291.75 | -0.94 | 0.349 | |
| condition: IVWR:lat[S.right]:sag[S.posterior]:stroop_z | 0.03 | 0.20 | 4291.77 | 0.17 | 0.864 | |

Table S25: Summary of Linear Mixed Model for the N400 amplitude predicted by condition (ORAI vs. ORIA), laterality, and sagittality

| | Estimate | Std. Error | df | t value | Pr(>\|t\|) | |
|---|---|---|---|---|---|---|
| (Intercept) | 0.74 | 0.34 | 52.71 | 2.16 | 0.035 | * |
| prestim | 0.28 | 0.02 | 4122.70 | 15.22 | <0.001 | *** |
| condition: ORAI | -0.20 | 0.39 | 57.13 | -0.51 | 0.615 | |
| lat: [S.medial] | 0.41 | 0.15 | 4367.24 | 2.70 | 0.007 | ** |
| lat: [S.right] | -0.43 | 0.15 | 4367.82 | -2.86 | 0.004 | ** |
| sag: [S.posterior] | -0.02 | 0.11 | 4383.99 | -0.21 | 0.835 | |
| prestim:conditionORAI | 0.06 | 0.03 | 3975.78 | 2.17 | 0.03 | * |
| condition: ORAI:lat[S.medial] | -0.14 | 0.21 | 4367.24 | -0.64 | 0.523 | |
| condition: ORAI:lat[S.right] | 0.03 | 0.21 | 4368.15 | 0.16 | 0.874 | |
| condition: ORAI:sag[S.posterior] | -0.22 | 0.15 | 4383.78 | -1.44 | 0.151 | |
| lat[S.medial]:sag[S.posterior] | -0.02 | 0.15 | 4367.56 | -0.14 | 0.887 | |
| lat[S.right]:sag[S.posterior] | 0.00 | 0.15 | 4367.24 | 0.00 | 1 | |
| condition: ORAI:lat[S.medial]:sag[S.posterior] | -0.01 | 0.21 | 4367.59 | -0.04 | 0.964 | |
| condition: ORAI:lat[S.right]:sag[S.posterior] | 0.05 | 0.21 | 4367.27 | 0.22 | 0.823 | |

Table S26: Summary of Linear Mixed Model for the N400 amplitude predicted by condition (ORAI vs. ORIA), laterality, sagittality, and age

| | Estimate | Std. Error | df | t value | Pr(>\|t\|) | |
|---|---|---|---|---|---|---|
| (Intercept) | 0.72 | 0.34 | 52.31 | 2.09 | 0.041 | * |
| prestim | 0.28 | 0.02 | 4126.64 | 15.21 | <0.001 | *** |
| condition: ORAI | -0.23 | 0.39 | 57.14 | -0.59 | 0.56 | |
| lat: [S.medial] | 0.39 | 0.15 | 4367.10 | 2.57 | 0.01 | * |
| lat: [S.right] | -0.43 | 0.15 | 4367.70 | -2.82 | 0.005 | ** |
| sag: [S.posterior] | -0.03 | 0.11 | 4383.88 | -0.29 | 0.772 | |
| age_z | -0.14 | 0.30 | 28.29 | -0.48 | 0.635 | |
| prestim:conditionORAI | 0.06 | 0.03 | 3976.97 | 2.21 | 0.027 | * |
| condition: ORAI:lat[S.medial] | -0.14 | 0.22 | 4367.11 | -0.63 | 0.529 | |
| condition: ORAI:lat[S.right] | 0.04 | 0.22 | 4368.03 | 0.18 | 0.859 | |
| condition: ORAI:sag[S.posterior] | -0.22 | 0.16 | 4383.44 | -1.40 | 0.162 | |
| lat[S.medial]:sag[S.posterior] | -0.02 | 0.15 | 4367.40 | -0.10 | 0.918 | |
| lat[S.right]:sag[S.posterior] | 0.00 | 0.15 | 4367.10 | 0.01 | 0.989 | |
| condition: ORAI:age_z | -0.23 | 0.29 | 26.51 | -0.79 | 0.437 | |
| lat[S.medial]:age_z | -0.13 | 0.16 | 4367.16 | -0.80 | 0.421 | |
| lat[S.right]:age_z | 0.02 | 0.16 | 4367.14 | 0.10 | 0.923 | |
| sag[S.posterior]:age_z | -0.07 | 0.11 | 4367.27 | -0.61 | 0.54 | |
| condition: ORAI:lat[S.medial]:sag[S.posterior] | -0.01 | 0.22 | 4367.44 | -0.06 | 0.952 | |
| condition: ORAI:lat[S.right]:sag[S.posterior] | 0.04 | 0.22 | 4367.13 | 0.21 | 0.837 | |
| condition: ORAI:lat[S.medial]:age_z | 0.01 | 0.23 | 4367.23 | 0.03 | 0.975 | |
| condition: ORAI:lat[S.right]:age_z | 0.03 | 0.23 | 4367.12 | 0.13 | 0.893 | |
| condition: ORAI:sag[S.posterior]:age_z | 0.02 | 0.16 | 4367.18 | 0.13 | 0.897 | |
| lat[S.medial]:sag[S.posterior]:age_z | 0.04 | 0.16 | 4367.09 | 0.27 | 0.784 | |
| lat[S.right]:sag[S.posterior]:age_z | 0.02 | 0.16 | 4367.13 | 0.09 | 0.925 | |
| condition: ORAI:lat[S.medial]:sag[S.posterior]:age_z | -0.03 | 0.23 | 4367.09 | -0.11 | 0.911 | |
| condition: ORAI:lat[S.right]:sag[S.posterior]:age_z | -0.03 | 0.23 | 4367.11 | -0.12 | 0.902 | |

Table S27: Summary of Linear Mixed Model for the N400 amplitude predicted by condition (ORAI vs. ORIA), laterality, sagittality, and PTA

| | Estimate | Std. Error | df | t value | Pr(>\|t\|) | |
|---|---|---|---|---|---|---|
| (Intercept) | 0.74 | 0.33 | 54.20 | 2.20 | 0.032 | * |
| prestim | 0.28 | 0.02 | 4115.49 | 15.16 | <0.001 | *** |
| condition: ORAI | -0.20 | 0.39 | 57.17 | -0.51 | 0.613 | |
| lat: [S.medial] | 0.41 | 0.15 | 4367.64 | 2.71 | 0.007 | ** |
| lat: [S.right] | -0.44 | 0.15 | 4368.23 | -2.89 | 0.004 | ** |
| sag: [S.posterior] | -0.02 | 0.11 | 4384.13 | -0.21 | 0.833 | |
| PTA_z | 0.21 | 0.26 | 28.21 | 0.80 | 0.429 | |
| att_resid_z | 0.33 | 0.27 | 28.81 | 1.24 | 0.227 | |
| prestim:conditionORAI | 0.06 | 0.03 | 3970.95 | 2.22 | 0.026 | * |
| condition: ORAI:lat[S.medial] | -0.14 | 0.21 | 4367.65 | -0.64 | 0.522 | |
| condition: ORAI:lat[S.right] | 0.04 | 0.21 | 4368.56 | 0.18 | 0.861 | |
| condition: ORAI:sag[S.posterior] | -0.22 | 0.15 | 4384.03 | -1.44 | 0.151 | |
| lat[S.medial]:sag[S.posterior] | -0.02 | 0.15 | 4367.96 | -0.14 | 0.888 | |
| lat[S.right]:sag[S.posterior] | 0.00 | 0.15 | 4367.64 | 0.00 | 0.999 | |
| condition: ORAI:PTA_z | -0.02 | 0.26 | 26.66 | -0.08 | 0.941 | |
| lat[S.medial]:PTA_z | 0.08 | 0.15 | 4367.84 | 0.54 | 0.592 | |
| lat[S.right]:PTA_z | -0.16 | 0.15 | 4367.89 | -1.13 | 0.257 | |
| sag[S.posterior]:PTA_z | 0.06 | 0.10 | 4367.72 | 0.57 | 0.569 | |
| condition: ORAI:lat[S.medial]:sag[S.posterior] | -0.01 | 0.21 | 4367.99 | -0.04 | 0.965 | |
| condition: ORAI:lat[S.right]:sag[S.posterior] | 0.05 | 0.21 | 4367.67 | 0.22 | 0.825 | |
| condition: ORAI:lat[S.medial]:PTA_z | 0.02 | 0.21 | 4367.74 | 0.09 | 0.925 | |
| condition: ORAI:lat[S.right]:PTA_z | 0.18 | 0.21 | 4367.77 | 0.90 | 0.37 | |
| condition: ORAI:sag[S.posterior]:PTA_z | -0.09 | 0.15 | 4367.80 | -0.63 | 0.528 | |
| lat[S.medial]:sag[S.posterior]:PTA_z | 0.01 | 0.15 | 4367.64 | 0.04 | 0.971 | |
| lat[S.right]:sag[S.posterior]:PTA_z | 0.01 | 0.15 | 4367.64 | 0.07 | 0.946 | |
| condition: ORAI:lat[S.medial]:sag[S.posterior]:PTA_z | 0.04 | 0.21 | 4367.65 | 0.18 | 0.856 | |
| condition: ORAI:lat[S.right]:sag[S.posterior]:PTA_z | -0.06 | 0.21 | 4367.64 | -0.30 | 0.767 | |

Table S28: Summary of Linear Mixed Model for the N400 amplitude predicted by condition (ORAI vs. ORIA), laterality, sagittality, and RS

| | Estimate | Std. Error | df | t value | Pr(>\|t\|) | |
|---|---|---|---|---|---|---|
| (Intercept) | 0.74 | 0.34 | 53.37 | 2.18 | 0.034 | * |
| prestim | 0.28 | 0.02 | 4157.72 | 15.19 | <0.001 | *** |
| condition: ORAI | -0.20 | 0.38 | 57.56 | -0.53 | 0.597 | |
| lat: [S.medial] | 0.40 | 0.15 | 4367.31 | 2.68 | 0.007 | ** |
| lat: [S.right] | -0.43 | 0.15 | 4367.84 | -2.85 | 0.004 | ** |
| sag: [S.posterior] | -0.02 | 0.11 | 4383.93 | -0.21 | 0.835 | |
| RS_z | -0.32 | 0.27 | 28.86 | -1.19 | 0.245 | |
| prestim:conditionORAI | 0.06 | 0.03 | 3976.99 | 2.23 | 0.026 | * |
| condition: ORAI:lat[S.medial] | -0.14 | 0.21 | 4367.32 | -0.64 | 0.525 | |
| condition: ORAI:lat[S.right] | 0.03 | 0.21 | 4368.22 | 0.16 | 0.876 | |
| condition: ORAI:sag[S.posterior] | -0.22 | 0.15 | 4384.04 | -1.44 | 0.151 | |
| lat[S.medial]:sag[S.posterior] | -0.02 | 0.15 | 4367.62 | -0.15 | 0.878 | |
| lat[S.right]:sag[S.posterior] | 0.00 | 0.15 | 4367.31 | 0.01 | 0.994 | |
| condition: ORAI:RS_z | -0.35 | 0.26 | 27.42 | -1.36 | 0.184 | |
| lat[S.medial]:RS_z | -0.07 | 0.15 | 4367.31 | -0.49 | 0.626 | |
| lat[S.right]:RS_z | 0.03 | 0.15 | 4367.52 | 0.23 | 0.82 | |
| sag[S.posterior]:RS_z | 0.00 | 0.11 | 4367.94 | 0.01 | 0.989 | |
| condition: ORAI:lat[S.medial]:sag[S.posterior] | -0.01 | 0.21 | 4367.66 | -0.03 | 0.973 | |
| condition: ORAI:lat[S.right]:sag[S.posterior] | 0.05 | 0.21 | 4367.35 | 0.22 | 0.824 | |
| condition: ORAI:lat[S.medial]:RS_z | -0.14 | 0.21 | 4367.42 | -0.65 | 0.518 | |
| condition: ORAI:lat[S.right]:RS_z | 0.01 | 0.21 | 4367.42 | 0.05 | 0.962 | |
| condition: ORAI:sag[S.posterior]:RS_z | -0.10 | 0.15 | 4367.96 | -0.66 | 0.507 | |
| lat[S.medial]:sag[S.posterior]:RS_z | -0.03 | 0.15 | 4367.31 | -0.21 | 0.835 | |
| lat[S.right]:sag[S.posterior]:RS_z | 0.02 | 0.15 | 4367.31 | 0.15 | 0.883 | |
| condition: ORAI:lat[S.medial]:sag[S.posterior]:RS_z | 0.10 | 0.21 | 4367.32 | 0.46 | 0.644 | |
| condition: ORAI:lat[S.right]:sag[S.posterior]:RS_z | 0.02 | 0.21 | 4367.32 | 0.11 | 0.909 | |

Table S29: Summary of Linear Mixed Model for the N400 amplitude predicted by condition (ORAI vs. ORIA), laterality, sagittality, and OS

| | Estimate | Std. Error | df | t value | Pr(>\|t\|) | |
|---|---|---|---|---|---|---|
| (Intercept) | 0.78 | 0.33 | 53.71 | 2.36 | 0.022 | * |
| prestim | 0.28 | 0.02 | 4085.26 | 15.18 | <0.001 | *** |
| condition: ORAI | -0.20 | 0.39 | 56.98 | -0.52 | 0.607 | |
| lat: [S.medial] | 0.42 | 0.15 | 4366.69 | 2.77 | 0.006 | ** |
| lat: [S.right] | -0.44 | 0.15 | 4367.32 | -2.88 | 0.004 | ** |
| sag: [S.posterior] | -0.02 | 0.11 | 4383.71 | -0.14 | 0.887 | |
| OS_z | -0.51 | 0.28 | 28.66 | -1.85 | 0.074 | . |
| prestim:conditionORAI | 0.06 | 0.03 | 3982.39 | 2.18 | 0.029 | * |
| condition: ORAI:lat[S.medial] | -0.13 | 0.21 | 4366.69 | -0.60 | 0.547 | |
| condition: ORAI:lat[S.right] | 0.05 | 0.21 | 4367.64 | 0.24 | 0.811 | |
| condition: ORAI:sag[S.posterior] | -0.21 | 0.15 | 4382.87 | -1.35 | 0.178 | |
| lat[S.medial]:sag[S.posterior] | -0.02 | 0.15 | 4367.02 | -0.12 | 0.902 | |
| lat[S.right]:sag[S.posterior] | 0.00 | 0.15 | 4366.69 | 0.03 | 0.979 | |
| condition: ORAI:OS_z | 0.07 | 0.28 | 27.52 | 0.25 | 0.804 | |
| lat[S.medial]:OS_z | -0.17 | 0.16 | 4366.69 | -1.07 | 0.287 | |
| lat[S.right]:OS_z | 0.05 | 0.16 | 4366.75 | 0.31 | 0.759 | |
| sag[S.posterior]:OS_z | -0.14 | 0.11 | 4367.91 | -1.24 | 0.217 | |
| condition: ORAI:lat[S.medial]:sag[S.posterior] | -0.02 | 0.21 | 4367.06 | -0.08 | 0.938 | |
| condition: ORAI:lat[S.right]:sag[S.posterior] | 0.04 | 0.21 | 4366.72 | 0.18 | 0.854 | |
| condition: ORAI:lat[S.medial]:OS_z | -0.04 | 0.22 | 4366.76 | -0.17 | 0.869 | |
| condition: ORAI:lat[S.right]:OS_z | -0.21 | 0.22 | 4366.74 | -0.95 | 0.345 | |
| condition: ORAI:sag[S.posterior]:OS_z | -0.11 | 0.16 | 4368.04 | -0.70 | 0.483 | |
| lat[S.medial]:sag[S.posterior]:OS_z | -0.04 | 0.16 | 4366.69 | -0.28 | 0.777 | |
| lat[S.right]:sag[S.posterior]:OS_z | -0.06 | 0.16 | 4366.71 | -0.40 | 0.69 | |
| condition: ORAI:lat[S.medial]:sag[S.posterior]:OS_z | 0.09 | 0.22 | 4366.72 | 0.42 | 0.675 | |
| condition: ORAI:lat[S.right]:sag[S.posterior]:OS_z | 0.11 | 0.22 | 4366.70 | 0.51 | 0.61 | |

Table S30: Summary of Linear Mixed Model for the N400 amplitude predicted by condition (ORAI vs. ORIA), laterality, sagittality, and IAF

| | Estimate | Std. Error | df | t value | Pr(>\|t\|) | |
|---|---|---|---|---|---|---|
| (Intercept) | 0.75 | 0.34 | 52.77 | 2.19 | 0.033 | * |
| prestim | 0.28 | 0.02 | 4136.82 | 15.21 | <0.001 | *** |
| condition: ORAI | -0.22 | 0.39 | 57.41 | -0.55 | 0.584 | |
| lat: [S.medial] | 0.41 | 0.15 | 4367.53 | 2.69 | 0.007 | ** |
| lat: [S.right] | -0.43 | 0.15 | 4368.13 | -2.80 | 0.005 | ** |
| sag: [S.posterior] | -0.02 | 0.11 | 4383.84 | -0.18 | 0.859 | |
| IAF_z | -0.11 | 0.29 | 29.16 | -0.36 | 0.719 | |
| prestim:conditionORAI | 0.06 | 0.03 | 3987.73 | 2.20 | 0.028 | * |
| condition: ORAI:lat[S.medial] | -0.16 | 0.22 | 4367.54 | -0.72 | 0.469 | |
| condition: ORAI:lat[S.right] | 0.02 | 0.22 | 4368.44 | 0.10 | 0.924 | |
| condition: ORAI:sag[S.posterior] | -0.23 | 0.16 | 4383.56 | -1.50 | 0.135 | |
| lat[S.medial]:sag[S.posterior] | -0.01 | 0.15 | 4367.85 | -0.09 | 0.925 | |
| lat[S.right]:sag[S.posterior] | 0.01 | 0.15 | 4367.53 | 0.04 | 0.967 | |
| condition: ORAI:IAF_z | 0.16 | 0.28 | 28.35 | 0.56 | 0.58 | |
| lat[S.medial]:IAF_z | -0.01 | 0.16 | 4367.55 | -0.07 | 0.942 | |
| lat[S.right]:IAF_z | -0.05 | 0.16 | 4367.57 | -0.32 | 0.747 | |
| sag[S.posterior]:IAF_z | -0.03 | 0.11 | 4367.53 | -0.25 | 0.8 | |
| condition: ORAI:lat[S.medial]:sag[S.posterior] | -0.02 | 0.22 | 4367.89 | -0.07 | 0.942 | |
| condition: ORAI:lat[S.right]:sag[S.posterior] | 0.04 | 0.22 | 4367.56 | 0.20 | 0.839 | |
| condition: ORAI:lat[S.medial]:IAF_z | 0.17 | 0.22 | 4367.58 | 0.77 | 0.443 | |
| condition: ORAI:lat[S.right]:IAF_z | 0.12 | 0.22 | 4367.55 | 0.53 | 0.599 | |
| condition: ORAI:sag[S.posterior]:IAF_z | 0.10 | 0.16 | 4367.56 | 0.63 | 0.528 | |
| lat[S.medial]:sag[S.posterior]:IAF_z | -0.06 | 0.16 | 4367.53 | -0.38 | 0.704 | |
| lat[S.right]:sag[S.posterior]:IAF_z | -0.05 | 0.16 | 4367.57 | -0.33 | 0.741 | |
| condition: ORAI:lat[S.medial]:sag[S.posterior]:IAF_z | 0.05 | 0.22 | 4367.55 | 0.22 | 0.829 | |
| condition: ORAI:lat[S.right]:sag[S.posterior]:IAF_z | 0.03 | 0.22 | 4367.54 | 0.15 | 0.879 | |

Table S31: Summary of Linear Mixed Model for the N400 amplitude predicted by condition (ORAI vs. ORIA), laterality, sagittality, and Flanker

| | Estimate | Std. Error | df | t value | Pr(>\|t\|) | |
|---|---|---|---|---|---|---|
| (Intercept) | 0.73 | 0.34 | 53.19 | 2.14 | 0.037 | * |
| prestim | 0.28 | 0.02 | 4114.75 | 15.23 | <0.001 | *** |
| condition: ORAI | -0.19 | 0.39 | 57.26 | -0.49 | 0.628 | |
| lat: [S.medial] | 0.39 | 0.15 | 4367.31 | 2.60 | 0.009 | ** |
| lat: [S.right] | -0.40 | 0.15 | 4367.90 | -2.67 | 0.008 | ** |
| sag: [S.posterior] | -0.02 | 0.11 | 4384.08 | -0.17 | 0.862 | |
| flanker_z | -0.29 | 0.30 | 29.11 | -0.97 | 0.338 | |
| prestim:conditionORAI | 0.06 | 0.03 | 3968.18 | 2.18 | 0.029 | * |
| condition: ORAI:lat[S.medial] | -0.13 | 0.21 | 4367.31 | -0.59 | 0.557 | |
| condition: ORAI:lat[S.right] | 0.02 | 0.21 | 4368.22 | 0.10 | 0.922 | |
| condition: ORAI:sag[S.posterior] | -0.23 | 0.15 | 4383.61 | -1.49 | 0.138 | |
| lat[S.medial]:sag[S.posterior] | -0.01 | 0.15 | 4367.64 | -0.10 | 0.922 | |
| lat[S.right]:sag[S.posterior] | -0.00 | 0.15 | 4367.31 | -0.01 | 0.99 | |
| conditionORAI:flanker_z | 0.12 | 0.29 | 28.87 | 0.39 | 0.696 | |
| lat[S.medial]:flanker_z | -0.23 | 0.17 | 4367.41 | -1.40 | 0.162 | |
| lat[S.right]:flanker_z | 0.44 | 0.17 | 4367.32 | 2.67 | 0.008 | ** |
| sag[S.posterior]:flanker_z | 0.06 | 0.12 | 4367.31 | 0.50 | 0.619 | |
| condition: ORAI:lat[S.medial]:sag[S.posterior] | -0.01 | 0.21 | 4367.68 | -0.06 | 0.952 | |
| condition: ORAI:lat[S.right]:sag[S.posterior] | 0.05 | 0.21 | 4367.34 | 0.25 | 0.805 | |
| conditionORAI:lat[S.medial]:flanker_z | 0.19 | 0.24 | 4367.36 | 0.78 | 0.435 | |
| conditionORAI:lat[S.right]:flanker_z | -0.28 | 0.24 | 4367.31 | -1.18 | 0.237 | |
| conditionORAI:sag[S.posterior]:flanker_z | -0.11 | 0.17 | 4367.40 | -0.63 | 0.531 | |
| lat[S.medial]:sag[S.posterior]:flanker_z | 0.11 | 0.17 | 4367.36 | 0.64 | 0.524 | |
| lat[S.right]:sag[S.posterior]:flanker_z | -0.03 | 0.17 | 4367.31 | -0.18 | 0.858 | |
| conditionORAI:lat[S.medial]:sag[S.posterior]:flanker_z | -0.07 | 0.24 | 4367.36 | -0.29 | 0.769 | |
| conditionORAI:lat[S.right]:sag[S.posterior]:flanker_z | 0.06 | 0.24 | 4367.36 | 0.26 | 0.793 | |

Table S32: Summary of Linear Mixed Model for the N400 amplitude predicted by condition (ORAI vs. ORIA), laterality, sagittality, and Stroop

| | Estimate | Std. Error | df | t value | Pr(>\|t\|) | |
|---|---|---|---|---|---|---|
| (Intercept) | 0.73 | 0.34 | 52.86 | 2.13 | 0.038 | * |
| prestim | 0.28 | 0.02 | 4162.90 | 15.19 | <0.001 | *** |
| condition: ORAI | -0.20 | 0.39 | 57.36 | -0.52 | 0.603 | |
| lat: [S.medial] | 0.40 | 0.15 | 4367.35 | 2.62 | 0.009 | ** |
| lat: [S.right] | -0.42 | 0.15 | 4367.95 | -2.77 | 0.006 | ** |
| sag: [S.posterior] | -0.04 | 0.11 | 4383.90 | -0.36 | 0.719 | |
| stroop_z | 0.17 | 0.28 | 29.12 | 0.61 | 0.548 | |
| prestim:conditionORAI | 0.05 | 0.03 | 4066.12 | 2.16 | 0.03 | * |
| condition: ORAI:lat[S.medial] | -0.13 | 0.21 | 4367.36 | -0.62 | 0.538 | |
| condition: ORAI:lat[S.right] | 0.01 | 0.21 | 4368.26 | 0.05 | 0.962 | |
| condition: ORAI:sag[S.posterior] | -0.22 | 0.15 | 4383.45 | -1.39 | 0.163 | |
| lat[S.medial]:sag[S.posterior] | -0.02 | 0.15 | 4367.67 | -0.12 | 0.906 | |
| lat[S.right]:sag[S.posterior] | -0.00 | 0.15 | 4367.35 | -0.03 | 0.977 | |
| condition: ORAI:stroop_z | 0.11 | 0.27 | 28.09 | 0.41 | 0.684 | |
| lat[S.medial]:stroop_z | 0.15 | 0.15 | 4367.78 | 1.00 | 0.318 | |
| lat[S.right]:stroop_z | -0.18 | 0.15 | 4367.48 | -1.14 | 0.252 | |
| sag[S.posterior]:stroop_z | 0.21 | 0.11 | 4367.61 | 1.90 | 0.058 | . |
| condition: ORAI:lat[S.medial]:sag[S.posterior] | -0.01 | 0.21 | 4367.69 | -0.03 | 0.972 | |
| condition: ORAI:lat[S.right]:sag[S.posterior] | 0.05 | 0.21 | 4367.38 | 0.23 | 0.821 | |
| condition: ORAI:lat[S.medial]:stroop_z | -0.01 | 0.22 | 4367.72 | -0.03 | 0.972 | |
| condition: ORAI:lat[S.right]:stroop_z | 0.37 | 0.22 | 4367.51 | 1.68 | 0.093 | . |
| condition: ORAI:sag[S.posterior]:stroop_z | -0.01 | 0.15 | 4367.73 | -0.09 | 0.927 | |
| lat[S.medial]:sag[S.posterior]:stroop_z | -0.05 | 0.15 | 4367.36 | -0.30 | 0.766 | |
| lat[S.right]:sag[S.posterior]:stroop_z | 0.05 | 0.15 | 4367.48 | 0.36 | 0.72 | |
| condition: ORAI:lat[S.medial]:sag[S.posterior]:stroop_z | -0.07 | 0.22 | 4367.39 | -0.31 | 0.76 | |
| condition: ORAI:lat[S.right]:sag[S.posterior]:stroop_z | 0.02 | 0.22 | 4367.52 | 0.09 | 0.926 | |

Table S33: Summary of Cumulative Link Mixed Model for the acceptability rating of a sentence belonging to the ORAI conditions, predicted by N400 amplitude

| | Model 1 |
|---|---|
| n400_z | $-0.20^{***}$ |
| | (0.02) |
| 1—2 | $-0.35$ |
| | (0.34) |
| 2—3 | $1.77^{***}$ |
| | (0.34) |
| 3—4 | $3.40^{***}$ |
| | (0.34) |
| Log Likelihood | -16916.48 |
| AIC | 33844.96 |
| BIC | 33891.71 |
| Num. obs. | 17894 |
| Groups (participant) | 29 |
| Groups (codeItem) | 27 |
| Variance: participant: (Intercept) | 2.36 |
| Variance: codeItem: (Intercept) | 0.93 |

$^{***}p < 0.001$, $^{**}p < 0.01$, $^{*}p < 0.05$
