## Supplementary Figures for "Individual Differences in Peripheral Hearing and Cognition Reveal Sentence Processing Differences in Healthy Older Adults"

Figure S1: Effects plots of P600 amplitude of the models with a significant condition*VOI interaction. P600 amplitude is plotted in six clusters defined by laterality and sagittality.

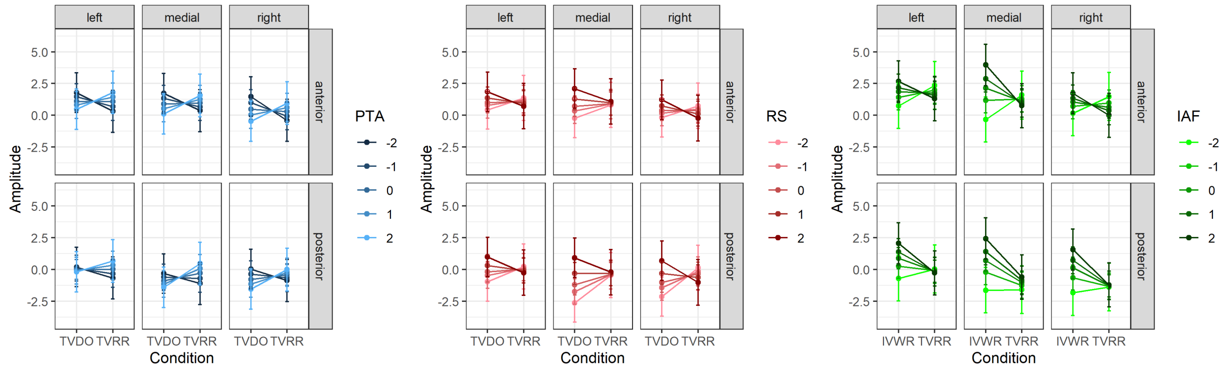

Figure S2: This figure shows the individual random slopes of condition by participant (left) and the individual random intercepts per condition (right) in the basic model for the N400 amplitude in the ORAI-ORIA comparison.

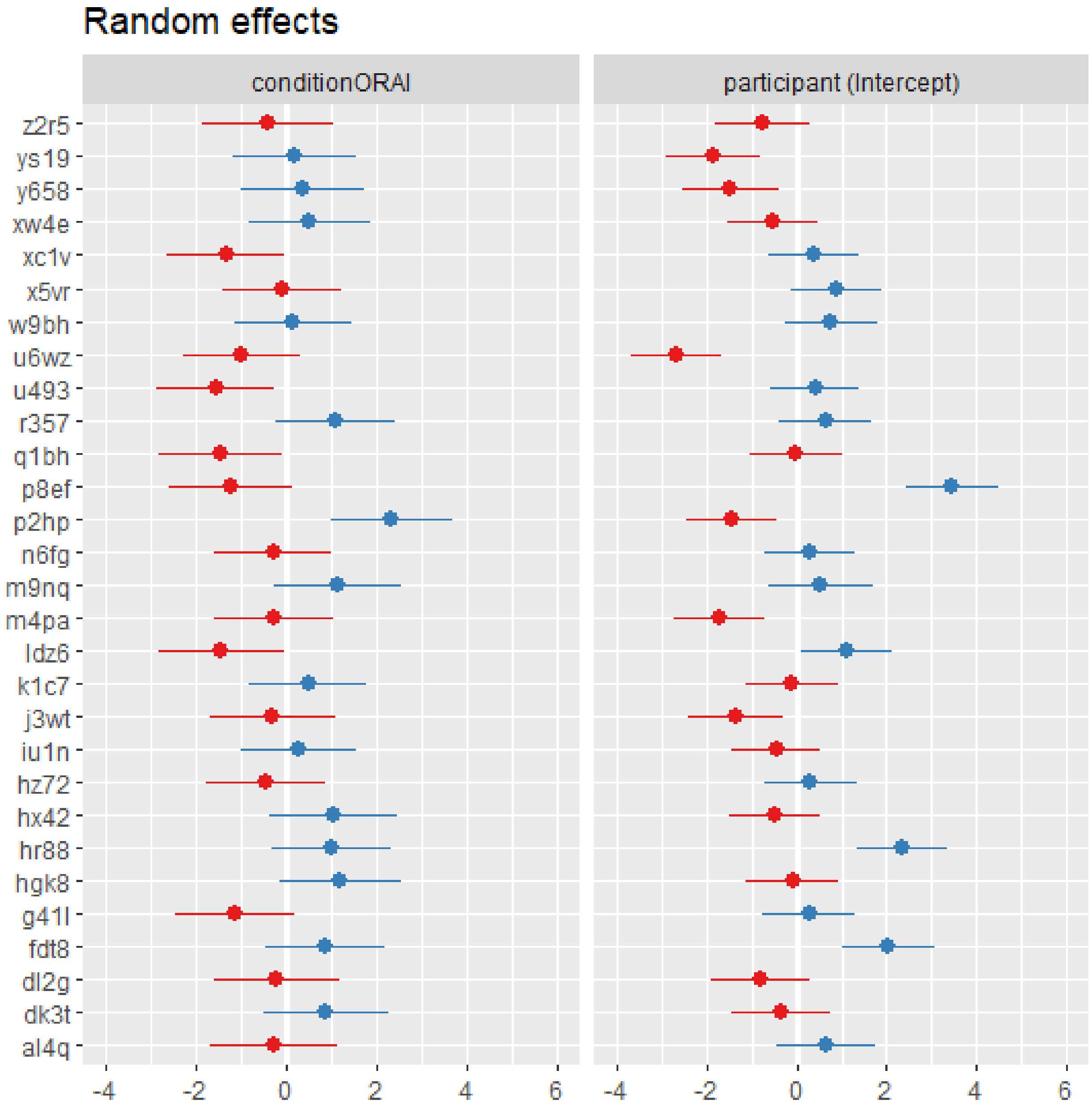

Figure S3: This figure shows the effect of n400_z on acceptability ratings, extracted from the model in Table S33.

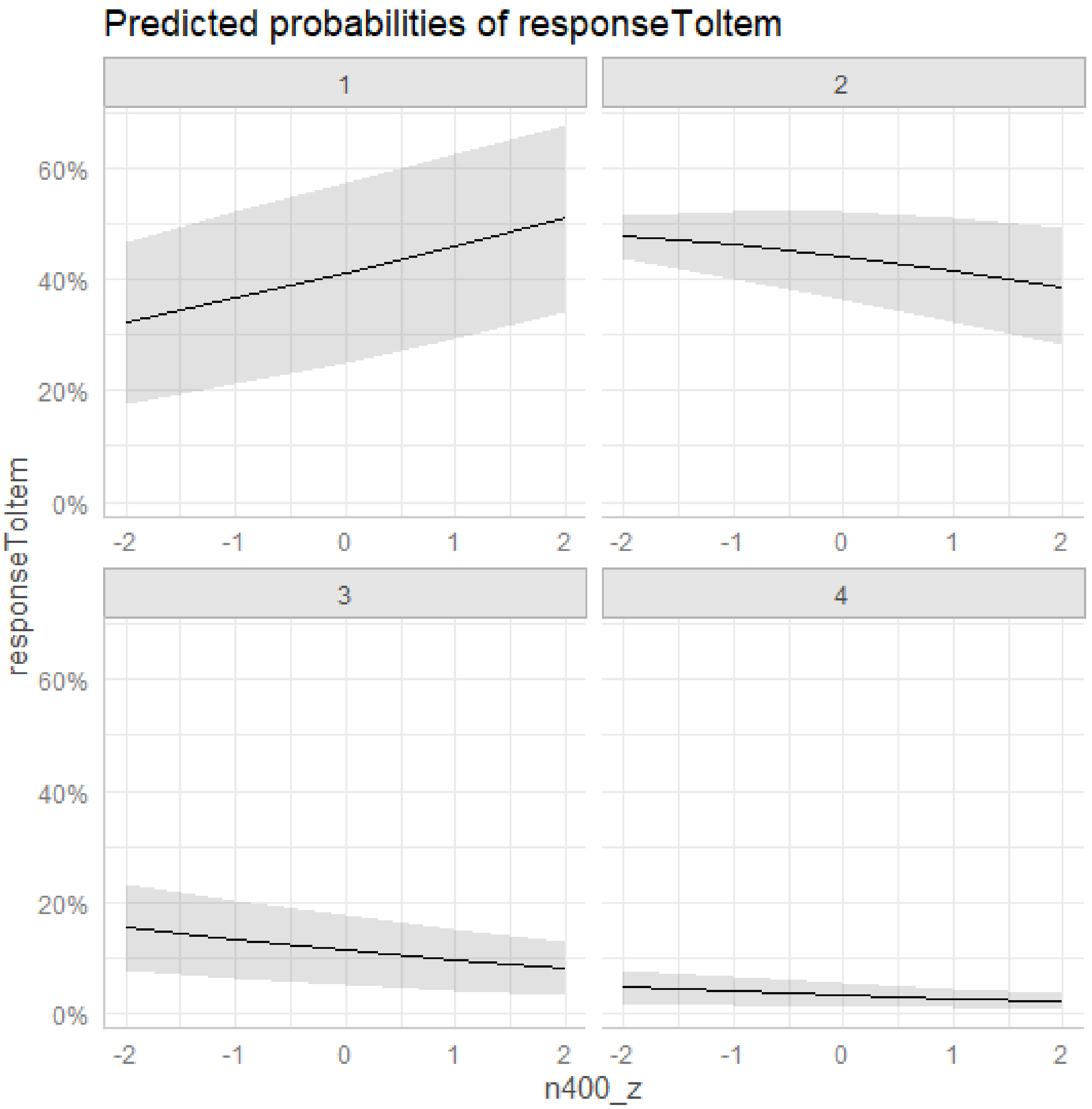
